# Dodecin as carrier protein for immunizations and bioengineering applications

**DOI:** 10.1101/2020.03.19.990861

**Authors:** Florian Bourdeaux, Yannick Kopp, Julia Lautenschläger, Ines Gößner, Hüseyin Besir, R. Martin Vabulas, Martin Grininger

## Abstract

In bioengineering, scaffold proteins have been increasingly used to recruit molecules to parts of a cell, or to enhance the efficacy of biosynthetic or signaling pathways. For example, scaffolds can be used to make weak or non-immunogenic small molecules immunogenic by attaching them to the scaffold, in this role called carrier. Here, we present the dodecin from *Mycobacterium tuberculosis* (*mt*Dod) as a new scaffold protein. *Mt*Dod is a homododecameric complex of spherical shape, high stability and robust assembly, which allows the attachment of cargo at its surface. We show that *mt*Dod, either directly loaded with cargo or equipped with domains for non-covalent and covalent loading of cargo, can be produced recombinantly in high quantity and quality in *Escherichia coli*. Fusions of *mt*Dod with proteins of up to four times the size of *mt*Dod, e.g. with monomeric superfolder green fluorescent protein creating a 437 kDa large dodecamer, were successfully purified, showing *mt*Dod’s ability to function as recruitment hub. Further, *mt*Dod equipped with SYNZIP and SpyCatcher domains for post-translational recruitment of cargo was prepared of which the *mt*Dod/SpyCatcher system proved to be particularly useful. In a case study, we finally show that *mt*Dod peptide fusions allow producing antibodies against human heat shock proteins and the C-terminus of heat shock cognate 70 interacting protein (CHIP).

**For Table of Contents Only:** 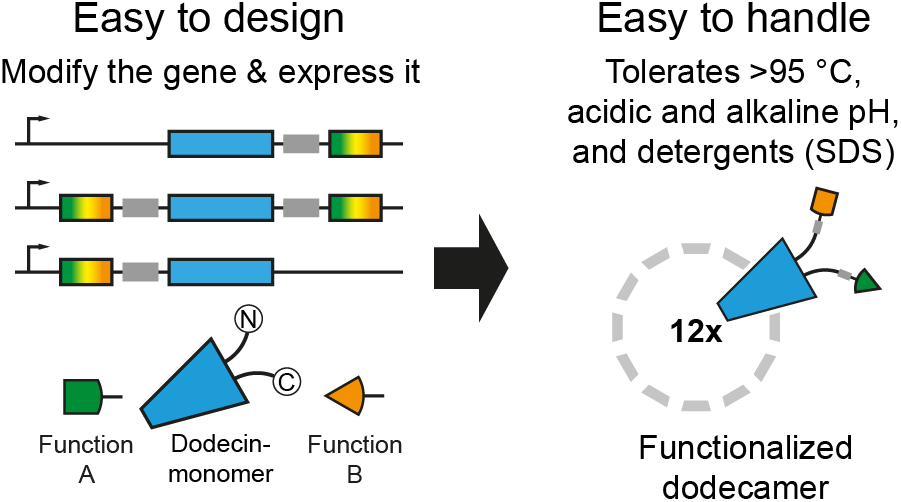

## Introduction

For being suited as carriers, proteins need to meet an array of requirements. They need to form a stable and water-soluble scaffold that is best insensitive to the attached cargo, and they should further allow the dense packing of the cargo in homovalent and ideally also in heterovalent fashion.^1–5^

One application of protein carriers is their conjugation with peptides for the generation of antibodies (AB) utilizing the increased immunogenicity of the carrier-peptide conjugate.^6^ Such ABs can identify proteins, which contain the peptides used for AB generation, in complex samples, and allow the specific labeling of proteins of interest in their spatiotemporal distribution; e.g. by immunofluorescence imaging or western blotting. As ABs preferentially recognize surface-exposed regions, termini of target proteins are usually a good source for exposed epitopes and their sequences are used as peptides for conjugation. In order to produce high-quality anti-peptide ABs, the peptide is ideally displayed at the carrier surface in similar orientation as in the intact protein. A dense packing of peptides at carrier surfaces is thought to be advantageous during immunization, because highly repetitive epitopes on particle surface facilitate B-cell activation through increased cell surface receptor oligomerization.^1,2^ In a typical process for forming peptide-carrier conjugates for AB production, an about 20 amino acid-long peptide is coupled to residues at the surface of a carrier protein by a chemical reaction.^7–9^ Commonly used carrier proteins are keyhole limpet hemocyanin (KLH), bovine serum albumin (BSA) and rabbit serum albumin (RSA), but also other proteins, e.g. tetanus toxoid (TT), and artificial carrier-systems, e.g. multiple antigen peptides (MAP) or virus-like particles (VLP), are used.^10,11^ While BSA bears typical carrier properties (likely also other albumins) and exposes the peptides at the surface at a potentially high density,^12,13^ KLH is often preferred as a carrier-protein due to its high immunogenicity.^14,15^ Notably, the immune system reacts to the entire conjugate, and, therefore, ABs are not just raised against the peptide of interest, but also against the carrier protein and the linker. Accordingly, it is important to use carrier-linker systems for immunization that are orthologous to the inventory of cells and tissues analyzed with the generated ABs.^15^ Heterovalent coating of carriers would allow for combining antigens to better represent complex pathogens as well as for co-coupling immunogenic sequences to broaden the immune response, as used in multi-epitope designs.^3–5^

The dodecin protein family was recently discovered as a flavin storage and buffering system that occurs in bacteria and archaea, but not in eukaryotes.^16–19^ Dodecins are 8 kDa small proteins of βαββ-topology. Although forming a small antiparallel β-sheet that partly enwraps the helix, the dodecin fold is unique. Dodecins largely meet the requirements of protein carriers. In the native dodecameric state, dodecins are of spherical shape with 23-cubic symmetry, and the N- and C-termini are exposed at the protein surface. Dodecins show pronounced thermostability (>95 °C)^18–20^, which likely originates from an extensive antiparallel β-sheet that is built upon protomer assembly.

Here, we present dodecin from *Mycobacterium tuberculosis* (*mt*Dod) as a new carrier protein for peptides and scaffold for bioengineering applications. Two approaches for charging *mt*Dod with peptide/protein antigens are presented: First, a cargo is directly fused by attaching the peptide/protein-encoding sequence at the gene level. Second, *mt*Dod is terminally modified with conjugation sites that allow post-translational covalent and non-covalent fusions of the peptide/protein as well as other chemical entities to the intact dodecin carrier (Fig. 1). *Mt*Dod constructs can be produced recombinantly in quantities of up to several 100 mg per liter of *Escherichia coli* culture. Purification from untreated cytosolic fractions is possible by first precipitating *E. coli* proteins at elevated temperatures followed by the precipitation of *mt*Dod constructs with organic solvents. *Mt*Dod constructs containing thermosensitive proteins or domains were successfully purified by affinity chromatography. As a case study, we demonstrate the suitability of *mt*Dod as carrier for producing anti-peptide ABs for laboratory use. ABs raised in rabbits against *mt*Dod-peptide fusions reached equal or surpassed quality compared to commercially available ABs as judged by western blotting.

**Fig. 1:**
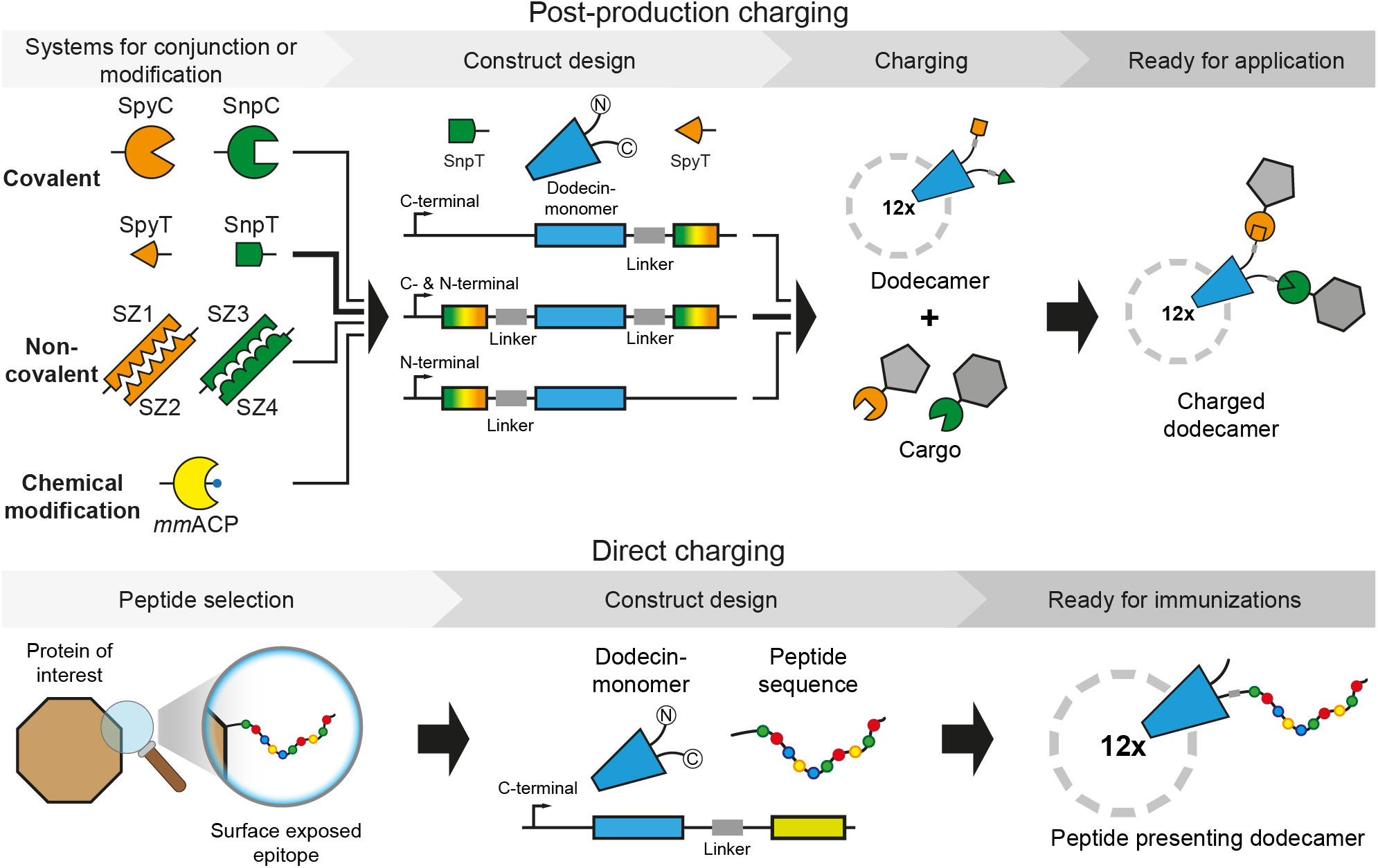
Schematic depiction of *mt*Dod constructs and workflow presented in this study. In this study, termini were modified directly at gene level with peptides, domains or proteins for subsequent charging with cargo (top) or direct coupling of peptides for immunizations (bottom). The dodecamers expose their termini at the outer surface. (Top) Selected example is highlighted by arrows in bold. SpyT and SpyC: SypTag and SpyCatcher.^**21,22**^ SnpT and SnpC: SnoopTag and SnoopCatcher.^**23**^ SZ1-SZ4: helical domains that bind to their specific counterpart, called SYNZIP.^**24**^ *Mm*ACP: *Mus musculus* acyl carrier protein (ACP), gene *Fasn*.

## Results and Discussion

### Dodecin can be recombinantly produced in high yields

To evaluate the suitability of *mt*Dod as a carrier protein, several *mt*Dod constructs were designed and purified. All constructs were expressed in *E. coli* BL21(DE3). Cells were grown in terrific broth (TB) medium to an optical density at 600 nm (OD_600_) of about 0.6-0.8 at 37 °C before induction with isopropyl-β-D-thiogalactopyranosid (IPTG; 0.5 mM final concentration), and expression was performed over night at 20 °C. Since *mt*Dod is a flavin binding protein (preferred flavin-ligand is riboflavin-5’-phosphate (FMN)),^19,20^ its overexpression causes increased amounts of cellular flavin, leading to a yellowish coloring of the cells. Cells were lysed by French press, and the cell debris was removed by centrifugation. Most *mt*Dod constructs were produced as soluble proteins, but some proteins, such as *mt*Dod-SZ1, *mt*Dod-SZ3 (SYNZIP constructs)^24^, H8-SpyC-*mt*Dod and *mt*Dod-SpyC-H8 (SpyC constructs)^21,22^ (nomenclature: N-terminal component named first and C-terminal component last) accumulated as inclusion bodies (Table 1).

**Table 1:** Selection of *mt*Dod constructs used for expression studies and divided into two groups: *mt*Dod-peptides (constructs with only short peptides fused to *mt*Dod) and *mt*Dod-proteins (constructs with domains or entire proteins fused to *mt*Dod). *mt*Dod(WT): wild type *mt*Dod. *se*ACP: *Saccharopolyspora erythraea* ACP, gene *chlB2*. msfGFP: monomeric superfolder green fluorescent protein.^**25**^ For a full description of constructs, see Table S1.

| Construct name | Linker system | Molar mass /Da | Expression state<sup>#</sup> |
| --- | --- | --- | --- |
| ***mtDod*-peptides** |  |  |  |
| *mtDod*(WT) | - | 7497 | soluble |
| *mtDod*-GSG-Lys | GSG | 8411 | soluble |
| *mtDod*-PAS-Met | PAS | 8876 | soluble |
| *mtDod*-SpyT | PASG | 10458 | soluble |
| SpyT-*mtDod* | GPAS | 10215 | soluble |
| *mtDod*-PAS2-SpyT | PAS2G | 11447 | soluble |
| SpyT-PAS2-*mtDod* | GPAS2 | 11205 | soluble |
| SpyT-*mtDod*-SnpT | GPAS / PASG | 13142 | soluble |
| ***mtDod*-proteins** |  |  |  |
| *mtDod*-mmACP | PAS | 19334 | soluble |
| *mtDod*-msfGFP-H8 | PAS | 36388 | soluble |
| *mtDod*-SpyC-H8* | PAS | 22413 | inclusion body |
| H8-SpyC-*mtDod* | PAS | 22072 | inclusion body |
| *mtDod*-SZ1** | PAS | 14232 | inclusion body |
| *mtDod*-SZ3** | PAS | 13396 | inclusion body |
| *mtDod*-seACP*** | PAS | 19966 | inclusion body |
| Linker details |  |  |  |
| GSG: | GGGGSGGGG | PAS: | SPAAPAPASPAS |
| PASG: | SPAAPAPASPASGGSG | GPAS: | GGSGSPAAPAPASPAS |
| PAS2G: | SPAAPAPASPASPAPSAPAASPAA<br>GGSG | GPAS2: | GGSGSPAAPAPASPASPAPSAPAA<br>SPAA |

<sup>#</sup> Describes whether the major fraction of the expressed construct is soluble or forms inclusion bodies. *MtDod* constructs with low solubility form often non-classical inclusion bodies (correctly folded protein),<sup>26</sup> leading to their yellowish coloring (flavin binding). Since flavin binding only requires intact *mtDod*, it is possible that inclusion bodies are yellow although the protein cargo is misfolded.
\* *MtDod*-SpyC-H8 seems to be soluble in cellular environment, but forms yellow aggregates after cell lysis.
\*\* SZ1-*mtDod* and SZ3-*mtDod* also formed inclusion bodies and behaved similarly as the C-terminal constructs (data not shown).
\*\*\* *MtDod*-seACP could not be obtained in soluble form; under all refolding conditions yellow aggregate was formed.

For soluble *mt*Dod constructs, most cytosolic *E. coli* proteins were removed by heat denaturation at about 75 °C. *Mt*Dod itself is stable to temperatures above 95 °C under standard conditions (pH ~7.5 and ionic strength >100 mM, e.g., in PBS), and the thermal stability can be further increased by adding the native FMN ligand in excess.^19,20^ Depending on the stability of the fused cargo, lower temperatures during the heat denaturation may be necessary, or different purification approaches need to be applied (affinity chromatography). For example, *mt*Dod-*mm*ACP started to precipitate at about 55-60 °C in spite of *mt*Dod staying intact, as indicated by maintained FMN binding and preserved dodecameric stability (Fig. S1). In this case, heat denaturation was conducted at about 55 °C. Lower temperatures during the heat denaturation step can affect the purity of preparations, because some *E. coli* proteins remain soluble. Following heat treatment, *mt*Dod constructs were generally further purified by two cycles of DMSO-induced precipitation (50% final DMSO concentration). Finally, size-exclusion chromatography (SEC) was performed to select for dodecameric fractions, which can be easily identified by the absorption bands of bound flavin (375 nm and 450 nm). For *mt*Dod-peptide fusions, the dodecamer turned out to be the main oligomeric species. While only minor peaks representing lower oligomeric states were detected, aggregation peaks were observed under high concentrations (Fig. S2). For larger *mt*Dod constructs with fused proteins, like *mt*Dod-*mm*ACP, significant aggregation was observed in the SEC profiles (see Fig. S2).

For the construct *mt*Dod-msfGFP-H8, purification by heat denaturation (70 °C, above that aggregation was observed) and purification by affinity chromatography were compared. GFP is a suitable cargo for this test, because GFP is highly thermostable.^27^ The dodecameric structure of dodecin causes a high density of surface exposed affinity tags allowing vigorous washing without severe protein loss during Ni-chelating affinity chromatography. Accordingly, *mt*Dod-msfGFP-H8 was washed with two column volumes of a 200 mM imidazole-containing wash buffer, while elution was performed at 400 mM imidazole. While with both purification strategies *mt*Dod-msfGFP-H8 dodecamer was obtained, the sample purified by heat denaturation showed severe aggregation in SEC (Fig. S3).

*Mt*Dod constructs that aggregate in inclusion bodies can be refolded by dialysis, as previously described ^19^, under conditions optimized for the respective fused cargo. All inclusion bodies were first washed and then dissolved by denaturation using 6 M guanidinium chloride. *Mt*Dod was refolded without further purification at different conditions ranging from pH 5.0^19^ to pH 8.5. Refolding was possible for all constructs obtained as inclusion bodies in this study, although the resolubilized proteins remained aggregation-prone, particularly during protein concentration and filtration. For a screen of buffer conditions for refolding constructs *mt*Dod-SpyC-H8 and H8-SpyC-*mt*Dod, see Fig. S4. Notably, for both constructs, a glycerol-containing buffer was found to be best suited for refolding.

Overall, all constructs presented in Table 1, except *mt*Dod-*se*ACP, were obtained in high purity (Fig. 2).

**Fig. 2:**
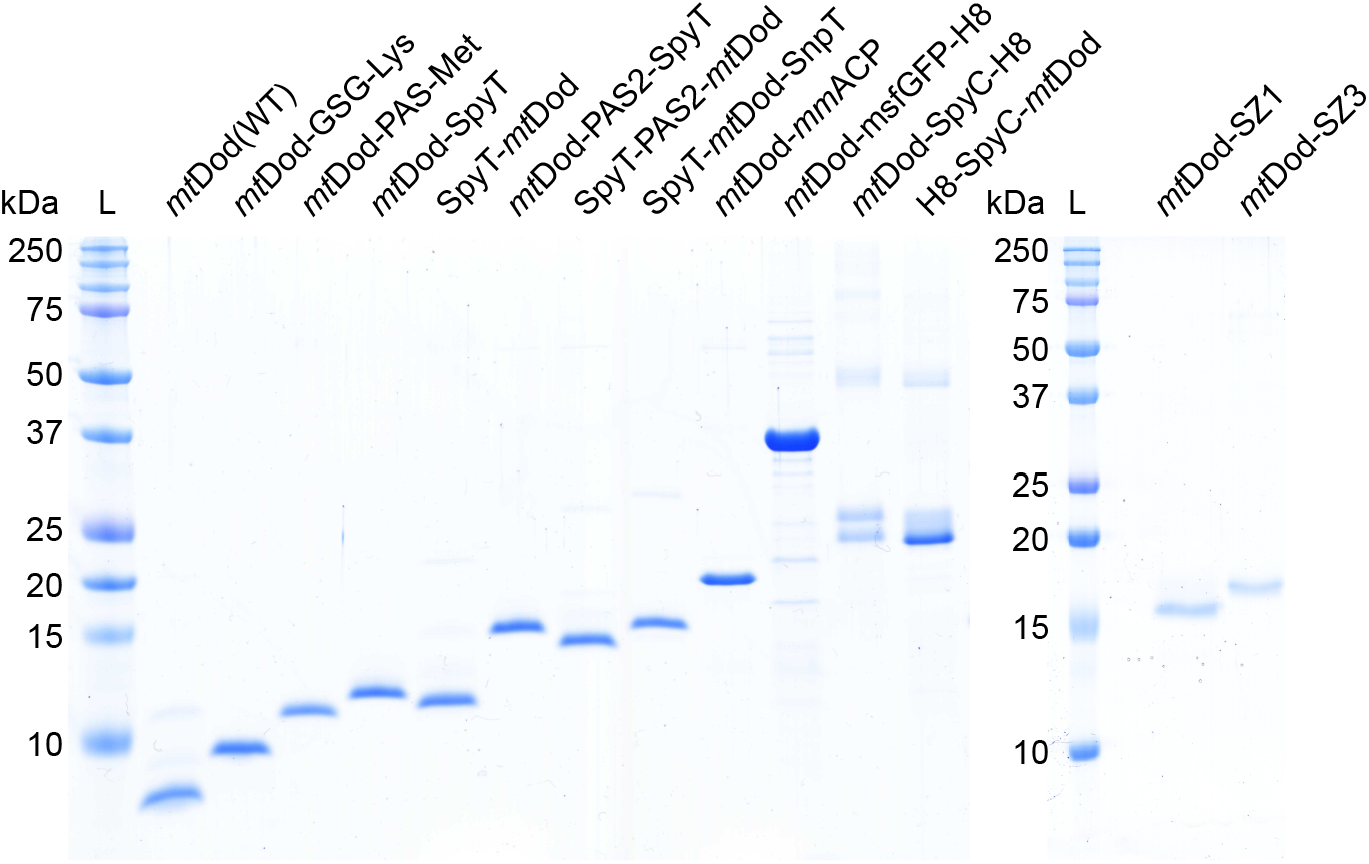
SDS-PAGE gel of purified *mt*Dod constructs. L: Ladder. For full dissociation of the dodecamers in SDS-PAGE, an acidic loading buffer containing 3.3% SDS was used during the heat treatment (5 min 60 °C). After the heat treatment, the pH was increased to about 6.8 using a glycerol- and Tris-HCl-containing buffer, followed by a second heat treatment (5 min 95 °C). *Mt*Dod-SZ1 and *mt*Dod-SZ2 were denatured with loading buffer containing ~7 M urea and 2.5% SDS (prolonged heat treatment: 15 min 95 °C). For most constructs, the denaturation with acidic loading buffer was more reliable and practical compared to urea-based protocols (in some cases even 7-8 M urea failed to dissociate the protein completely). Of note, when treated with acidic loading buffer, some constructs showed additional bands (mainly SpyC constructs, Fig. S5). The origin of this behavior was not further investigated.

We thought that the insolubility and aggregation problems observed for some constructs may be solved by the formation of *mt*Dod-heterododecamers, because then the density of entities on the surface could be reduced. To probe heterododecamer formation with *mt*Dod in vitro and in vivo, we worked with the two species *mt*Dod-PAS-Strep and *mt*Dod(WT). For in vitro heterododecamer formation, *mt*Dod(WT) and *mt*Dod-PAS-Strep were jointly refolded in different relative concentrations, while for the formation of heterododecamers in vivo, three combinations of the *mt*Dod constructs were expressed polycistronically. The analysis of heterododecamer compositions was possible by the high stability of the dodecamers (see Fig. S5) and the different migration behavior of constructs in SDS-PAGE (Fig. 3). Data indicates that the composition of heterododecamers is controlled by the relative concentration of species in the refolding solution; i.e., the higher the concentration of a construct, the more abundant it is in the refolded dodecamer. Of note, assuming that the heterododecamer formation is just controlled by the concentration of each construct (see Fig. 3 a), the band patterns for heterododecamers assembled in vivo can be used to estimate gene order related expression strength (see Fig. 3 b), as described in the literature for other methods, e.g. FRET.^28^ The estimated relative expression strength of each gene is for the bicistronic vector: first > second, and for the tricistronic vector first > third > second.

**Fig. 3:**
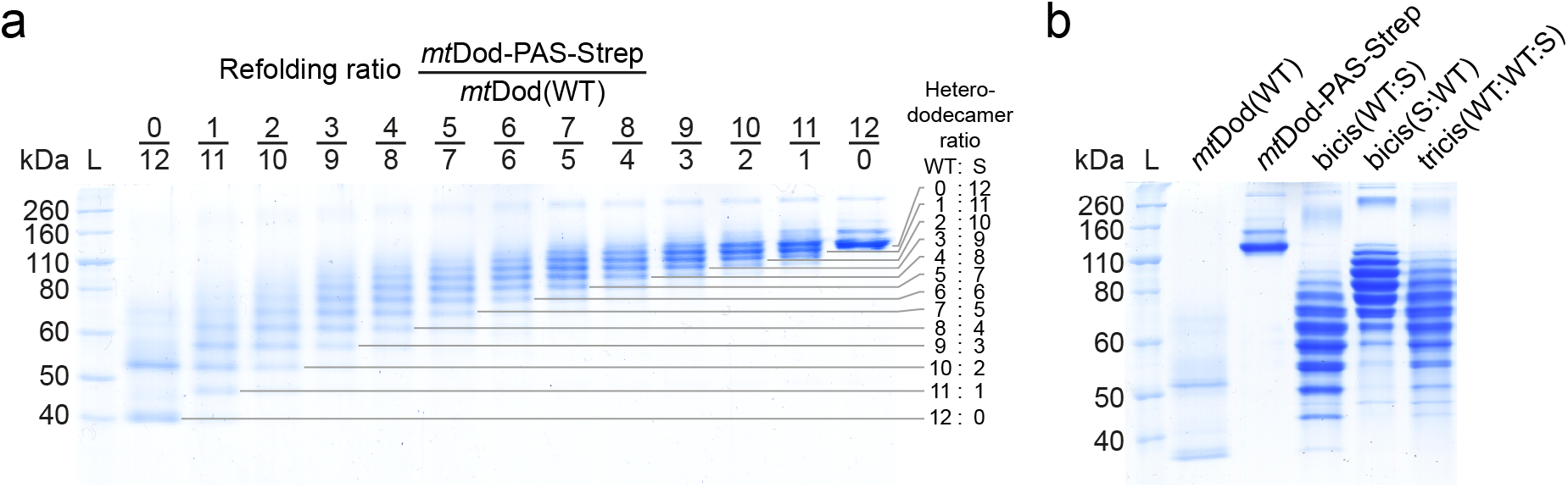
SDS-PAGE gel of purified heterododecamers of *mt*Dod(WT) (WT) and *mt*Dod-PAS-Strep (S). For analysis of heterododecamer composition, the tris-glycine (Lämmli) SDS-PAGE system was used, and samples were prepared without heat treatment and SDS. L: Ladder. a) SDS-PAGE gel of heterododecamers obtained by refolding *mt*Dod(WT) and *mt*Dod-PAS-Strep at different ratios. Each band represents a heterododecamer with a defined composition of *mt*Dod-PAS-Strep and *mt*Dod(WT) (13 bands for 13 possible compositions, higher artifact bands excluded). For *mt*Dod(WT) two bands are observable (both below the calculated mass of 90 kDa), the lower band is likely the dodecamer and the upper band an artifact. Such higher bands are observable for all constructs to some degree; the origin of this behavior was not further investigated. The origin of the exceptional migration properties (lower molecular weight) of *mt*Dod(WT) is not clear. b) SDS-PAGE gel of purified heterododecamers formed during polycistronic expression of *mt*Dod(WT) and *mt*Dod-PAS-Strep. Bicis(WT:S): Bicistronic expression vector design: *mt*Dod(WT) encoding gene first and *mt*Dod-PAS-Strep encoding gene second. Bicis(S:WT): Bicistronic expression vector design: *mt*Dod-PAS-Strep encoding gene first and *mt*Dod(WT) encoding gene second. Tricis(WT:WT:S): Tricistronic expression vector design: *mt*Dod(WT) encoding gene first and second and *mt*Dod-PAS-Strep encoding gene third.

### Dodecin is highly stable

We have recently established the cyclic thermal shift assay, termed thermocyclic fluorescence assay, to determine the stability of dodecins.^19^ This assay is based on the fluorescence quenching that is observed when flavins bind to dodecin. In each binding pocket of the dodecamer, the two isoalloxazine ring systems of two bound flavins are embedded between symmetry-related tryptophans.^18,19,29^ Since dodecins can only bind flavins in the dodecameric state, the fluorescence intensity of flavins can be used to estimate the amount of dodecameric *mt*Dod in solution. In contrast to standard melting analysis, in which the temperature is continuously increased, the thermocyclic fluorescence assay runs cyclic temperature profiles that contain a heating phase (temperature increased per cycle) and a cooling phase (for all cycles 5 °C). At the heating phase, FMN is released from the binding pocket and the fluorescence intensity increases. During cooling, FMN can rebind to the dodecamer (cooling phase) restoring initial low fluorescence values. As soon as the dodecamer denatures irreversibly, the fluorescence intensity remains at elevated levels. By plotting the fluorescence intensity of the cooling phase against the heating phase temperature, the thermal stability of the dodecamer of the *mt*Dod constructs can be determined.

Since all constructs, except *mt*Dod-SZ1 and *mt*Dod-SpyC-H8, proved to be stable in PBS buffer throughout the entire temperature range, we identified the slightly destabilizing conditions of pH 4.2 as suited to sense the impact of the cargo on the integrity of the *mt*Dod dodecameric scaffold (see Fig. 4). Under this condition, the thermally stable constructs *mt*Dod(WT) and *mt*Dod-peptides denatured at 75-80 °C. It is important to note that the thermocyclic fluorescence assay does only monitor the dodecameric stability, which may be influenced by the attached cargo. For example, in screening temperatures for the heat denaturation of *mt*Dod-*mm* ACP, we observed the formation of yellowish agglomerates above 55-60 °C, indicating that the construct is intact in the *mt*Dod scaffold, as capable of FMN binding, but precipitated by the thermally unfolded *mm* ACP (Fig. S6). The high dodecameric stability of *mt*Dod is also observed in SDS-PAGE using the standard loading buffer (2.5% SDS, pH 6.8) (see Fig. S5). Under these conditions, further depending on the heat treatment for sample preparation, a dodecameric fraction remains intact, as indicated by the high molecular weight band representing the dodecamer. In accordance to the lower stability at pH 4.2, observed in the thermocyclic fluorescence assay, a two-component acidic loading buffer (3.3% SDS and pH < 4.2 during heat treatment, afterwards 2.5% SDS and pH 6.8) was applied to fully denature the dodecamer (Fig. 2).

**Fig. 4:**
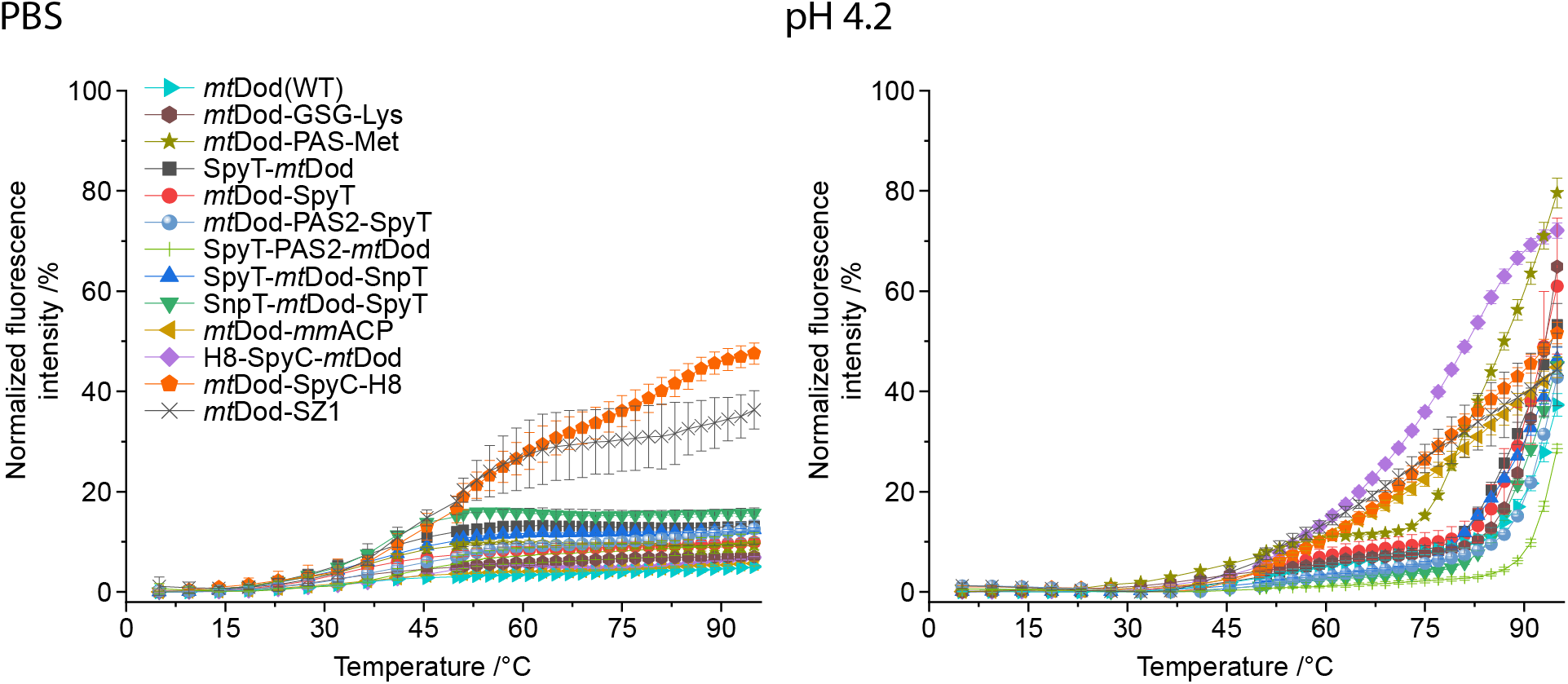
Thermal stablity of *mt*Dod constructs. The FMN fluorescence at the rebinding/cooling phase is plotted against the heating phase temperature. The increase of FMN fluorescence indicates disassembly of the dodecamer at the heating phase, as the flavin cannot rebind in the cooling phase and its fluorescence is not quenched. In PBS, only a negligible increase of fluorescene is observed within the entire temperature range, except for *mt*Dod-SpyC and *mt*Dod-SZ1, indicating that the dodecameric *mt*Dod core structures do not disassemble. The minor increase of fluorescence of up to 20% around 45-50 °C might be caused by hindered rebinding of FMN and not by disassembling of the dodecamer. At pH 4.2, for all constructs a steep (compared to PBS data) increase of fluorescence is observable indicating the dodecamer disassambly. Most constructs behave like *mt*Dod(WT) and are stable to about 80 °C, except *mt*Dod-PAS-Met, *mt*Dod-*mm* ACP, *mt*Dod-SpyC-H8 and H8-SpyC-*mt*Dod, of which the latter three start denaturing already at 50 °C. *mt*Dod-PAS-Met is only slightly less stable, and starts to denature around 75 °C. Note that *mt*Dod-SpyC-H8 and *mt*Dod-SZ1 suffer from strongly impaired FMN binding leading to non-saturated binding sites, and data needs to be treated with care. The FMN binding may also be altered by denaturing/aggregation of the fused fold, causing different curve profiles, e.g. *mt*Dod-*mm* ACP. For all measurements, fluorescence was normalized to the maximum values recorded in the heating phase, corrected by the temperature-induced fluorescence decline of FMN.

We observed that most *mt*Dod constructs can be frozen and thawed several times without noticeable aggregation. Of note, a glycerol-containing buffer was used for the SpyC constructs, and *mt*Dod-msfGFP-H8 showed minor formation of green fluorescent aggregates. *Mt*Dod SYNZIP constructs, while not showing severe lower stability, had solubility problems and formed yellow precipitate after thawing.

Overall, this data shows that *mt*Dod is an ideal carrier protein in its properties to be stable at high temperatures and during long-term storage. *Mt*Dod is tolerant towards the use of a wide range of conditions, and will accept a variety of protocols to attach cargos.

### Peptides/Proteins fused to *mt*Dod remain functional

The accessibility and functionality of folds and peptides fused to *mt*Dod were tested by the reactivity of the SpyT/-C and SnpT/-C pairs.^21–23^ These systems allow the covalent conjugation between two entities of which one is equipped with a peptide tag (Tag) and the other with a small protein fold (Catcher).^22^ Applications range from attaching proteins from pathogens to scaffolds, like VLPs and IMX313 (heptamer forming coiled coils), for immunizations,^30,31^ to recruiting enzymes to a scaffold hub for creating assemblies with elevated substrate turnover.^32^ In this study, *se*ACP-SpyC and mClover3-SnpC were prepared as cargo for performing SpyT/-C and SnpT/-C reactions with the respective Tag-labeled *mt*Dod constructs. For the inverse reaction, *mt*Dod SpyC constructs and SpyT-*se*ACP were used. For all reactions, the scaffold was saturated with two molar equivalents of cargo. The reactions were incubated for 20 h at 22 °C, and analyzed by SDS-PAGE (Fig. 5).

**Fig. 5:**
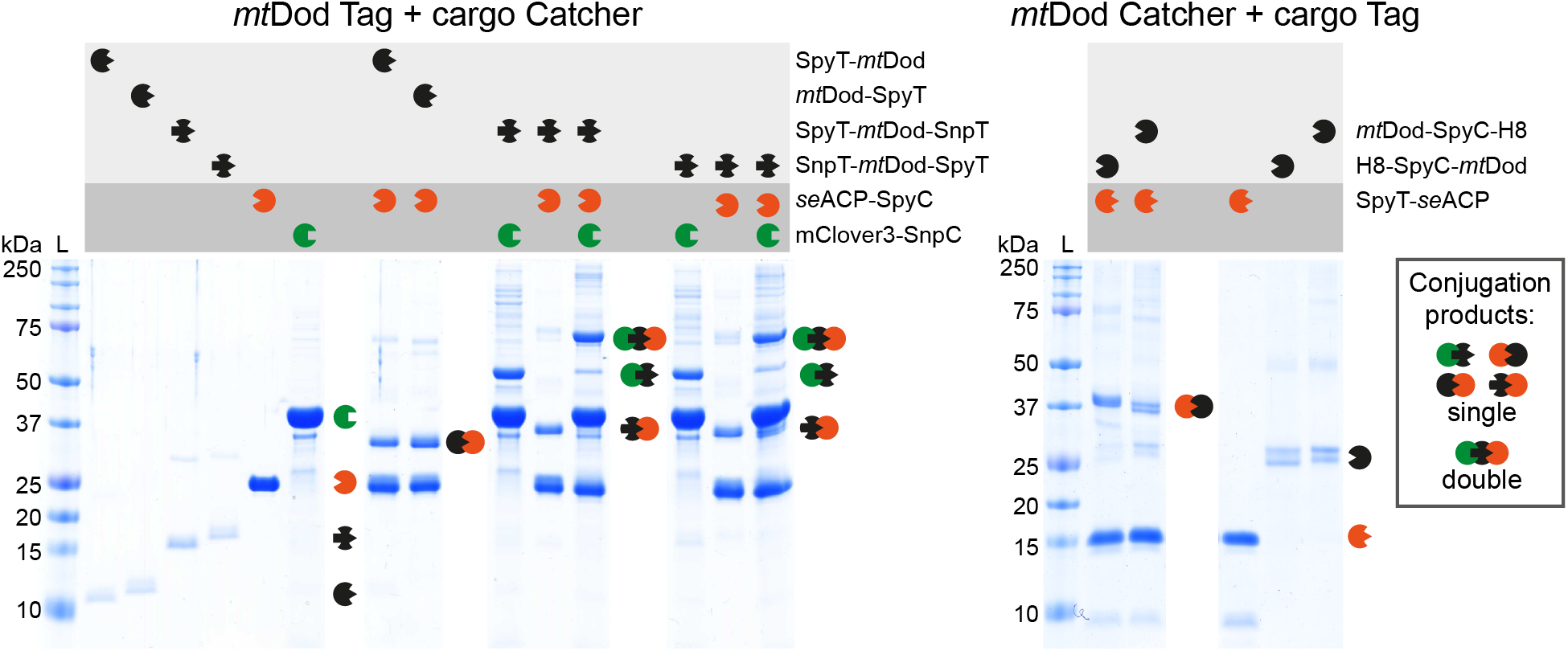
SDS-PAGE of the SpyT/-C and SnpT/-C reactions. Left: Reactions of *m*tDod Spy-/SnpT constructs with *se*ACP-SpyC and/or mClover3-SnpC. Right: Inverse reactions of *m*tDod SpyC constructs with SpyT-*se*ACP. In all reactions, bands of higher mass representing the conjugation products are observed. As mentioned above, the acidic loading buffer causes the appearance of double bands (*mt*Dod SpyC constructs) and smearing bands (*se*ACP-SpyC-H8) for some constructs.

For all combinations of *mt*Dod scaffold and cargo, the expected product band(s) of *mt*Dod and the specific cargo(s) were observed in SDS-PAGE. While for *mt*Dod SpyT/SnpT constructs no unreacted scaffold proteins was observed, for the inverse setting with *mt*Dod SpyC constructs, bands of unreacted scaffold monomer were visible (possibly caused by aggregation problems of the *mt*Dod SpyC constructs). We note that *mt*Dod SpyT/SnpT constructs are lower in molecular mass than the *mt*Dod SpyC constructs, and traces of unreacted scaffold protein may be less visible on SDS-PAGE gels. This data shows that a high degree of saturation was achieved, indicating that SpyT/-C and SnpT/-C are well accessible at the *mt*Dod dodecamer scaffold. Double-tagged constructs SpyT-*mt*Dod-SnpT or SnpT-*mt*Dod-SpyT, heterovalently loaded with *se*ACP-SpyC and mClover3-SnpC, revealed bands of single-charged *mt*Dod monomers in SDS-PAGE. We explain this observation by an increased density at the surface of *mt*Dod that sterically constrains the conjugation with both cargos. Similar as the SpyT/-C and SnpT/-C constructs also the SYNZIP constructs can be used for recruiting proteins to the *mt*Dod scaffold (although non-covalently). Due to the limited solubility and high aggregation tendencies of SYNZIP constructs, we only tested if *mt*Dod SYNZIP constructs are able to interact with the respective SYNZIP counterpart (e.g., *mt*Dod-SZ1 with SZ2-mClover3). For both *mt*Dod SYNZIP constructs, we observed the formation of *mt*Dod cargo adducts, indicated by higher apparent molecular mass peaks in SEC (Fig. S7). This shows that also SYNZIP domains fused to *mt*Dod are functional and accessible. However, we deemed the SpyT/-C and SnpT/-C systems more suitable for *mt*Dod constructs, and did not further investigated the SYNZIP system.

In order to probe the accessibility and functionality of linked folds further, we tested the labeling of *mm*ACP linked to *mt*Dod with a 4’-phosphopantetheine CoA fluorophore mediated by the 4’-phosphopantetheine transferase from *Bacillus subtilis* (Sfp). The Sfp-mediated modification of ACP with CoA-modified fluorophores (CoA-488; ATTO-TEC dye ATTO 488) has been frequently used for the labeling of cellular compounds.^33^ All reactions were conducted at 25 °C for 1 h in triplicates, and stopped by the addition of acidic loading buffer and analyzed by SDS-PAGE. To determine the relative accessibility of *mm*ACP linked to *mt*Dod, fluorescence intensities of *mt*Dod-*mm*ACP and *mt*Dod-*mm* ACP-H8 were compared to free *mm*ACP after labeling (Fig. 6).

**Fig. 6:**
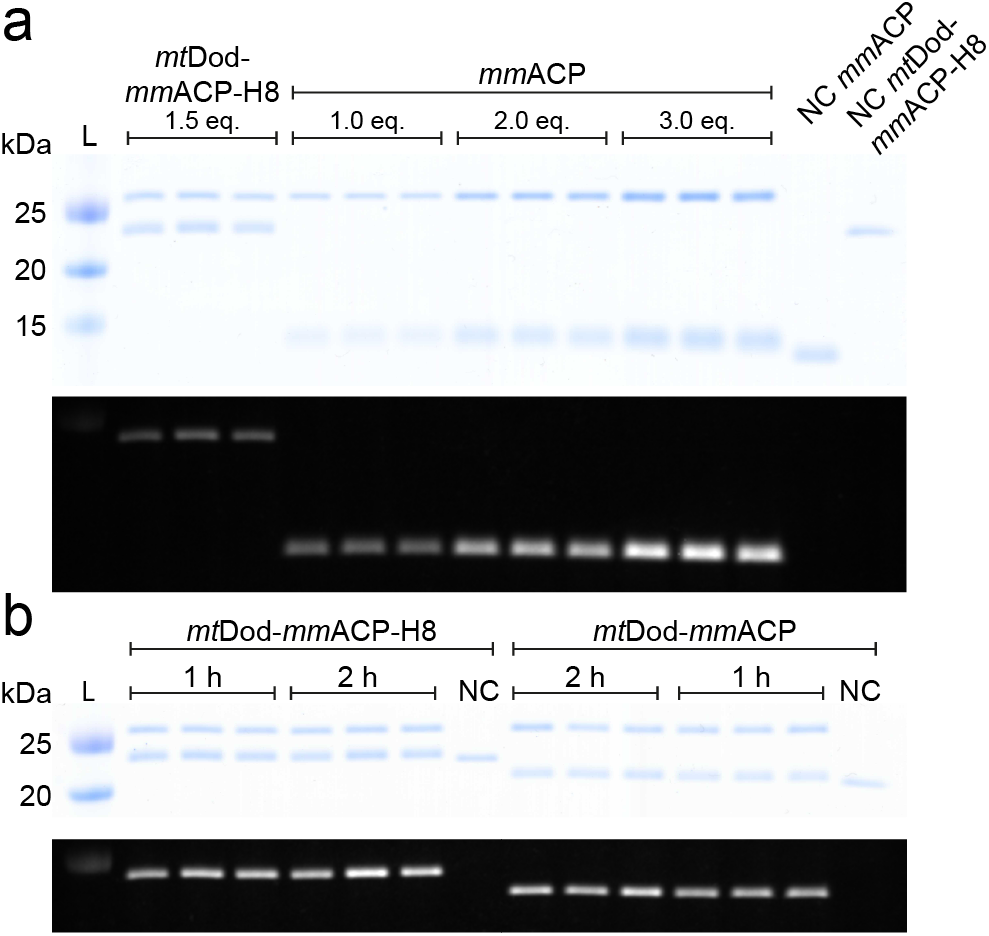
Modification of *mt*Dod-*mm*ACP, *mt*Dod-*mm*ACP-H8 and *mm*ACP by Sfp with fluorescent CoA. L: Ladder. NC: negative control reaction without Sfp. Eq.: molar equivalents of *mm* ACP loaded onto the SDS-PAGE gel. a) Top: Coomassie stained SDS-PAGE gel of the reaction solution and negative controls. Sfp, runs at an apparent molecular weight of slightly less than 25 kDa, *mt*Dod-*mm*ACP-H8 slightly more than 25 kDa and *mm*ACP at about 15 kDa. Unmodified *mm*ACP and *mt*Dod-*mm* ACP-H8 (negative controls) show lower apparent molecular weights indicating a different running behavior between CoA-488 modified and unmodified proteins. Bottom: In-gel fluorescence taken before Coomassie staining. Only proteins modified with the CoA-488 are visible. Higher bands represent labeled *mt*Dod-*mm*ACP-H8 and lower bands labeled *mm*ACP. b) Top: Coomassie stained SDS-PAGE gel of the reaction solutions of *mt*Dod-*mm* ACP-H8 and *mt*Dod-*mm* ACP after 1 h and 2 h. Bottom: In-gel fluorescence taken before Coomassie staining. For both constructs, fluorescence intensity increases at longer reaction times. For uncropped images and mtDod-mmACP blots see Fig. S8.

By comparing the fluorescence intensities of CoA-488-labeled *mt*Dod-*mm*ACP and *mt*Dod-*mm*ACP-H8 with CoA-488-labeled free *mm*ACP, the relative degree of labeling was determined to about 31% ± 8% and 36% ± 8% respectively. After an additional hour of labeling in-gel fluorescence of *mt*Dod-*mm*ACP and of *mt*Dod-*mm*ACP-H8 further increased by 14% ± 8% and 24% ± 12%, respectively. The overall low relative degree of labeling and the increase after an additional hour of reaction time indicates a reduced accessibility of *mm*ACP fused to *mt*Dod. It cannot be ruled out that the *mm*ACP fold at the *mt*Dod is instable or partly unfolded. Note that in SDS-PAGE, *mt*Dod-*mm* ACP runs at just two different apparent molecular weights corresponding to labeled and non-labeled protein (see Fig. 6). It seems that SDS-PAGE is limited in its efficiency of separating partly labeled species.

### *Mt*Dod-PAS-Pep constructs for AB production

Protein carriers are generally used for the production of ABs against peptides or proteins.^8^ In the standard approach, the peptide or the protein of interest is linked to the carrier, usually BSA or KLH, by chemical ligation.^7–9^ While the method is well-established and broadly used for AB production, problems can arise during conjugating the peptide/hapten to the carrier, e.g., owing to the low stability or solubility of the conjugate (or even for the peptide alone) or altered antigenic properties of the peptide.^34^ The dodecameric structure with the exposed termini allows *mt*Dod to be charged with 12 or 24 peptides/proteins on its surface by simply fusing the peptide/protein encoding sequence to the *mt*Dod gene. In order to evaluate the suitability of *mt*Dod for AB production, 11 fusion constructs were produced in *E. coli*, of which each is comprised of *mt*Dod, a PAS linker and a peptide of interest (Table 2). Peptide sequences originated from human heat shock proteins (HSP), proheparin-binding EGF-like growth factor (HB-EGF) and C-terminus of the heat shock cognate protein 70 interacting protein (CHIP) (for detailed peptide origin see Table S2).

**Table 2:**
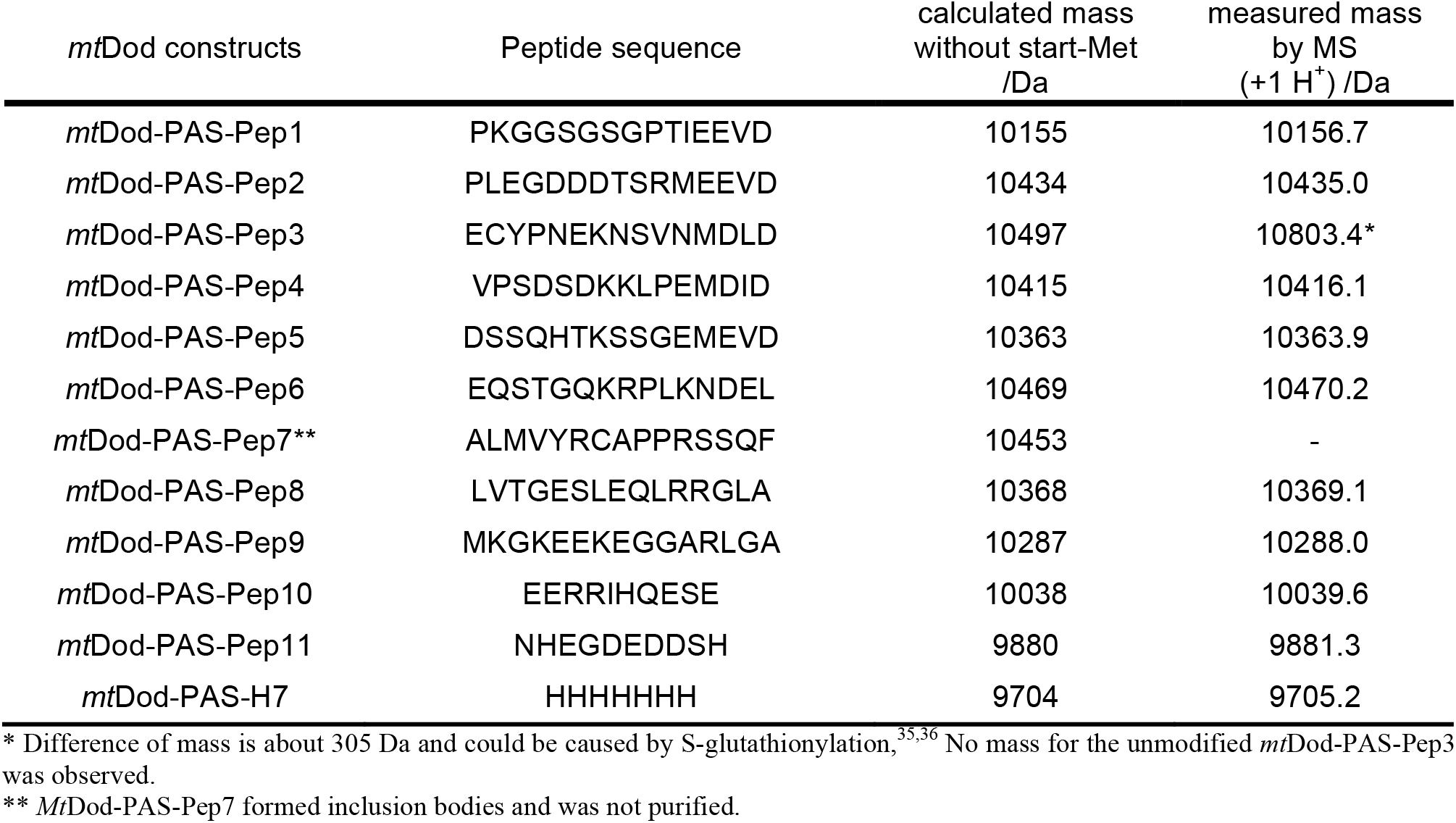
*Mt*Dod constructs for AB production. After purification by the heat treatment protocol, the correct size of the proteins was verified by ESI-MS.

Pep-encoding sequences, provided on oligonucleotide primers, were introduced in single-step by ligation-free cloning. Recombinant expressions and purifications followed the established protocols described above. All constructs were received as soluble proteins, except *mt*Dod-PAS-Pep7 that formed inclusion bodies (see Table 2). The yellow color of the inclusion bodies indicated assembled dodecamer, and we assume that aggregation of *mt*Dod-PAS-Pep7 was induced by the cysteine in Pep7 forming disulfide-bridges between the dodecamers. All constructs, except mtDod-PAS-Pep7, were further purified by two cycles of DMSO-induced precipitations. FMN was added before constructs were eventually forwarded to SEC to remove unbound FMN and remaining DMSO as well as to select for dodecameric species (Fig. 7 a). We assumed that *mt*Dod remains saturated with FMN due to its high affinity and cooperative binding mode.^19^ FMN-saturated *mt*Dod constructs (FMN:*mt*Dod constructs) can be determined in concentration by absorbance at 450 nm, and are amenable to stability measurements by the thermocyclic fluorescence assay. All constructs were received as dodecamers as indicated by SEC (Fig. S9). *Mt*Dod-PAS-Pep3 shows in addition to the dodecamer species higher oligomeric states in SEC, which we assume to result from disulfide-bridges formed by the cysteine in Pep3. The dodecamer containing fractions were pooled, and the purity was controlled by SDS-PAGE (see Fig. 7 b). The thermocyclic fluorescence assay revealed the high thermal stability of all *mt*Dod-PAS-Pep constructs, similar as the wild type protein (Fig. 7 c).^19^ Molecular masses of the constructs were measured with ESI-MS and confirmed full-length protein (see Table 2).

**Fig. 7:**
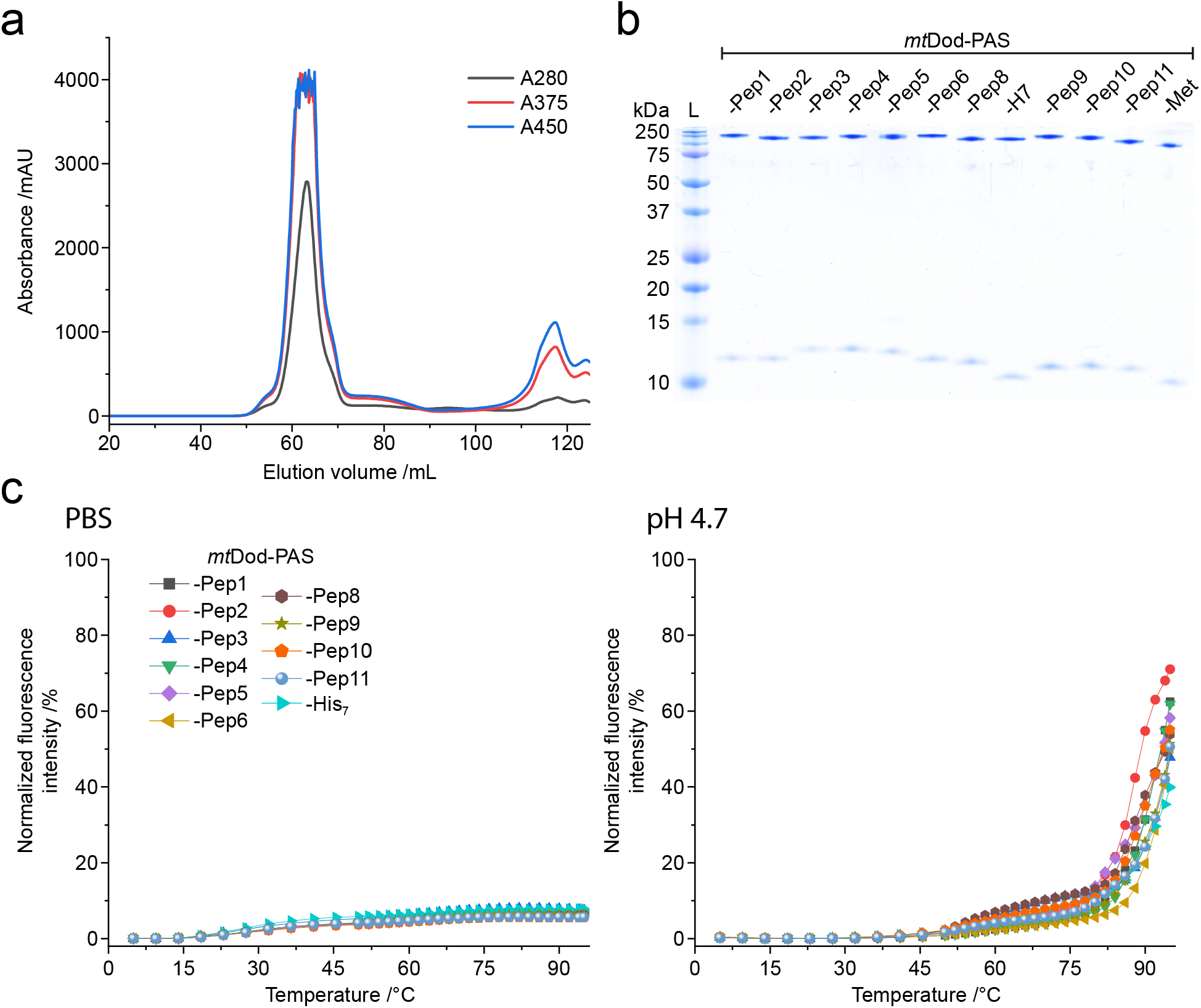
*Mt*Dod-PAS-Pep purity and stability. a) SEC profile of FMN:*mt*Dod-PAS-Pep1. The dodecameric species can be identified by the FMN absorption at 375 nm and 450 nm. In addition, unbound FMN is visible at ~117 mL. All constructs, except *mt*Dod-PAS-Pep3, showed comparable chromatographic profiles. b) SDS-PAGE gel of all purified *mt*Dod-PAS-Pep constructs. Use of standard loading buffer (pH 6.8, 2.5% SDS) at prolonged heat incubation (30 min, 95 °C) allows the observation of monomer and dodecamer. c) Thermocylic fluorescence assay of *mt*Dod-PAS-Pep constructs. Measurements were performed in PBS and at pH 4.7 as described above (see Fig. 4)

The amount of purified protein/dodecamer of *mt*Dod-PAS-Pep constructs were between 20-50 mg, of which *mt*Dod-PAS-Pep3 was received at the lowest amount (about 20 mg), which may be explained from agglomeration of dodecamers due to disulfide bridges. In general, just a fraction of the total protein received per expression was purified. The approximated (theoretical) yield per liter of *E. coli* expression culture is 200-500 mg. Constructs *mt*Dod-PAS-Pep3 and *mt*Dod-PAS-Pep7 indicate that cysteine containing peptides can cause problems when processed via the described purification strategy. A changed protocol, in which the oxidative conditions imposed by the high concentrations of FMN are avoided, could lead to improved results.

Endotoxin concentrations, measured in endotoxin units (EU) via a *Limulus* amebocyte lysate (LAL) test, were determined to avoid an endotoxin shock in immunizations. *Mt*Dod-PAS-Pep3 and Dod-PAS-Pep6 contained the highest amount of endotoxin with 73 EU/mg and 55 EU/mg, respectively; all other samples showed values less than 30 EU/mg (average of all constructs 30 ± 23 EU/mg). Since less than 0.1 mg protein was used per injection, none of the samples were critical in endotoxin levels (above 5-10 EU/kg of rabbit per injection)^37,38^.

Purified *mt*Dod constructs were eventually submitted to an AB production company for immunization in rabbits and AB purification (Davids Biotechnologie GmbH, Germany). AB productions were induced in rabbits by 5 injections of proteins over 63 days using adjuvants MF59/AddaVax or Montanide ISA 51. The ABs were purified by affinity chromatography with the respective *mt*Dod-PAS-Pep construct immobilized on the column matrix. For all the 10 *mt*Dod-PAS-Pep constructs, which were submitted to immunizations, purified ABs were obtained. We divided the obtained ABs into three quality classes based on their properties in western blotting: “Class 1” terms ABs recognizing the proteins of interest provided as recombinantly purified protein (produced in *E. coli*), “class 2” ABs do not recognize recombinant protein of interest, but protein of expected apparent molecular weight in HEK293T human cell lysates, and “class 3” ABs only recognize *mt*Dod-PAS constructs (like the respective *mt*Dod-PAS-Pep or *mt*Dod-PAS-Met). 6 of the 10 ABs were rated as “class 1” (Fig. 8), ABs derived from Dod-PAS-Pep5 & -Pep10 qualified as “class 2” (see Fig. 8 a), and the 2 remaining ABs derived from Dod-PAS-Pep6 & -Pep8 as “class 3” (western blots not shown). Overall, the antibodies did not cause recurring background signals, but showed high specificity in the used lysates (see Fig. 8 a), indicating that *mt*Dod-PAS is a suitable matrix in this respect.

**Fig. 8:**
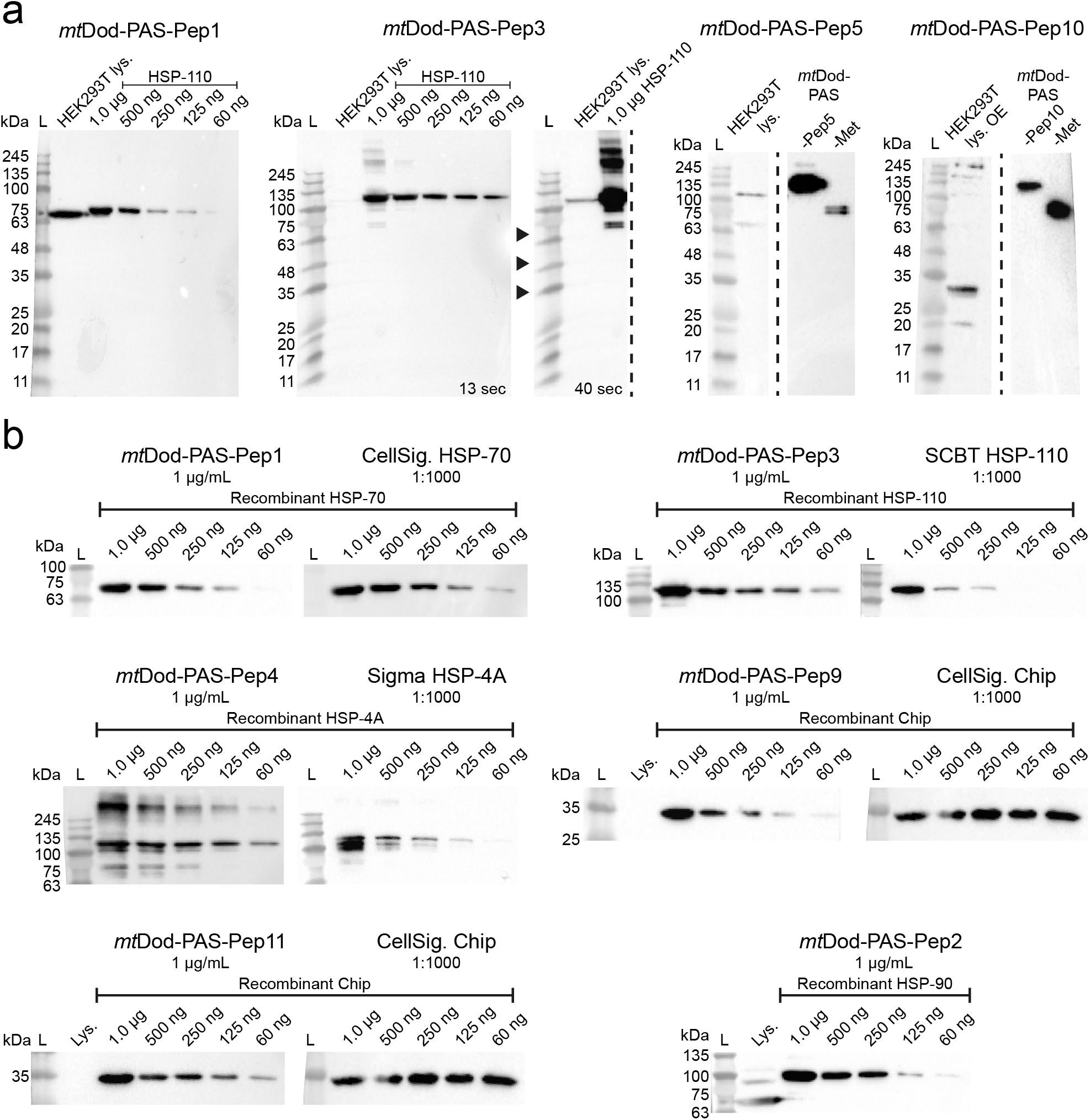
Western blots with selected *mt*Dod-PAS-Pep construct ABs. L: Ladder. Lys.: Lysate. OE: protein overexpressing cells. a) Detection of the target protein in purified form and in the lysate. ABs derived from *mt*Dod-PAS-Pep1 and *mt*Dod-PAS-Pep3 detect target proteins in the lysate. For *mt*Dod-PAS-Pep3, the same blot is shown in two exposure times. ABs derived from *mt*Dod-PAS-Pep5 and *mt*Dod-PAS-Pep10 did not recognize purified target protein, but seem to recognize a protein in overexpressing cells (uncropped western blots see Fig. S10) b) Comparison with commercially available ABs (CellSig.: Cell Signaling Technology, Inc; SCBT: Santa Cruz Biotechnology, Inc; Sigma: Sigma-Aldrich, Merck KGaA). Different amounts (1 μg to 60 ng) of purified target protein were loaded and analyzed by *mt*Dod-PAS-Pep derived ABs (1 μg/mL) and commercial available ABs (at recommended concentration). Results are summarized in Table 3. For uncropped western blots see Fig. S11. ABs derived from *mt*Dod-PAS-Pep2 also recognize HSP-70. For a comparison of *mt*Dod-PAS-Pep1-3 derived ABs, see Fig. S12 a.

For all “class 1” ABs, the recognition limit/range of purified recombinant protein was analyzed and compared with commercially available antibodies (Table 3).

**Table 3:**
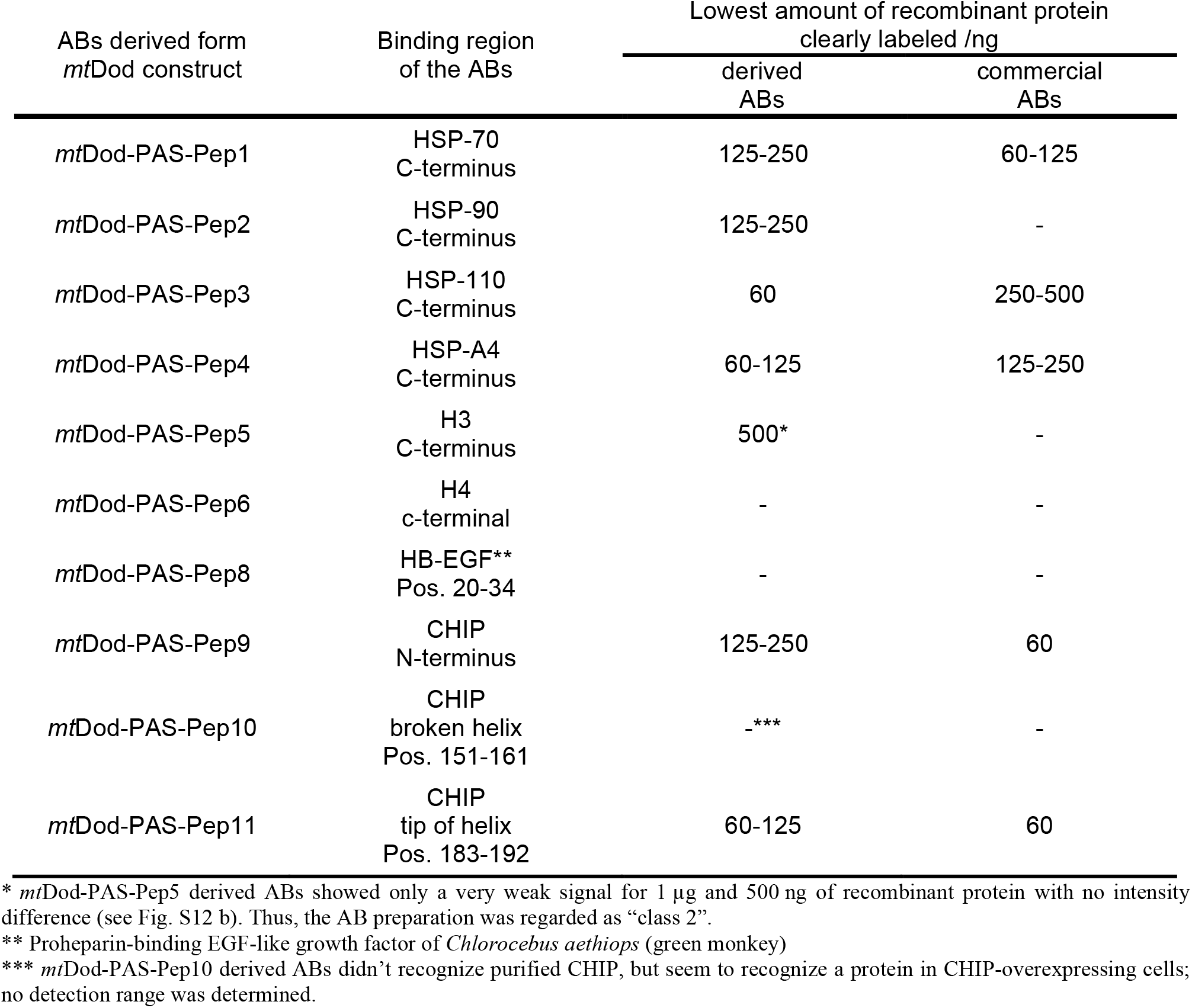
Recognition strength of *mt*Dod-PAS-Pep antibodies. The lowest amount of purified recombinant protein (in ng) that could be clearly detected with the respective ABs is given as detection limit. *Mt*Dod-PAS-Pep-derived antibodies were used at a final concentration of 1 μg/mL and commercial antibodies were used at the recommended dilution. For comparison purposes, the exposure time for each AB pair (*mt*Dod-PAS-Pep-derived and commercial) was the same, while it was varied between the different pairs.

The *mt*Dod-PAS-Pep-derived antibodies show that *mt*Dod-PAS is a well-suited carrier system for the production of peptide-specific antibodies. *Mt*Dod-PAS-based fusions benefits from the easy cloning, uncomplicated production/purification and the high protein yields. In our case study, problems in the purification of Cys-containing constructs emerged from the oxidative conditions induced by high FMN concentrations, which may, however, be overcome when working under reducing conditions or alternatively replacing cysteines with serines.

## Discussion

During the last years, dodecins have been characterized as flavin binding proteins involved in flavin homeostasis.^17,19^ In addition, the unique protein fold and particularly the exceptional flavin binding mode were harnessed in technological applications, although exclusively on the archaeal protein from *Halobacterium salinarum*.^39,40^ In this study, we present *mt*Dod as a versatile scaffold protein to attach peptides and small proteins. The *mt*Dod dodecameric fold stays intact at high temperatures, is stable over time and at various chemical conditions, which allows protein purification by quick heat-induced denaturation, protein precipitation with solvents and holds out the prospect to broadly accept conditions for chemical ligation reactions. In addition to its high stability, *mt*Dod can be produced in high yields in *E. coli*, as demonstrated by the purification of up to several hundreds of milligrams of *mt*Dod-peptide fusions per liter of bacterial culture. Proteins fused to *mt*Dod are presented at the *mt*Dod cage surface and have been shown to remain accessible and functional. Both the SpyT/-C and the ACP/Sfp system allowed attaching the cargo at the *mt*Dod surface. In this respect, *mt*Dod is comparable to the recently presented IMX313 scaffold, suggested for use in vaccine development^31^. When evaluating *mt*Dod as a scaffold, we observed that constructs can suffer from low solubility in response to the properties of the attached cargo. While agglomeration by disulfide formation, observed for *mt*Dod-PAS-Pep3 and *mt*Dod-PAS-Pep7, could simply be avoided by reducing conditions during protein preparation or replacing of cysteines with serines, solubility problems induced by hydrophobic and structurally unstable fold may be solved by heterododecamer formation to dilute the aggregation-inducing species on the *mt*Dod surface. For the construct *mt*Dod-PAS-Strep, we demonstrated that heterododecamer formation with the wild type protein is readily possible in vitro and in vivo by simply providing both proteins during refolding or recombinant protein production, respectively (see Fig. 3).

As a pilot run for evaluating the suitability of *mt*Dod as a carrier matrix, we chose 11 peptides originating from different human proteins like CHIP or HSP-70, and fused them to *mt*Dod for AB production. One of the 11 peptide constructs formed inclusion bodies, while all other constructs were purified by the standard heat-denaturation purification protocol without any need for individual adaptions. From all immunizations performed in this study, ABs that at least recognized the *mt*Dod-PAS scaffold were received, as revealed by western blotting. Overall, 8 of the 10 ABs recognized proteins in HEK293T human cell lysate at expected molecular weight in western blotting. For 6 of them, correct target recognition could be confirmed with the recombinantly purified protein as reference. No AB preparation showed any unspecific reactivity in HEK293T human cell lysate demonstrating that *mt*Dod is a suitable matrix for AB production against human proteins. Here, it seems that a benefit in the use of *mt*Dod as a carrier matrix lies in its unique fold and its absence in eukaryotes.^16,20^ We have used the *mt*Dod scaffold for the direct fusion of peptides that is not possible with the standard carrier matrices used for immunizations. Labs that are experienced in protein expression and want to produce ABs targeting peptides without relying on peptide synthesis and chemical crosslinking may find this approach attractive. We expect that the exposed termini are also suited for chemical ligation of haptens or antigens, following standard immunization protocols, or for Click chemical modification.^41^ However, in proofing the concept of dodecin for peptide immunizations, we did not elaborate on this. Finally, we note that the availability of dodecins with similar features (e.g. *Streptomyces coelicolor*, *Streptomyces davaonensis* and *Thermus thermophilus* dodecins)^18,42^ is advantageous when aiming for heterologous prime/boost protocols by using two dodecin scaffolds with low sequence identity fused with the same antigen.^43^

While the *mt*Dod has been mainly tested as carrier matrix for AB production in this study, the properties of *mt*Dod call for its application as a scaffold in a broad range of biotechnological and bioengineering applications. We encourage to explore the m*t*Dod as a scaffold when defined particles with specific surface properties are required. Such constructs can be valuable in e.g. diffusion measurements,^44^ for formation of biomaterials^45–48^ and in creating enzyme scaffolds.^32,49,50^ *Mt*Dod hetero-dodecamers may be applied for pull down assays when combining a *mt*Dod construct bearing a protein recruiting peptide and a *mt*Dod construct with a purification tag.

## Material and Methods

### Cloning

Expression constructs were cloned using standard PCR methods and In-Fusion® HD Cloning (TaKaRa Bio Europe). Primers were ordered from Sigma-Aldrich. Inserts were verified by Sanger sequencing (by Microsynth Seqlab, Göttingen, Germany). For polycistronic constructs spacer DNA sequences (between genes) were designed with EGNAS (version 1158).^51^ For a list of all constructs see Table S1.

### Expression and cell lysis

Plasmids were transformed into BL21 (DE3) Gold cells and cells were plated onto LB-agar plates containing 100 ng/μL ampicillin and 1 g/mL glucose. 10 mL LB medium with 100 ng/μL ampicillin and 1 g/mL glucose were inoculated with a single colony and incubated at 37 °C and 180 rpm overnight. 1 L TB medium with 100 ng/μL ampicillin was inoculated with 10 mL overnight LB culture, and incubated at 37 °C and 180 rpm until the OD_600_ reached about 0.8. The cultures were cooled to 20-30 °C, and the expression was induced with 1 mL 1 M IPTG solution. The cultures were incubated overnight at 20 °C 160 rpm for protein production. *Mt*Dod-PAS-Pep constructs were expressed in 500 mL TB medium induced with 500 μL 1 M IPTG. Cells were harvested at 4,000 rcf and frozen in liquid nitrogen or directly processed. For purification by heat denaturation (*mt*Dod constructs), cell pellets were resuspended in 30 mL standard dodecin buffer: 300 mM NaCl, 5 mM MgCl_2_ and 20 mM Tris-HCl (pH 7.4, adjusted with HCl). For purification by His-tag affinity chromatography, cell pellets were resuspended in 30 mL Ni-NTA wash buffer I: 200 mM NaCl, 35 mM K_2_HPO_4_ and 15 mM KH_2_PO_4_ (pH 7.4, adjusted with NaOH or HCl) and 40 mM imidazole. To the resuspended cells, PMSF and DNase I were added and cells were disrupted by French press. Cell debris was removed by centrifugation (50,000 rcf, 20 min). All steps after cell harvest were conducted at 4 °C or on ice.

### Purification by heat denaturation

The cell debris free lysates (about 30 mL) were divided in about 10 mL aliquots and incubated at 75 °C for 15 min. Lysate containing *mt*Dod-*mm*ACP or *mt*Dod-msfGFP-H8 was incubated at 55 °C or 70 °C respectively. Heat-denatured proteins were removed by centrifugation (15,000 rcf, 10 min). Proteins in the supernatant (combined from 3 aliquots 20-25 mL) were precipitated with 50% (v/v) DMSO (final concentration) and pelleted by centrifugation (15,000 rcf, 10 min). Depending on the construct, higher concentrations or other organic solvents (MeOH and acetone) might be needed (precipitation with 75% (v/v) acetone (final concentration) turned out to be fastest and most reliable, but might cause aggregation). The obtained protein pellets were dissolved in about 20 mL standard dodecin buffer at RT and afterwards cooled on ice. Precipitation was repeated once, and the pellets were dissolved in about 5 mL. Insoluble precipitate was removed by centrifugation (15,000 rcf, 10 min). To the protein solutions, FMN was added in excess (above its solubility limit) (F6750, Sigma-Aldrich: 70% pure, free RbF ≤ 6%) and samples were incubated on ice for at least 1 hour. About 250-500 μL of protein solution were further purified by SEC. For loading the column, the concentration of the protein solution was judged by the yellow tone of the solution (depending on the dodecin concentration FMN solution turn from yellow to orange-brown).

### Purification by His-tag affinity chromatography

The cleared lysate was poured on 5 mL packed Ni-NTA agarose (His60 Superflow, TaKaRa Bio Europe) gravity flow columns, pre-equilibrated with Ni-NTA wash buffer. The loaded resin was washed with 15 mL Ni-NTA wash buffer I and with 15 mL Ni-NTA wash buffer II (Ni-NTA wash buffer I with 80 mM imidazole). MtDod-msfGFP-H8 was additionally washed with 10 mL Ni-NTA wash buffer III (Ni-NTA wash buffer I with 200 mM imidazole). Proteins were eluted in 15 mL elution buffer (200 mM NaCl, 35 mM K_2_HPO_4_ and 15 mM KH_2_PO_4_ (pH 7.4, adjusted with NaOH or HCl) and 500 mM imidazole). Eluted proteins were concentrated with ultra centrifugal filters (Amicon®, Merck) with the appropriated mass cutoff and further purified by SEC.

### Refolding of *mt*Dod-*se*ACP,*mt*Dod SpyC and *mt*Dod SYNZIP constructs

Yellowish inclusion bodies obtained after cell disruption and centrifugation (50,000 rcf, 30 min) were manually separated from other solid cell debris and then washed three times (resuspended and centrifuged) with inclusion body wash buffer (137 mM NaCl, 2.7 mM KCl, 10 mM Na_2_HPO_4_, 0.5 mM KH_2_PO_4_ (PBS, pH 7.4 not adjusted), 5 mM EDTA and 2% (v/v) Triton X-100). The washed inclusion bodies were then dissolved in 10 mL GdmCL buffer (6 M guanidinium hydrochloride, 20 mM Tris-HCl (pH 8.0, adjusted with HCl)). For refolding, 0.5 mL protein solution were diluted with 4.5 mL GdmCL buffer containing L-Arginine (1 M final concentration). Refolding was performed by dialyzing twice against the 100-fold volume of the respective buffer containing 1 mM FMN (omitted for *mt*Dod SYNZIP constructs). For SpyC constructs, a phosphate borate buffer (100 mM NaCl 25 mM Na_2_HPO_4_, 25 mM H_3_BO_3_ (pH 8.5 adjusted with NaOH) and 20% glycerol) was used, and for all other constructs the standard dodecin buffer was used. Dialysis was conducted at 4 °C and without stirring for the first ~12 h (over night). Aggregated protein was removed from the solution by centrifugation (3000 rcf, 10 min). Refolded proteins were concentrated with ultra centrifugal filters (Amicon®, Merck) with the appropriate mass cutoff and further purified by SEC.

### SEC

Prior to injection, all samples were filtered with 0.22 μm membrane filters (DURAPORE®, Merck). All proteins in this study, except *mt*Dod-PAS-Pep constructs, were purified by using a equilibrated Superdex 200 increase 10/300 column (GE Healthcare) on an ÄKTA Explorer or ÄKTA Basic device. For *mt*Dod-PAS-Pep constructs, a HiLoad Superdex 200 16/600 pg was used. Running buffer for all constructs, except *mt*Dod SpyC constructs and *mt*Dod-msfGFP-H8, was the standard dodecin buffer. The flow rate was 0.5 mL/min for the Superdex 200 increase 10/300 column or 1.0 mL/min for the HiLoad Superdex 200 16/600 pg column with fraction resolution of 0.3 mL or 2.0 mL, respectively. The running buffer for the *mt*Dod SpyC constructs and *mt*Dod-msfGFP-H8 was phosphate borate buffer (as used for refolding), and the flow rate was reduced to 0.45 mL. All runs were conducted at 4 °C. Fractions were pooled and analyzed by SDS-PAGE with Coomassie staining. Pooled fractions were aliquoted, frozen in liquid nitrogen and stored at −80 °C.

### Protein concentrations

The concentration of FMN saturated *mt*Dod constructs was determined by the FMN absorption and the corresponding extinction coefficients (375 nm, 450 nm, 473 nm respective extinction coefficients 10,000 M^−1^×cm^−1^, 12,000 M^−1^×cm^−1^ and 9,200 M^−1^×cm^−1^). The concentration of fluorescent proteins were determined by the chromophore absorption and the corresponding extinction coefficient (mClover3 constructs: 506 nm, 109,000 M^−1^×cm^−1^ (ref^52^); mRuby3 constructs: 558 nm, 128,000 M^−1^×cm^−1^ (ref^52^); msfGFP constructs: 485 nm 82,400 M^−1^×cm^−1^ (ref^25^). For other proteins, the concentrations were determined by the absorption at 280 nm and applying the calculated extinction coefficient. For proteins with low amounts of bound flavin, the absorption at 280 nm was corrected using the absorptions at 450 nm and the 280/450 nm ratio of pure FMN.

### SDS-PAGE

SDS-PAGE was performed at initial 70 V (15 min) followed by 200 V (about 70 min) on Tris-Tricine gels (self casted 10% gels, as described in ref^53^) using an Mini-PROTEAN Tetra Cell system (Bio-Rad). For heterododecamer samples tris-tricine (Laemmli) gels (self casted 12% gels) and SDS free loading buffer (4×: 50% glycerol, 5 mM FMN) were used. Laemmli SDS-PAGE was performed at initial 70 V (15 min) followed by 200 V (about 150 min). For non-*mt*Dod construct samples or if full dodecamer denaturation was not necessary/wanted, standard SDS-PAGE loading buffer (4×: 50% (v/v) glycerol, 375 mM Tris-HCl (pH 6.8, adjusted with HCl), 10% (w/v) SDS, 10% (v/v) 2-mercaptoethanol, 50 mM EDTA, bromophenol blue) was used. For full denaturation of the *mt*Dod dodecamer, a 2-component acidic SDS-PAGE loading buffer (4× acidic part 1: 10% (w/v) SDS, 300 mM acetic acid; 4× acidic part 2: 50% (v/v) glycerol, 300 mM Tris (unbuffered), 200 mM Tris-HCl (pH 6.8, adjusted with HCl), 10% (v/v) 2-mercaptoethanol, 50 mM EDTA, bromophenol blue) was used. Amount of acetic acid and unadjusted Tris can be Samples were mixed with the standard SDS-PAGE loading buffer or acidic SDS-PAGE loading buffer component 1 and heat treated at 95 °C (10 min). After heat treatment, acidic SDS-PAGE loading buffer component 2 was added to the acidic samples and a second heat treatment was applied. Gels were stained over night with InstantBlue Coomassie stain (Expedeon) and imaged using a scanner (EPSON expression 1680 pro).

### Thermocyclic fluorescence assay

For stability measurements, a clear 96 well PCR plate (MLL9601; Bio-Rad) was prefilled with 23 μL of respective buffer per well; i.e., PBS or acetate buffer (150 mM NaCl, 100 mM acetic acid (pH 4.2, adjusted with NaOH)). Plates were placed on ice and 2 μL of the corresponding 50 mM *mt*Dod construct solution were added to the wells. Plates were then sealed with optical tape (iCycler iQ^®^; Bio-Rad), centrifuged (3000 rcf, 2 min) and placed into a precooled (5 °C) real-time PCR instrument (C1000™ Thermal Cycler and CFX96™ Real-Time System; Bio-Rad). For the fluorescence detection, excitation/emission filter bandwidth of 450-490/560-580 nm was used. After 1 h incubation at 5 °C, the heating and cooling cycles were started, with each cycle containing a heating phase for 6 min and a cooling (5 °C) phase for 30 min. The heating phase temperature was raised stepwise from 5 °C to 95 °C. Until 50 °C, the step size was 4.5 °C while at higher temperatures the step size was reduced to 2.0 °C. Data points were taken after each phase. The complete temperature protocol was applied to every sample. For the stability measurements of *mt*Dod-PAS-Pep constructs, the acetate buffer was replaced with a MES buffer (150 mM NaCl, 100 mM MES (pH 4.7, adjusted with HCl) and the heating phase duration was prolonged to 10 min.

### Modification of *mm*ACP,*mt*Dod-*mm*ACP and *mt*Dod-*mm*ACP-H8 with Sfp

Sfp used for phosphopantetheinylation was produced in *E. coli* (Strain M15; helper plasmid pREP4 and pQE-60 encoding Sfp) and purified by His-tag affinity chromatography, dialysis overnight against imidazole free buffer (250 mM NaCl, 2 mM MgCl_2_, 10% glycerol, 50 mM HEPES (pH 8.0 adjusted with HCl)) and SEC (HiLoad Superdex 200 16/60 pg).^54^ Holo *mm*ACP was produced in *E. coli* and purified by His-tag affinity chromatography (basic buffer: 200 mM NaCl, 35 mM K_2_HPO_4_ and 15 mM KH_2_PO_4_ (pH 7.4, adjusted with NaOH or HCl), 10% glycerol; washing buffer contained 20 mM imidazole and elution buffer contained 300 mM imidazole) and SEC (HiLoad Superdex 75 16/60 pg), similar as described for ACP-GFP by Heil and Rittner et al.^55^ Pooled SEC fractions of Sfp and *mm*ACP were aliquoted, frozen in liquid nitrogen and stored at −80 °C. The modification of *mm*ACP and *mt*Dod-*mm*ACP (both variants with H8-Tag and without) with CoA 488 (CoA modified with ATTO-TEC dye ATTO 488, NEB #S9348) by Sfp was conducted in phosphate borate buffer (pH 7.4, adjusted with HCl) at 25 °C for 1 h at following conditions: 10 μM *mt*Dod-*mm* ACP or *mm*ACP, 4 μM Sfp, 1 mM DTT, 10 mM MgCl_2_ and 10 μM CoA 488. The final reaction volume was 50 μL, and reactions were carried out in triplicates. For calibration of the fluorescent signals, 3 different amounts of each *mm*ACP-H8 reaction solution (1 μL, 2 μL, 3 μL) were analyzed.1.5 μL of the corresponding *mt*Dod-*mm*ACP construct reaction solution were used for the determination of the degree of labeling. The reaction mixtures were by analyzed by SDS-PAGE, in-gel fluorescence, using the Fusion SL Fluorescence Imaging System (Vilber Lourmat) with the emission filter F-595 Y3 (520-680 nm), and Coomassie staining. For quantification of fluorescence signals, the FusionCapt Advance Solo 4 16.08a software was used. The degree of modification of *mt*Dod-*mm*ACP was determined by linear regression of the normalized (by molar amount) fluorescence signals.

### SpyC and SnpC reactions

The reactions were carried out in phosphate borate buffer (pH 8.5) at 25 °C for 20 h. Concentration of the respective carrier constructs were 10 μM (1 eq.) and 20μM of the respective cargo constructs (2 eq.), reaction volume was 50 μL. Each protein also was separately prepared in the same concentration as used in the reaction and incubated under the same conditions. 5 μL of each reaction and control were analyzed by SDS-PAGE and Coomassie staining, for all samples the acidic loading buffer was used.

### LC-MS

For the LC-MS analysis of the *mt*Dod-PAS-Pep constructs, 500 μg protein were precipitated with 75% (v/v) acetone (−20 °C, final concentration), pelleted by centrifugation (20,000 rcf, 5 min) and dissolved in 100 μL water. After removal of undissolved aggregates by centrifugation (20,000 rcf, 5 min), the solution was diluted with 5% (v/v) acetonitrile:water to a final concentration of 0.1 mg/mL. The injection/sample size was 2.5 μL (250 ng). Samples were analyzed by using a Dionex UltiMate 3000 RSLC (Thermo Fischer Scientific) coupled to a micrOTOF-Q II (Bruker Daltonik GmbH) equipped with an electrospray ionization source. Chromatographic separation (further desalting) was performed on a Discovery® BIO Wide Pore C5 column (100 × 2.1 mm, particle size 3 μm, Sigma-Aldrich) at 55 °C with a mobile-phase system consisting of water and acetonitrile (each containing 0.1% formic acid). A linear gradient ranging from 5% to 95% acetonitrile over 14 min at a flow rate of 0.4 mL min^−1^ was used. MS data was acquired in positive mode in the range from 200-2500 m/z and later analyzed using DataAnalysis 4.0 software (Bruker Daltonik GmbH).

### Western blots

Samples separated by SDS-PAGE were transferred to 0.45 μm nitrocellulose membranes at 100 V for 35-40 min in a cooled Criterion Blotter filled with transfer buffer at 4 °C (transfer buffer: 25 mM Tris, 192 mM glycine, 20 % (v/v) methanol). Membranes were blocked in 5% (w/v) milk powder in TBS (50 mM TRIS-HCl pH 7.6, 150 mM NaCl) with 0.1% Tween-20 added (TBST) for 1 h at RT, before addition of primary AB in indicated dilutions and incubation over night at 4°C. Membranes were washed twice with TBST, followed by addition of secondary AB in a 1:3000 dilution in blocking buffer for 1 h at RT. Finally, membranes were washed twice with TBST. Chemiluminescence was developed with SuperSignal® West Pico PLUS, and images were acquired with a ChemiDoc MP imaging system and quantified using Image Lab 5.0 software. Recombinant proteins used for AB comparison were produced in *E. coli* and purified by affinity chromatography and SEC. Proteins HSP-70, HSP-90, HSP-110 and HSP-A4 were expressed based on pPROEX vector system with a N-terminal His-tag that was removed by a TEV protease after affinity chromatography. CHIP was expressed based on a pGEX-6P1 vector system with a N-terminal GST-tag that was removed by the PreScission protease after affinity chromatography. All ABs were stored at −20 °C prior use.

## Supporting Information

**Fig. S1:** SDS-PAGE gel of supernatant after heat denaturation at different temperatures.

**Fig. S2:** SEC chromatograms of various *mt*Dod constructs.

**Fig. S3:** Comparison of SEC chromatograms of *mt*Dod-msfGFP-H8 and *mt*Dod *mm*ACP constructs purified by Ni-NTA affinity chromatography and/or by heat denaturation.

**Fig. S4:** SDS-PAGE gel of *mt*Dod SpyC constructs refolded under different conditions via dialysis.

**Fig. S5:** SDS-PAGE gel for comparison of loading buffers.

**Fig. S6:** SDS-PAGE gel of *mt*Dod-*mm*ACP precipitating during heat denaturation and other samples.

**Fig. S7:** HPLC-SEC chromatograms of *mt*Dod SYNZIP constructs and SYNZIP fluorescence protein constructs.

**Fig. S8:** Uncropped SDS-PAGE gels of the modification of *mt*Dod-*mm*ACP and *mt*Dod-*mm*ACP-H8 by Sfp.

**Fig. S9:** SEC chromatograms of *mt*Dod-PAS-Pep constructs, *mt*Dod-PAS-H7 and *mt*Dod(WT)

**Fig. S10:** Uncropped western blots of Fig. 8 a.

**Fig. S11:** Uncropped western blots of Fig. 8 b.

**Fig. S12:** Additional western blots with *mt*Dod-PAS-Pep derived ABs.

**Table S1:** Table of encoding sequences for all constructs used in this study, except recombinant proteins used in western blotting.

**Table S2:** Amino acid sequence of proteins from which the peptide sequences were selected for the

*mt*Dod-PAS-Pep constructs.

## Author contributions

F.B. cloned several protein expression constructs and prepared several proteins. F.B. further established protocols, designed experiments, analyzed data, visualized data, and designed research. Y.K. cloned protein expression constructs and prepared proteins used for western blotting (except *mt*Dod constructs). Y.K. also conducted the western blots and analyzed data. J.L. (under supervision of F.B.) cloned several protein expression constructs, purified *mt*Dod-PAS-Pep constructs, prepared several proteins, and analyzed data. I.G. cloned *mt*Dod-PAS-Pep constructs and expressed them in *E. coli*. H.B. designed research. R.M.V. selected peptide sequences used for AB production. M.G. analyzed data and designed research. F.B. and M.G. wrote the paper. Y.K., R.M.V., and H.B. reviewed and edited the paper.

## Acknowledgements

We thank Kim Remans for helpful discussions. We are grateful to Ilka Siebels for providing Sfp and Alexander Rittner for providing holo *mm*ACP-H8. We thank Emily Hensch and Dominik Scheliu, who assisted in this project during her bachelor thesis and master thesis, respectively.

## Abbreviations

ACP: acyl carrier protein
AB: antibody
BSA: bovine serum albumin
Catcher: small protein fold (SpyCatcher or SnoopCatcher) that binds and reacts with Tag
CellSig: Cell Signaling Technology
CHIP: C-terminus of heat shock cognate 70 interacting protein
EU: endotoxin units
FMN: riboflavin-5’-phosphate
GFP: green fluorescent protein
GSG, PAS, PAS2 GPAS, GPAS2, PASG, PAS2G: linker systems, see Table 1
HB-EGF: proheparin-binding EGF-like growth factor
HSP: heat shock protein
IPTG: isopropyl-β-D-thiogalactopyranosid
KLH: keyhole limpet hemocyanin
L: Ladder (only used in figures)
LAL: *Limulus* amebocyte lysate
Lys: lysate (only used in figures)
MAP: multiple antigen peptides
*mm*ACP: *Mus musculus* acyl carrier protein
msfGFP: monomeric superfolder green fluorescent protein
*mt*Dod: *Mycobacterium tuberculosis* dodecin
*mt*Dod(WT): *Mycobacterium tuberculosis* dodecin wild type
OD600: optical density at 600 nm
OE: over expressing cells
RSA: rabbit serum albumin
SCBT: Santa Cruz Biotechnology
*se*ACP: *Saccharopolyspora erythraea* acyl carrier protein
SEC: size exclusion chromatography
Sfp: 4’-phosphopantetheine transferase from *Bacillus subtilis*
Sigma: Sigma-Aldrich
SnpC: SnoopCatcher
SnpT: SnoopTag
SpyC: SpyCatcher
SpyT: SpyTag
SZ: SYNZIP domain
Tag: small peptide sequence that interacts with Catcher’s (SpyTag or SnoopTag)
TB: terrific broth
TBS: Tris-HCl buffered saline
TBST: Tris-HCl buffered saline with Tween-20
TT: tetanus toxoid
VLP: virus-like particle.

## Literature

1. Tolar, P., Hanna, J., Krueger, P. D. & Pierce, S. K. The Constant Region of the Membrane Immunoglobulin Mediates B Cell-Receptor Clustering and Signaling in Response to Membrane Antigens. Immunity 30, 44–55 (2009).

2. Bachmann, M. F. & Zinkernagel, R. M. NEUTRALIZING ANTIVIRAL B CELL RESPONSES. Annu. Rev. Immunol. 15, 235–270 (1997).

3. Chauhan, V., Rungta, T., Goyal, K. & Singh, M. P. Designing a multi-epitope based vaccine to combat Kaposi Sarcoma utilizing immunoinformatics approach. Sci. Rep. 9, (2019).

4. Schubert, B. & Kohlbacher, O. Designing string-of-beads vaccines with optimal spacers. Genome Med. 8, (2016).

5. Bennett, N. R., Zwick, D. B., Courtney, A. H. & Kiessling, L. L. Multivalent Antigens for Promoting B and T Cell Activation. ACS Chem. Biol. 10, 1817–1824 (2015).

6. Mariani, M. et al. Immunogenicity of a free synthetic peptide: Carrier-conjugation enhances antibody affinity for the native protein. Mol. Immunol. 24, 297–303 (1987).

7. Grant, G. A. Synthetic Peptides for Production of Antibodies that Recognize Intact Proteins. Curr. Protoc. Mol. Biol. 59, 11.16.1–11.16.19 (2002).

8. Trier, N. H., Hansen, P. R. & Houen, G. Production and characterization of peptide antibodies. Methods 56, 136–144 (2012).

9. Peeters, J. M., Hazendonk, T. G., Beuvery, E. C. & Tesser, G. I. Comparison of four bifunctional reagents for coupling peptides to proteins and the effect of the three moieties on the immunogenicity of the conjugates. J. Immunol. Methods 120, 133–143 (1989).

10. Sakarellos-Daitsiotis, M., Krikorian, D., Panou-Pomonis, E. & Sakarellos, C. Artificial Carriers: A Strategy for Constructing Antigenic/Immunogenic Conjugates. Curr. Top. Med. Chem. 6, 1715–1735 (2006).

11. Hume, H. K. C. et al. Synthetic biology for bioengineering virus-like particle vaccines. Biotechnol. Bioeng. 116, 919–935 (2019).

12. Singh, K. V., Kaur, J., Varshney, G. C., Raje, M. & Suri, C. R. Synthesis and Characterization of Hapten-Protein Conjugates for Antibody Production against Small Molecules. Bioconjug. Chem. 15, 168–173 (2004).

13. Adamczyk, M. et al. Characterization of Protein-Hapten Conjugates. 1. Matrix-Assisted Laser Desorption Ionization Mass Spectrometry of Immuno BSA-Hapten Conjugates and Comparison with Other Characterization Methods. Bioconjug. Chem. 5, 631–635 (1994).

14. Harris, J. R. & Markl, J. Keyhole limpet hemocyanin (KLH): a biomedical review. Micron 30, 597–623 (1999).

15. Swaminathan, A., Lucas, R. M., Dear, K. & McMichael, A. J. Keyhole limpet haemocyanin – a model antigen for human immunotoxicological studies. Br. J. Clin. Pharmacol. 78, 1135–1142 (2014).

16. Bieger, B., Essen, L.-O. & Oesterhelt, D. Crystal Structure of Halophilic Dodecin: A Novel, Dodecameric Flavin Binding Protein from Halobacterium salinarum. Structure 11, 375–385 (2003).

17. Grininger, M., Staudt, H., Johansson, P., Wachtveitl, J. & Oesterhelt, D. Dodecin Is the Key Player in Flavin Homeostasis of Archaea. J. Biol. Chem. 284, 13068–13076 (2009).

18. Meissner, B., Schleicher, E., Weber, S. & Essen, L. O. The dodecin from Thermus thermophilus, a bifunctional cofactor storage protein. J. Biol. Chem. 282, 33142–33154 (2007).

19. Bourdeaux, F. et al. Flavin Storage and Sequestration by Mycobacterium tuberculosis Dodecin. ACS Infect. Dis. 4, 1082–1092 (2018).

20. Liu, F. et al. Structural and biophysical characterization of Mycobacterium tuberculosis dodecin Rv1498A. J. Struct. Biol. 175, 31–38 (2011).

21. Zakeri, B. et al. Peptide tag forming a rapid covalent bond to a protein, through engineering a bacterial adhesin. Proc. Natl. Acad. Sci. 109, E690–E697 (2012).

22. Li, L., Fierer, J. O., Rapoport, T. A. & Howarth, M. Structural Analysis and Optimization of the Covalent Association between SpyCatcher and a Peptide Tag. J. Mol. Biol. 426, 309–317 (2014).

23. Veggiani, G. et al. Programmable polyproteams built using twin peptide superglues. Proc. Natl. Acad. Sci. 113, 1202–1207 (2016).

24. Thompson, K. E., Bashor, C. J., Lim, W. A. & Keating, A. E. SYNZIP protein interaction toolbox: in vitro and in vivo specifications of heterospecific coiled-coil interaction domains. ACS Synth Biol 1, 118–29 (2012).

25. Pédelacq, J.-D., Cabantous, S., Tran, T., Terwilliger, T. C. & Waldo, G. S. Engineering and characterization of a superfolder green fluorescent protein. Nat. Biotechnol. 24, 79 (2006).

26. Jevševar, S. et al. Production of Nonclassical Inclusion Bodies from Which Correctly Folded Protein Can Be Extracted. Biotechnol. Prog. 21, 632–639 (2005).

27. Ishii, M., Kunimura, J. S., Jeng, H. T., Penna, T. C. V. & Cholewa, O. Evaluation of the pH-and thermal stability of the recombinant green fluorescent protein (GFP) in the presence of sodium chloride. in Applied Biochemistry and Biotecnology 555–571 (Springer, 2007).

28. Shieh, Y.-W. et al. Operon structure and cotranslational subunit association direct protein assembly in bacteria. Science 350, 678–680 (2015).

29. Grininger, M., Zeth, K. & Oesterhelt, D. Dodecins: A Family of Lumichrome Binding Proteins. J. Mol. Biol. 357, 842–857 (2006).

30. Brune, K. D. et al. Plug-and-Display: decoration of Virus-Like Particles via isopeptide bonds for modular immunization. Sci. Rep. 6, (2016).

31. Brune, K. D. et al. Dual Plug-and-Display Synthetic Assembly Using Orthogonal Reactive Proteins for Twin Antigen Immunization. Bioconjug. Chem. 28, 1544–1551 (2017).

32. Jia Lili, Minamihata Kosuke, Ichinose Hirofumi, Tsumoto Kouhei & Kamiya Noriho. Polymeric SpyCatcher Scaffold Enables Bioconjugation in a Ratio-Controllable Manner. Biotechnol. J. 12, 1700195 (2017).

33. Yin, J., Lin, A. J., Golan, D. E. & Walsh, C. T. Site-specific protein labeling by Sfp phosphopantetheinyl transferase. Nat. Protoc. 1, 280–285 (2006).

34. Briand, J. P., Muller, S. & Van Regenmortel, M. H. V. Synthetic peptides as antigens: Pitfalls of conjugation methods. J. Immunol. Methods 78, 59–69 (1985).

35. Hill, B. G., Ramana, K. V., Cai, J., Bhatnagar, A. & Srivastava, S. K. MEASUREMENT AND IDENTIFICATION OF S-GLUTATHIOLATED PROTEINS. Methods Enzymol. 473, 179–197 (2010).

36. Zhang, H. et al. Glutathionylation of the Bacterial Hsp70 Chaperone DnaK Provides a Link between Oxidative Stress and the Heat Shock Response. J. Biol. Chem. 291, 6967–6981 (2016).

37. Wachtel, R. E. & Tsuji, K. Comparison of limulus amebocyte lysates and correlation with the United States Pharmacopeial pyrogen test. Appl. Environ. Microbiol. 33, 1265–1269 (1977).

38. Pearson, F. C. et al. Comparison of several control standard endotoxins to the National Reference Standard Endotoxin--an HIMA collaborative study. Appl. Environ. Microbiol. 50, 91–93 (1985).

39. Nöll, G., Trawöger, S., von Sanden-Flohe, M., Dick, B. & Grininger, M. Blue-Light-Triggered Photorelease of Active Chemicals Captured by the Flavoprotein Dodecin. ChemBioChem 10, 834–837 (2009).

40. Gutiérrez Sánchez, C., Su, Q., Schönherr, H., Grininger, M. & Nöll, G. Multi-Ligand-Binding Flavoprotein Dodecin as a Key Element for Reversible Surface Modification in Nano-biotechnology. ACS Nano 9, 3491–3500 (2015).

41. Kolb, H. C., Finn, M. G. & Sharpless, K. B. Click Chemistry: Diverse Chemical Function from a Few Good Reactions. Angew. Chem. Int. Ed. 40, 2004–2021 (2001).

42. Bourdeaux, F. et al. Comparative biochemical and structural analysis of the flavin-binding dodecins from Streptomyces davaonensis and Streptomyces coelicolor reveals striking differences with regard to multimerization. Microbiology (2019) doi:10.1099/mic.0.000835.

43. Lu, S. Heterologous Prime-Boost Vaccination. Curr. Opin. Immunol. 21, 346–351 (2009).

44. Gutiérrez Sánchez, C., Su, Q., Wenderhold-Reeb, S. & Nöll, G. Nanomechanical properties of protein–DNA layers with different oligonucleotide tethers. RSC Adv. 6, 56467–56474 (2016).

45. Chen, A. Y. et al. Synthesis and patterning of tunable multiscale materials with engineered cells. in Nature materials (2014). doi:10.1038/nmat3912.

46. Botyanszki, Z., Tay, P. K. R., Nguyen, P. Q., Nussbaumer, M. G. & Joshi, N. S. Engineered catalytic biofilms: Site-specific enzyme immobilization onto E. coli curli nanofibers. Biotechnol. Bioeng. 112, 2016–2024 (2015).

47. Sun, F., Zhang, W.-B., Mahdavi, A., Arnold, F. H. & Tirrell, D. A. Synthesis of bioactive protein hydrogels by genetically encoded SpyTag-SpyCatcher chemistry. Proc. Natl. Acad. Sci. 111, 11269–11274 (2014).

48. Fletcher, J. M. et al. Self-Assembling Cages from Coiled-Coil Peptide Modules. Science 340, 595–599 (2013).

49. Giessen, T. W. & Silver, P. A. A Catalytic Nanoreactor Based on in Vivo Encapsulation of Multiple Enzymes in an Engineered Protein Nanocompartment. ChemBioChem 17, 1931–1935 (2016).

50. Siu, K.-H. et al. Synthetic scaffolds for pathway enhancement. Curr. Opin. Biotechnol. 36, 98–106 2015.

51. Kick, A., Bönsch, M. & Mertig, M. EGNAS: an exhaustive DNA sequence design algorithm. BMC Bioinformatics 13, 138 (2012).

52. Bajar, B. T. et al. Improving brightness and photostability of green and red fluorescent proteins for live cell imaging and FRET reporting. Sci. Rep. 6, (2016).

53. Schägger, H. Tricine–SDS-PAGE. Nat. Protoc. 1, 16–22 (2006).

54. Mofid, M. R., Marahiel, M. A., Ficner, R. & Reuter, K. Crystallization and preliminary crystallographic studies of Sfp: a phosphopantetheinyl transferase of modular peptide synthetases. Acta Crystallogr. D Biol. Crystallogr. 55, 1098–1100 (1999).

55. Heil, C. S., Rittner, A., Goebel, B., Beyer, D. & Grininger, M. Site-Specific Labelling of Multidomain Proteins by Amber Codon Suppression. Sci. Rep. 8, (2018).

