## Supporting Figures and Tables for "Dodecin as carrier protein for immunizations and bioengineering applications"

Author affiliations:
(a) Institute of Organic Chemistry and Chemical Biology, Buchmann Institute for Molecular Life Sciences, Cluster of Excellence for Macromolecular Complexes, Goethe University Frankfurt, Max-von-Laue-Str. 15, D-60438 Frankfurt am Main, Germany
(b) Institute of Biophysical Chemistry, Buchmann Institute for Molecular Life Sciences, Goethe University Frankfurt, Max-von-Lauer Str. 15, D-60438 Frankfurt am Main, Germany
(c) European Molecular Biology Laboratory, 69117 Heidelberg, Germany; currently PROGEN Biotechnik GmbH, 69123 Heidelberg, Germany
(d) Institute of Biochemistry, Charité University Medicine, Charitéplatz 1, D-10117 Berlin

<u>Contents:</u>


| | | |
|---|---|---|
| Page | 2 | Abbreviations |
| Pages | 3-14 | Supporting Figures S1-S12 |
| Pages | 15-22 | Supporting Table S1-S2 |
| Page | 23 | Literature |

**Abbreviations:**

ACP, acyl carrier protein; AB, antibody; BSA, bovine serum albumin; Catcher, small protein fold (SpyCatcher or SnoopCatcher) that binds and reacts with Tag; CDS, coding sequence; CellSig., Cell Signaling Technology; CHIP, C-terminus of heat shock cognate 70 interacting protein; EU, endotoxin units; FMN, riboflavin-5'-phosphate; GFP, green fluorescent protein; GSG, PAS, PAS2 GPAS, GPAS2, PASG, PAS2G, linker systems, see Table 1; HB-EGF, proheparin-binding EGF-like growth factor; HSP, heat shock protein; IPTG, isopropyl-β-D-thiogalactopyranosid; KLH, keyhole limpet hemocyanin; L, Ladder (only used in figures); LAL, *Limulus* amebocyte lysate; Lys., lysate (only used in figures); MAP, multiple antigen peptides; mCHIP, middle fragment of C-terminus of heat shock cognate 70 interacting protein; MG, proteasome inhibitor MG-132; *mm*ACP, *Mus musculus* acyl carrier protein; msfGFP, monomeric superfolder green fluorescent protein; *mt*Dod, *Mycobacterium tuberculosis* dodecin; *mt*Dod(WT), *Mycobacterium tuberculosis* dodecin wild type; OD600, optical density at 600 nm; OE, over expressing cells; RSA, rabbit serum albumin; SCBT, Santa Cruz Biotechnology; *se*ACP, *Saccharopolyspora erythraea* acyl carrier protein; SEC, size exclusion chromatography; Sfp, 4'-phosphopantetheine transferase from *Bacillus subtilis*; Sigma, Sigma-Aldrich; SnpC, SnoopCatcher; SnpT, SnoopTag; SpyC, SpyCatcher; SpyT, SpyTag; SZ, SYNZIP domain; Tag, small peptide sequence that interacts with Catcher's (SpyTag or SnoopTag); TB, terrific broth; TBS, Tris-HCl buffered saline; TBST, Tris-HCl buffered saline with Tween-20; TT, tetanus toxoid; VLP, virus-like particle.

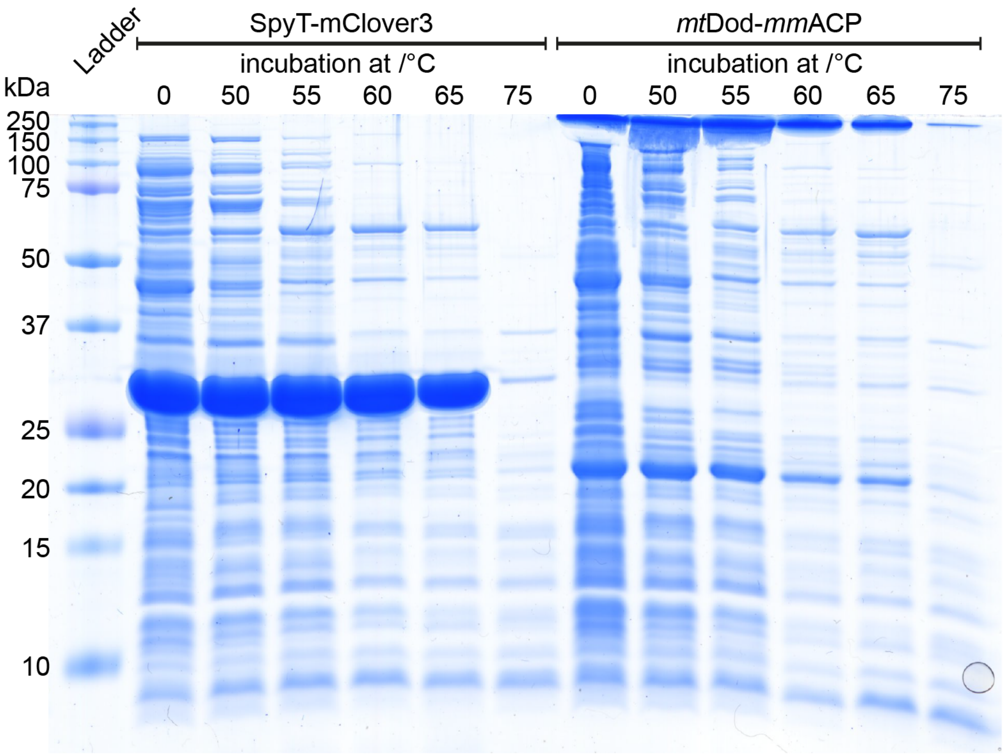


**Fig. S1:** SDS-PAGE gel of supernatant after heat denaturation at different temperatures. Standard loading buffer (pH 6.8, 2.5% SDS) was used for sample preparation with 15 min heat treatment (95 °C). Under these conditions, the dodecamer stayed largely intact (band at top boarder of the gel), and only a small fraction of monomer is observable (band at about 20 kDa). At temperatures higher than 55 °C, *mt*Dod-*mm*ACP concentration dropped, indicated by the weaker monomer band, and at 75 °C, nearly all *mt*Dod-*mm*ACP is precipitated during the heat denaturing step (see **Fig. S6** for SDS-PAGE of the pellet). Left: Heat denaturation of SpyT-mClover3 at different temperatures, performed to test the suitability of the method for mClover3 constructs.

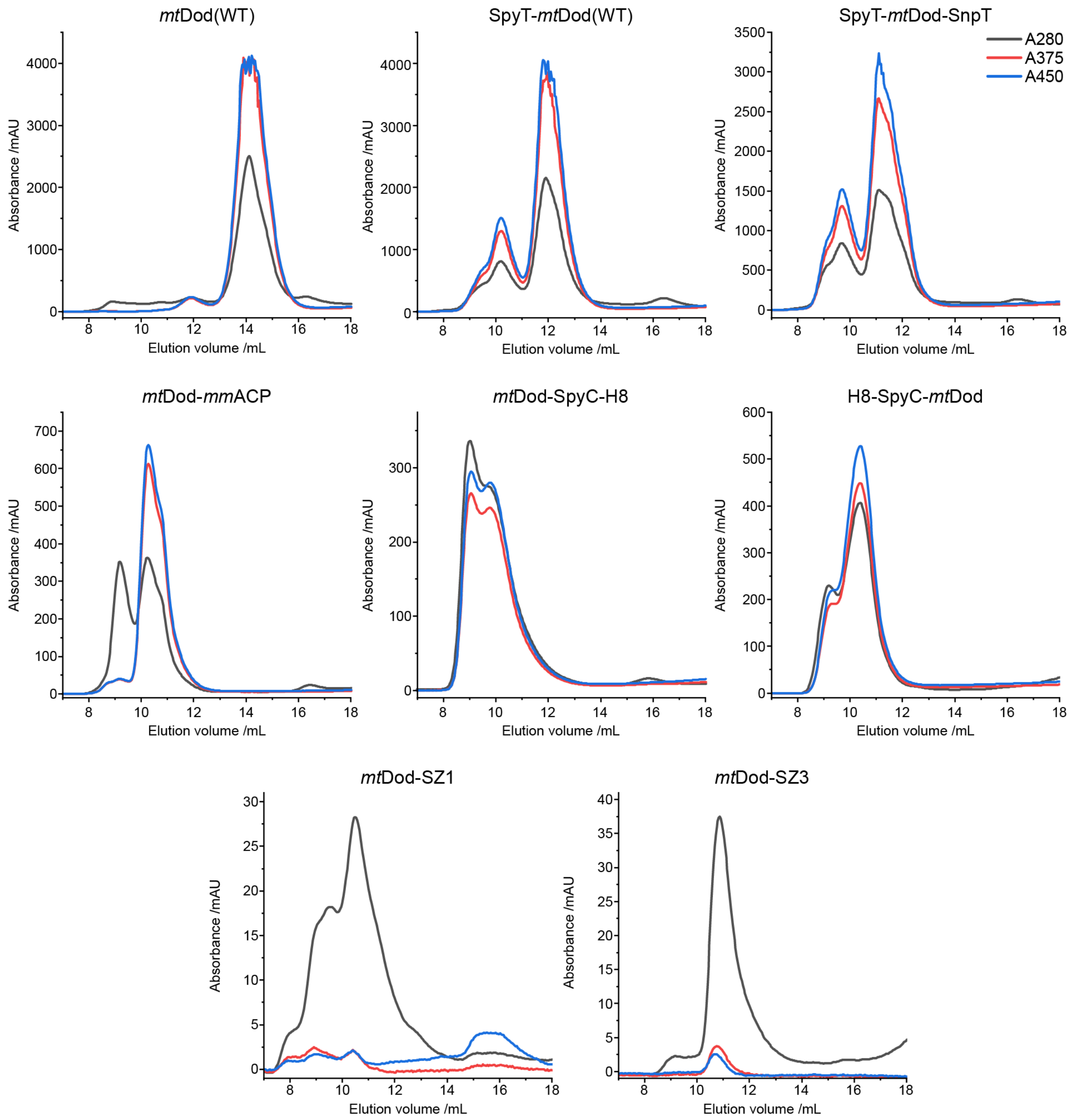


**Fig. S2:** SEC chromatograms of various *mt*Dod constructs. Used column: Superdex 200 increase 10/300. Top row: Example of *mt*Dod constructs purified by the heat denaturation protocol. The second DMSO precipitation step was used to concentrate samples as high as possible (buffer added in small portions, until the pellet was nearly fully dissolved but not completely). Under these high concentrations, some *mt*Dod constructs seem to form aggregates that still bind FMN. Chromatogram of *mt*Dod-*mm*ACP purified by the heat denaturation strategy shows a prominent aggregation peak at about 9 mL. Chromatograms of refolded *mt*Dod SpyC constructs form FMN binding dodecamers, but tend to aggregate (peak at about 9 mL). H8-SpyC-*mt*Dod seems to have lower aggregation tendencies than *mt*Dod-SpyC-H8 (the separation of dodecamer and aggregates was not possible). *Mt*Dod SYNZIP constructs were refolded and purified without additional FMN. The high aggregation tendencies of SYNZIP constructs (especially *mt*Dod-SZ1) made the purification of higher amounts challenging, as samples tended to suddenly precipitate during concentration and filtration at higher concentrations.

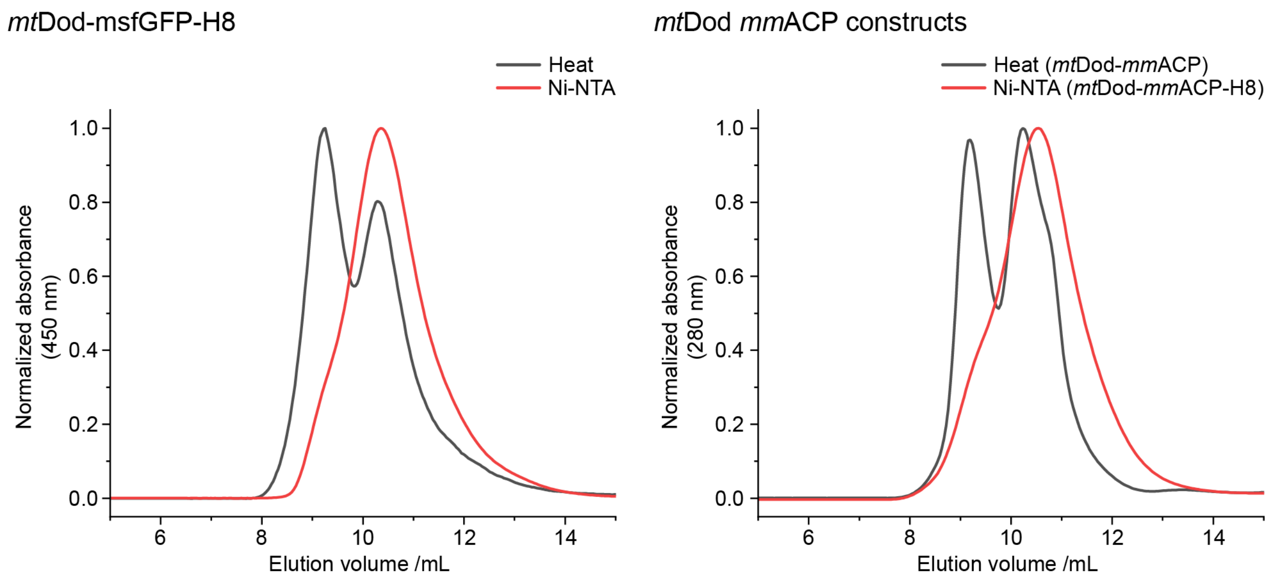


**Fig. S3:** Comparison of SEC chromatograms of *mt*Dod-msfGFP-H8 and *mt*Dod *mm*ACP constructs purified by Ni-NTA affinity chromatography and/or by heat denaturation. In both cases, the Ni-NTA affinity chromatography purification caused less aggregation. However, dodecameric fractions of *mt*Dod-msfGFP-H8 and *mt*Dod-*mm*ACP could also be received from the heat denaturation protocol.

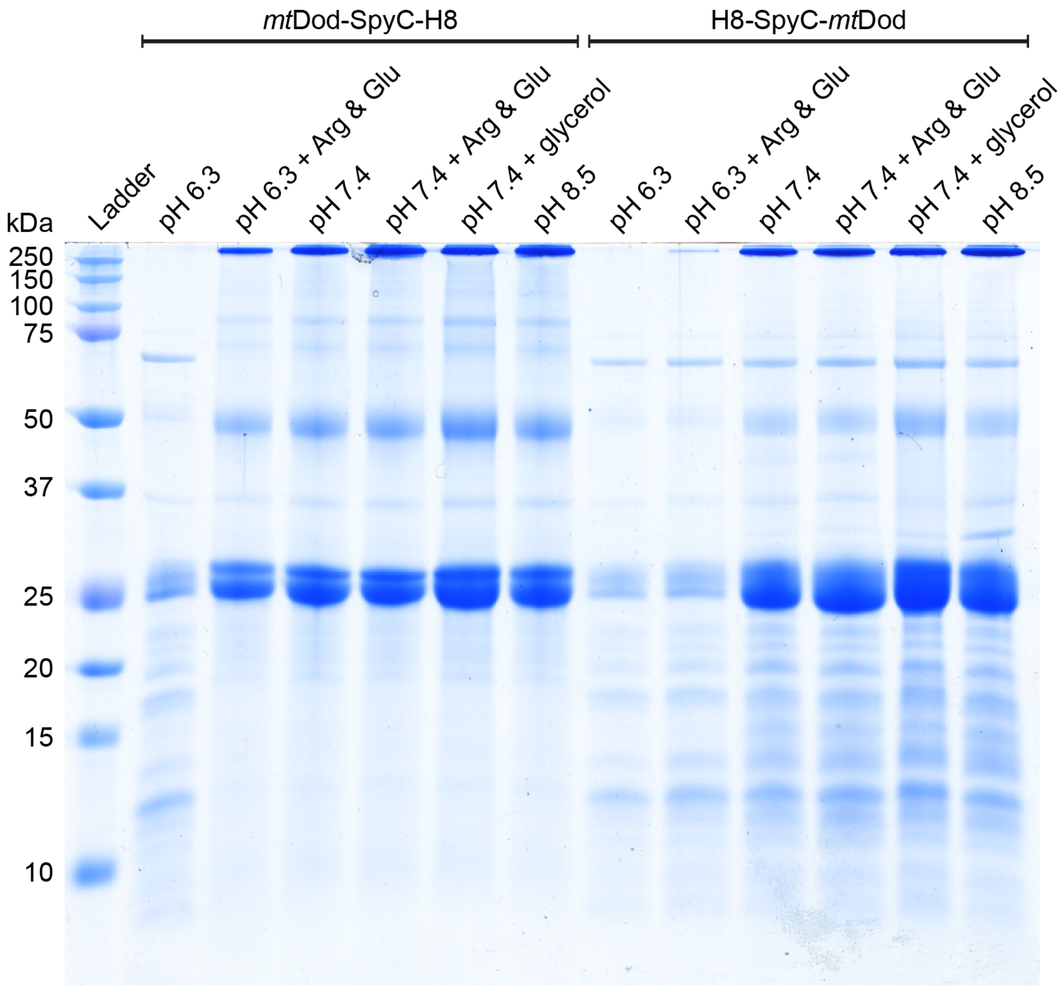


**Fig. S4:** SDS-PAGE gel of *mt*Dod SpyC constructs refolded under different conditions via dialysis. Acidic loading buffer used for sample preparation contained 50 mM acetic acid, which did not allow a full denaturation of the dodecamer. General buffer solution for refolding (final concentrations): 100 mM NaCl, 25 mM $Na_2HPO_4$ and 25 mM boric acid. The pH was adjusted to the shown values with HCl or NaOH. Refolding additives were 50 mM arginine and 50 mM glutamic acid (Arg & Glu)[1] and 20% glycerol (glycerol). Except under slightly acidic conditions, refolding of *mt*Dod SpyC constructs is possible. Arginine and glutamic acid seem to be beneficial for refolding (clearly seen for *mt*Dod-SpyC-H8 at pH 6.3). Overall, the best tested condition seems to be pH 7.4 with 20% glycerol (highest band intensity and lowest amount of precipitate was observed).

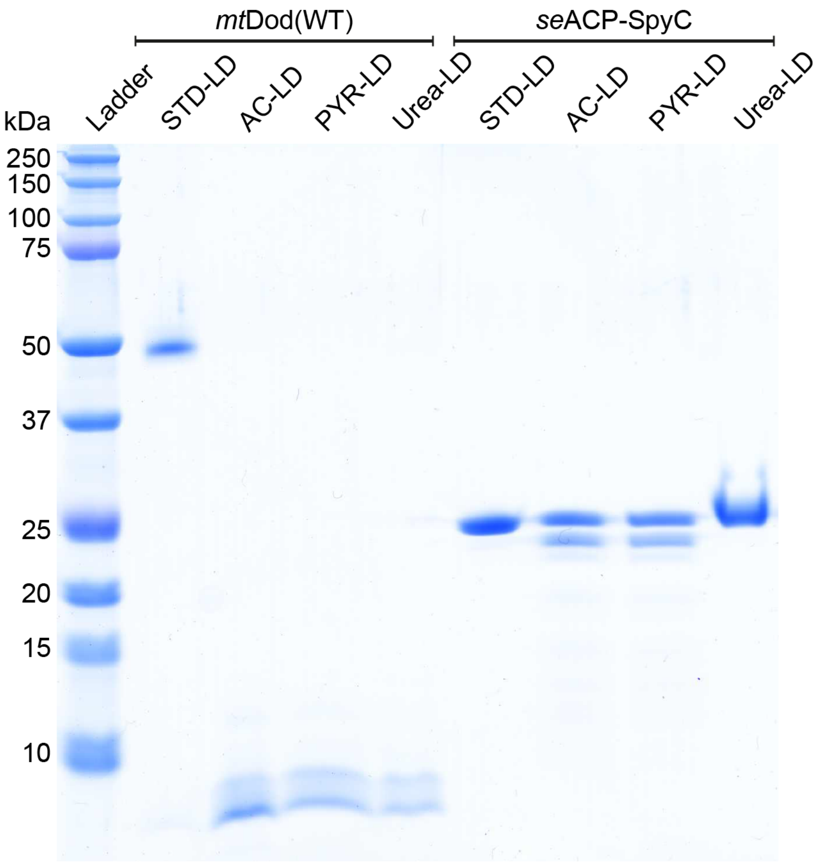


**Fig. S5:** SDS-PAGE gel for comparison of loading buffers. *Mt*Dod(WT) and *se*ACP-SpyC were used. STD-LD: standard loading buffer (denaturing step conditions: pH 6.8, 2.5% SDS). AC-LD: Acetic acid two-component loading buffer (denaturing step conditions: pH < 5.0, 3.3% SDS). PYR-LD: Pyridine two-component loading buffer (denaturing step conditions: pH < 5.0, 3.3% SDS). Urea-LD: standard loading buffer with 8 M urea (denaturing step conditions: pH 6.8, 2.5% SDS, 8 M Urea). All samples were denatured for 5 min at 95 °C. After denaturation, the second component buffer was added to the two-component loading buffer samples (final pH, SDS and glycerol content as in standard loading buffer). All loading buffers, except the standard loading buffer, are able to denature the *mt*Dod(WT) dodecamer, indicated by the monomer band (below 10 kDa). The smearing double bands seem to be an artifact of this specific SDS-PAGE gel, as also the lower molecular weight bands of the ladder show this phenomenon. The gel shows that the *mt*Dod dodecamer tolerates SDS, as long the conditions do not become too acidic. For seACP-SpyC, the acidic denaturing conditions cause the appearance of a band at lower molecular weight. The formation of this band is also temperature dependent. Heat treatment at 60°C with acidic loading buffer did not cause the formation of double bands (see Fig 5).

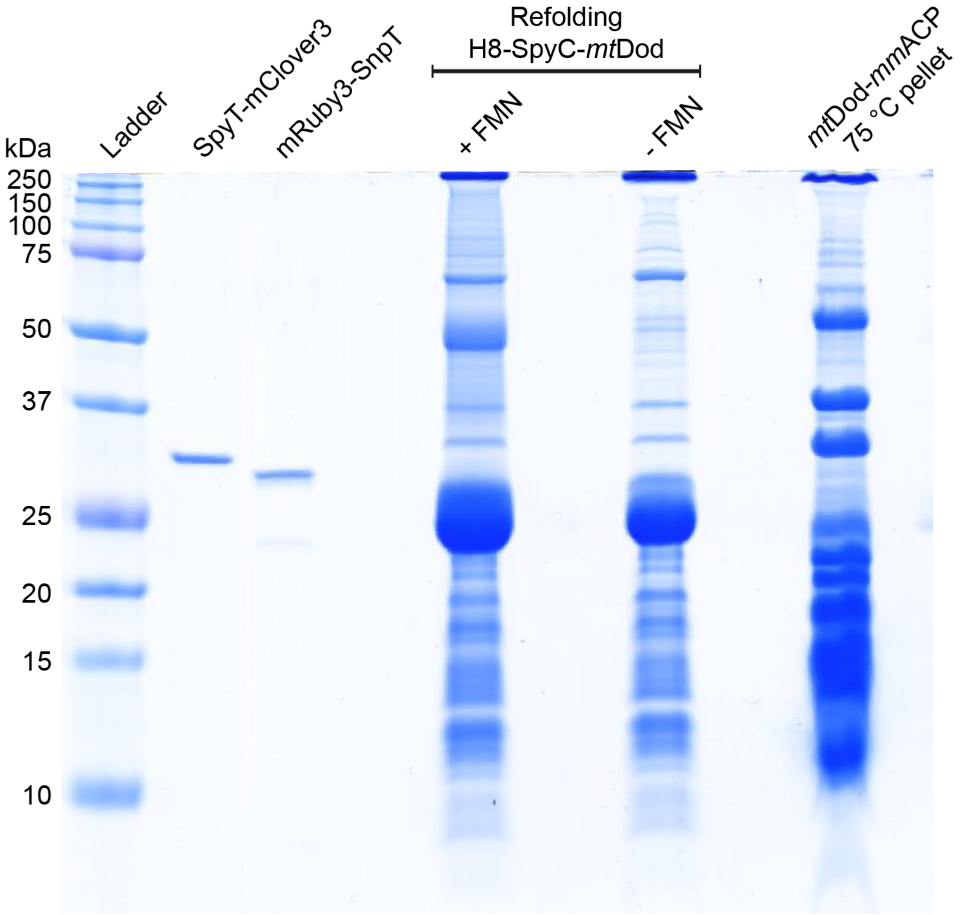


**Fig. S6:** SDS-PAGE gel of *mt*Dod-*mm*ACP precipitating during heat denaturation and other samples. Standard loading buffer was used to preserve dodecameric species. Right sample: Cytosol containing *mt*Dod-*mm*ACP was incubated at 75 °C for 25 min and aggregated proteins were pelleted by centrifugation (15,000 rcf, 10 min). A small amount of the pellet was dissolved in standard SDS loading buffer at 95 °C (about 15 min) and loaded to the gel. In the stained gel, a strong band at high molecular weight (top of the gel) is visible indicating intact dodecamer of *mt*Dod-*mm*ACP. Other samples: SpyT-Clover-H8 and mRuby3-SnpT: pooled SEC fractions (purified by a heat denaturation based protocol). Refolding H8-SpyC-*mt*Dod: first refolding attempt of H8-SpyC-*mt*Dod in our standard buffer, aggregated protein was removed by centrifugation (15,000 rcf, 10 min).

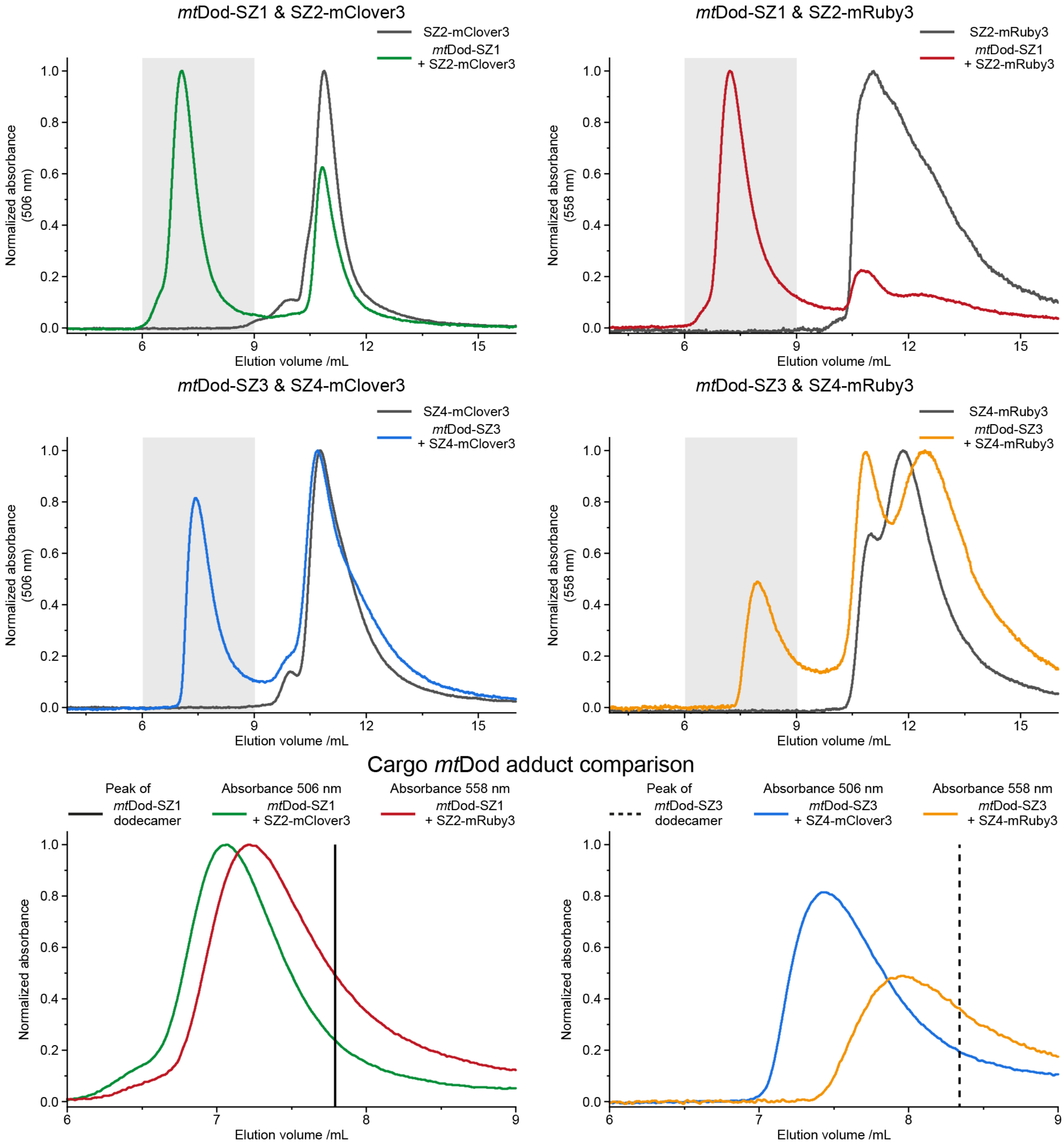


**Fig. S7:** HPLC-SEC chromatograms of *mt*Dod SYNZIP constructs and SYNZIP fluorescence protein constructs. Chromophore absorption of mRuby3 or mClover3 constructs was measured to observe the formation of high molecular weight species. Peaks of the formed adducts (grey highlighted areas) are overlaid in the bottom row. For comparison, elution volumes of the SYNZIP *mt*Dod constructs are shown as a straight (*mt*Dod-SZ1) or dashed (*mt*Dod-SZ3) line (based on absorbance at 280 nm). While for all combinations adduct species were observed, there is a clear difference in dodecamer to cargo composition. The SZ1-SZ2 pair overall performed better than the SZ3-SZ4 pair. For adduct formation, equimolar amounts of carrier and cargo (67 µM) were used and first incubated at 37 °C for 1 h and then on ice for 1 h. Single protein samples were treated the same way. As reaction buffer and running buffer, the standard dodecin buffer (300 mM NaCl, 5 mM $MgCl_2$ and 20 mM Tris-HCl (pH 7.4) was used. Samples were filtered (0.22 µm) and 8 µL were loaded. Runs were performed at 0.3 mL/min and 22 °C. Used column: bioZen$^{TM}$ 1.8 µm SEC-3 (300 × 4.6 mm). Absorbance at 280 nm, 375 nm, 506 nm and 558 nm was measured.

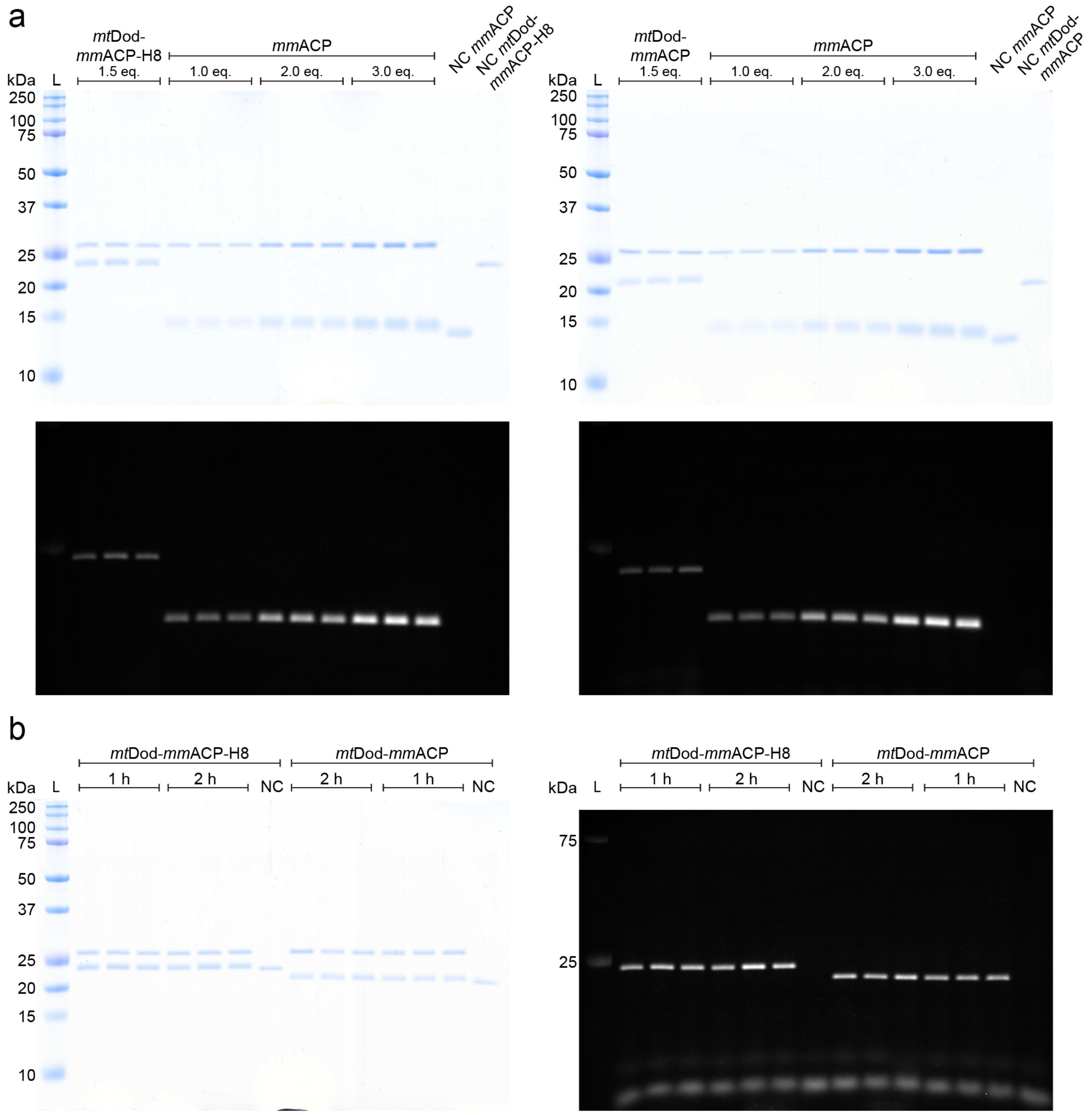


**Fig. S8:** Uncropped SDS-PAGE gels of the modification of *mt*Dod-*mm*ACP and *mt*Dod-*mm*ACP-H8 by Sfp. a) Coomassie stained gels at top, fluorescence images at the bottom. b) (left panel) Coomassie stained gels, (right) fluorescence images. At the bottom of the gel, free fluorophore is observable. In a) no free fluorophore is visible, because of the shorter exposure time used for the images

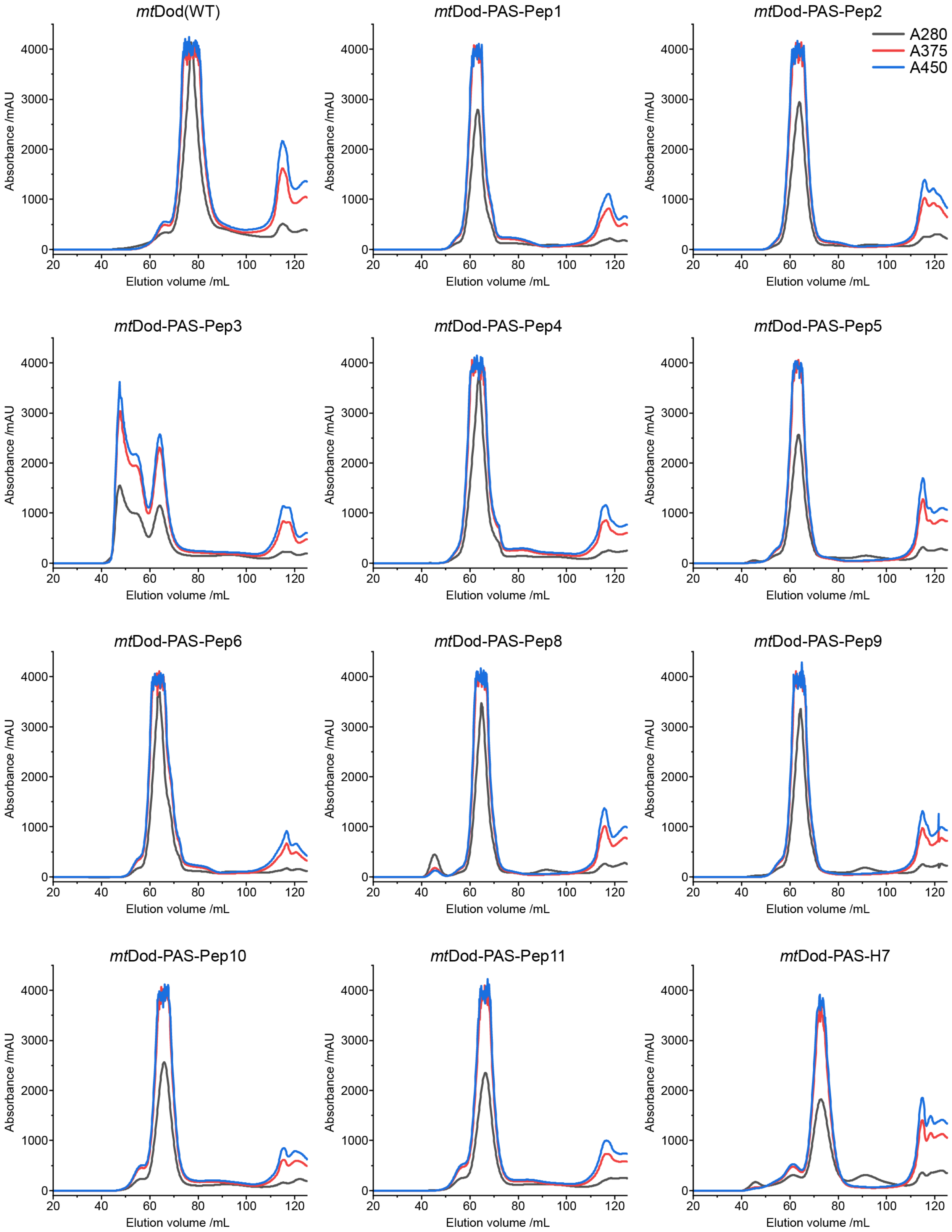


**Fig. S9:** SEC chromatograms of *mt*Dod-PAS-Pep constructs, *mt*Dod-PAS-H7 and *mt*Dod(WT). Used column: Superdex 200 16/60 pg column. Except for *mt*Dod-PAS-Pep3, only minor amounts of aggregates are visible. At about 117 mL, unbound FMN is eluting.

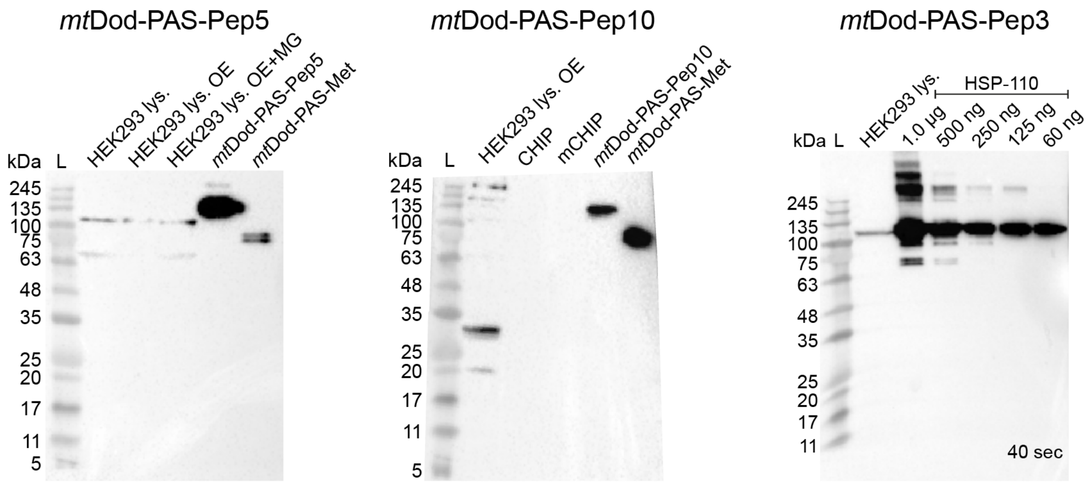


**Fig. S10:** Uncropped western blots of Fig. 8 a. L: Ladder. Lys.: Lysate. OE: protein overexpressing cells. MG: proteasome inhibitor MG-132 added. mCHIP: fragment of CHIP.

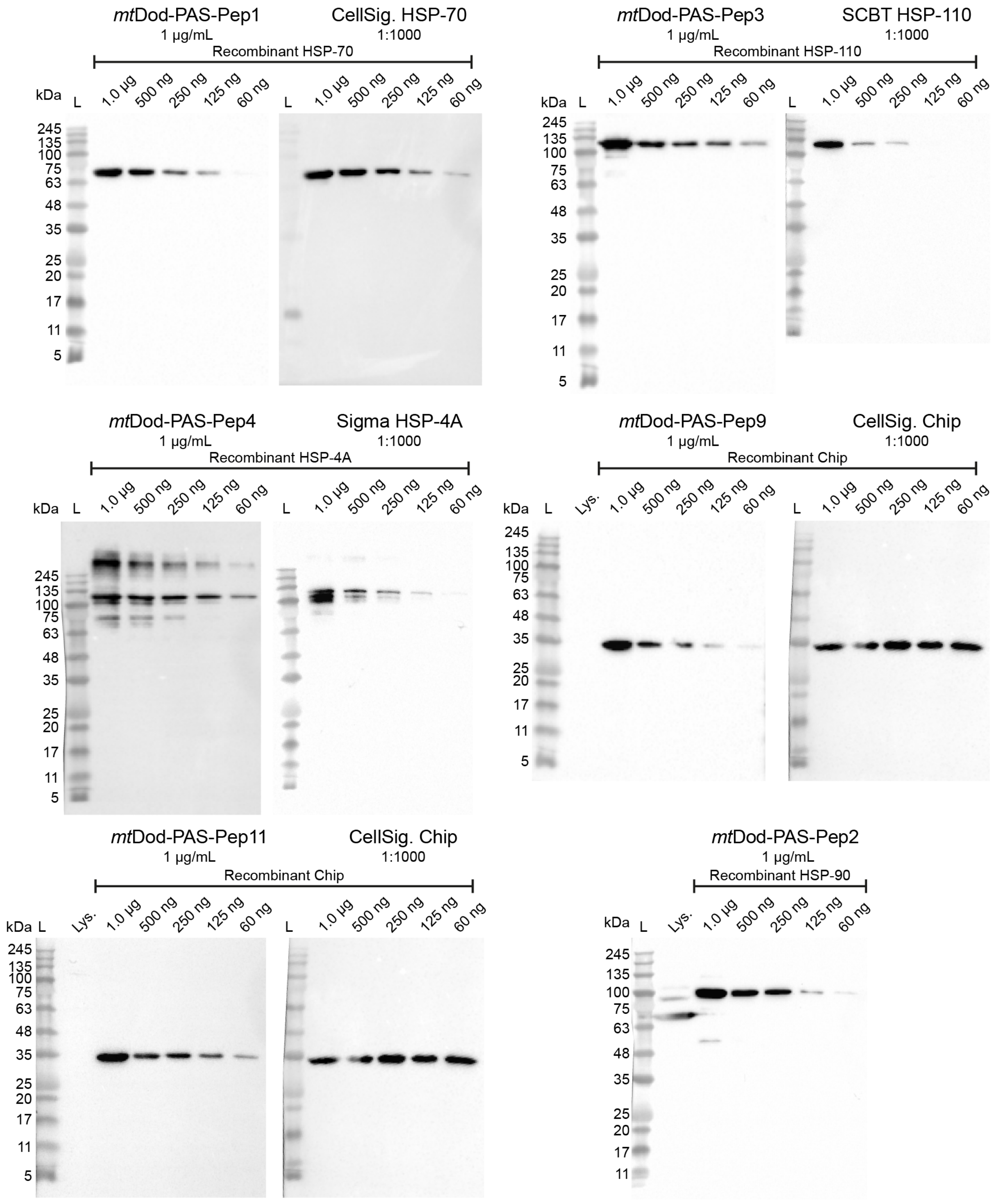


**Fig. S11:** Uncropped western blots of Fig 8 b. L: Ladder. Lys.: Lysate.

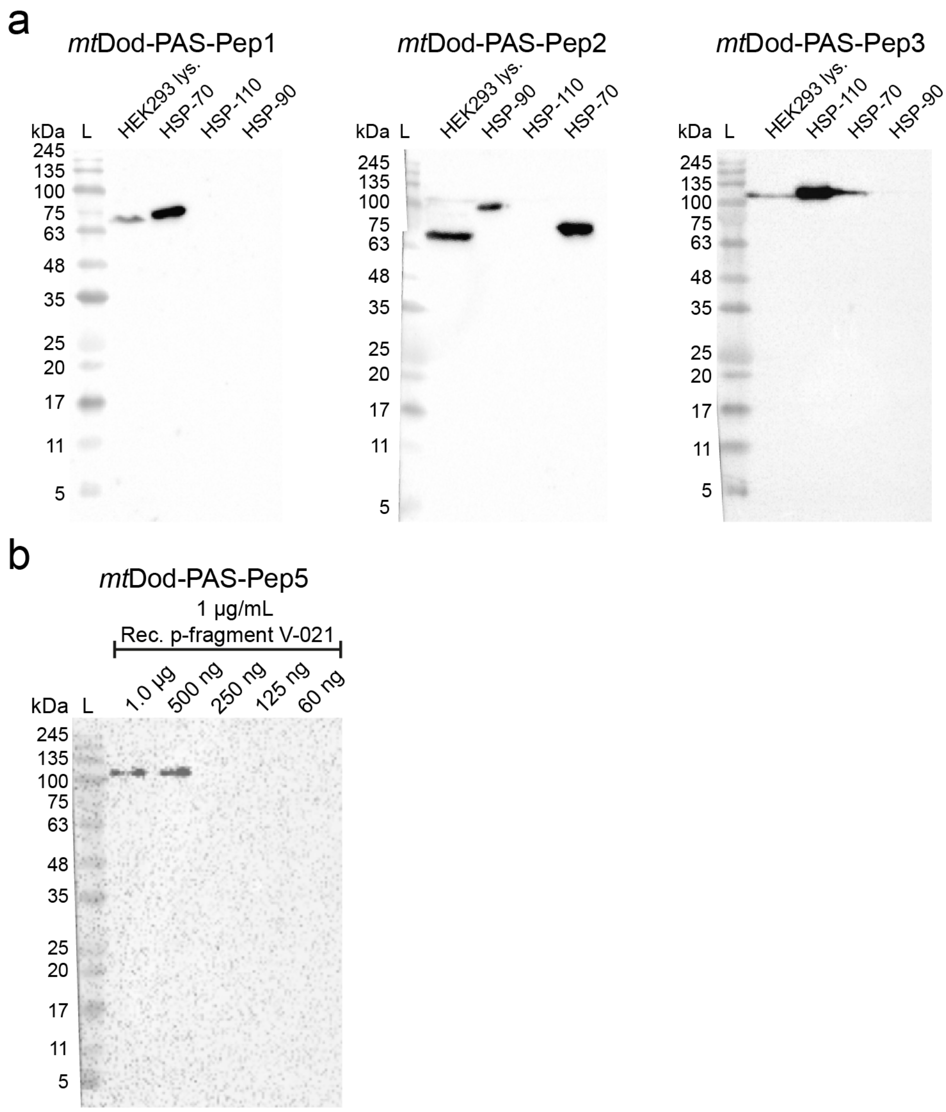


**Fig. S12:** Additional western blots with *mt*Dod-PAS-Pep derived ABs. L: Ladder. a): Comparison of HSP recognizing ABs. ABs derived from *mt*Dod-PAS-Pep2 (designed for HSP-90) recognizes also HSP-70. AB derived from *mt*Dod-PAS-Pep1 and *mt*Dod-PAS-Pep3 recognize only the protein of interest (HSP-90 and HSP-110, respectively). b): Western blot with ABs derived from *mt*Dod-PAS-Pep5. Bands were observed after an exposure time of about 300 sec.

**Table S1**: Table of encoding sequences for all constructs used in this study, except recombinant proteins used in western blotting. All plasmids used for expression were based on a pET22b vector backbone (*lacI* coding sequence, ampicillin resistance, pBR322 origin and f1 origin). Sequences encoding mClover3, mRuby3, SpyC and SnpC were cloned from vectors obtained from Addgene (Plasmid: #74252, pKanCMV-mClover3-mRuby3[2]; Plasmid: #72324, pET28a SpyCatcher-SnoopCatcher[3]). For the polycistronic constructs, spacer DNA sequences between stop codon of the previous gene and the +42 up stream bases of the next gene (based on the pET22b vector and an added restriction side) were designed with EGNAS (version 1158, to minimize secondary structures).[4] These spacer regions were used for cloning (annealing area for In-Fusion® HD Cloning (TaKaRa Bio Europe)). Amino acid sequences of linkers and restriction sites are highlighted in yellow.

| Construct | DNA sequence<br>T7 promotor to T7 terminator<br>(CDS uppercase) | Amino acid sequence<br>(linker) |
|---|---|---|
| ***mt*Dod-peptides** | | |
| *mt*Dod(WT) | cccgcgaaattaatacgactcactataggggaattgtgagcggataacaattcccctctagaaataattttgtttaactttaagaag<br>gagatatacatATGAGCAATCACACCTACCGAGTGATCGAGATCGTCGGGACCTCGCCCGA<br>CGGCGTCGACGCGGCAATCCAGGGCGGTCTGGCCCGAGCTGCGCAGACCATGCGCG<br>CGCTGGACTGGTTCGAAGTACAGTCAATTCGAGGCCACCTGGTCGACGGAGCGGTCG<br>CGCACTTCCAGGTGACTATGAAAGTCGGCTTCCGCCTGGAGGATTCCTGActcgagcacca<br>ccaccaccaccactgagatccggctgctaacaaagcccgaaaggaagctgagttggctgctgccaccgctgagcaataacta<br>gcataaccccttggggcctctaaacgggtcttgaggggttttttg | MSNHTYRVIEIVGTSPDGVDAAIQGGLA<br>RAAQTMRALDWFEVQSIRGHLVDGAVA<br>HFQVTMKVGFRLEDS* |
| *mt*Dod-GSG-Lys | cccgcgaaattaatacgactcactataggggaattgtgagcggataacaattcccctctagaaataattttgtttaactttaagaag<br>gagatatacatATGAGCAATCACACCTACCGAGTGATCGAGATCGTCGGGACCTCGCCCGA<br>CGGCGTCGACGCGGCAATCCAGGGCGGTCTGGCCCGAGCTGCGCAGACCATGCGCG<br>CGCTGGACTGGTTCGAAGTACAGTCAATTCGAGGCCACCTGGTCGACGGAGCGGTCG<br>CGCACTTCCAGGTGACTATGAAAGTCGGCTTCCGCCTGGAGGATTCCCTCGAGGGGG<br>GTGGCGGCAGTGGTGGCGGCGGTAAATGAGGTGACTCTCTGTCTTGGCTGCTGCGTC<br>TGCTGAACtgagatccggctgctaacaaagcccgaaaggaagctgagttggctgctgccaccgctgagcaataactagc<br>ataaccccttggggcctctaaacgggtcttgaggggttttttg | MSNHTYRVIEIVGTSPDGVDAAIQGGLA<br>RAAQTMRALDWFEVQSIRGHLVDGAVA<br>HFQVTMKVGFRLEDSLEGGGGSGGGG<br>K* |
| *mt*Dod-PAS-Met | cccgcgaaattaatacgactcactataggggaattgtgagcggataacaattcccctctagaaataattttgtttaactttaagaag<br>gagatatacatATGAGCAATCACACCTACCGAGTGATCGAGATCGTCGGGACCTCGCCCGA<br>CGGCGTCGACGCGGCAATCCAGGGCGGTCTGGCCCGAGCTGCGCAGACCATGCGCG<br>CGCTGGACTGGTTCGAAGTACAGTCAATTCGAGGCCACCTGGTCGACGGAGCGGTCG<br>CGCACTTCCAGGTGACTATGAAAGTCGGCTTCCGCCTGGAGGATTCCCTCGAGTCTCC<br>AGCTGCGCCTGCTCCGGCAAGCCCTGCGAGCATGTGAggatcccaccaccaccaccaccactgag<br>atccggctgctaacaaagcccgaaaggaagctgagttggctgctgccaccgctgagcaataactagcataaccccttggggcc<br>tctaaacgggtcttgaggggttttttg | MSNHTYRVIEIVGTSPDGVDAAIQGGLA<br>RAAQTMRALDWFEVQSIRGHLVDGAVA<br>HFQVTMKVGFRLEDSLESPAAPAPASPA<br>SM* |
| *mt*Dod-SpyT | cccgcgaaattaatacgactcactataggggaattgtgagcggataacaattcccctctagaaataattttgtttaactttaagaag<br>gagatatacatATGAGCAATCACACCTACCGAGTGATCGAGATCGTCGGGACCTCGCCCGA<br>CGGCGTCGACGCGGCAATCCAGGGCGGTCTGGCCCGAGCTGCGCAGACCATGCGCG<br>CGCTGGACTGGTTCGAAGTACAGTCAATTCGAGGCCACCTGGTCGACGGAGCGGTCG<br>CGCACTTCCAGGTGACTATGAAAGTCGGCTTCCGCCTGGAGGATTCCCTCGAGTCTCC<br>AGCTGCGCCTGCTCCGGCAAGCCCTGCGAGCGGTGGCAGCGGTGCACATATCGTCAT<br>GGTTGATGCGTACAAACCGACCAAATGAggatcccaccaccaccaccaccactgagatccggctgctaac<br>aaagcccgaaaggaagctgagttggctgctgccaccgctgagcaataactagcataaccccttggggcctctaaacgggtctt<br>gaggggttttttg | MSNHTYRVIEIVGTSPDGVDAAIQGGLA<br>RAAQTMRALDWFEVQSIRGHLVDGAVA<br>HFQVTMKVGFRLEDSLESPAAPAPASPA<br>SGGSGAHIVMVDAYKPTK* |
| SpyT-*mt*Dod | cccgcgaaattaatacgactcactataggggaattgtgagcggataacaattcccctctagaaataattttgtttaactttaagaag<br>gagatatacatATGGCACATATCGTCATGGTTGATGCGTACAAACCGACCAAAGGTGGCAGC<br>GGTTCTCCAGCTGCGCCTGCTCCGGCAAGCCCTGCGAGCAGCAATCACACCTACCGA<br>GTGATCGAGATCGTCGGGACCTCGCCCGACGGCGTCGACGCGGCAATCCAGGGCGG<br>TCTGGCCCGAGCTGCGCAGACCATGCGCGCGCTGGACTGGTTCGAAGTACAGTCAAT<br>TCGAGGCCACCTGGTCGACGGAGCGGTCGCGCACTTCCAGGTGACTATGAAAGTCGG<br>CTTCCGCCTGGAGGATTCCTGActcgagcaccaccaccaccaccactgagatccggctgctaacaaagcccg<br>aaaggaagctgagttggctgctgccaccgctgagcaataactagcataaccccttggggcctctaaacgggtcttgaggggttttt<br>tg | MAHIVMVDAYKPTKGGSGSPAAPAPAS<br>PASSNHTYRVIEIVGTSPDGVDAAIQGGL<br>ARAAQTMRALDWFEVQSIRGHLVDGAV<br>AHFQVTMKVGFRLEDS* |
| *mt*Dod-PAS2-SpyT | cccgcgaaattaatacgactcactataggggaattgtgagcggataacaattcccctctagaaataattttgtttaactttaagaag<br>gagatatacatATGAGCAATCACACCTACCGAGTGATCGAGATCGTCGGGACCTCGCCCGA<br>CGGCGTCGACGCGGCAATCCAGGGCGGTCTGGCCCGAGCTGCGCAGACCATGCGCG<br>CGCTGGACTGGTTCGAAGTACAGTCAATTCGAGGCCACCTGGTCGACGGAGCGGTCG<br>CGCACTTCCAGGTGACTATGAAAGTCGGCTTCCGCCTGGAGGATTCCCTCGAGTCTCC<br>AGCTGCGCCTGCTCCGGCAAGCCCTGCGTCTCCGGCACCGTCTGCGCCAGCTGCATC<br>TCCAGCAGCGGGTGGCAGCGGTGCACATATCGTCATGGTTGATGCGTACAAACCGAC<br>CAAATGAggatcccaccaccaccaccaccactgagatccggctgctaacaaagcccgaaaggaagctgagttggctgct<br>gccaccgctgagcaataactagcataaccccttggggcctctaaacgggtcttgaggggttttttg | MSNHTYRVIEIVGTSPDGVDAAIQGGLA<br>RAAQTMRALDWFEVQSIRGHLVDGAVA<br>HFQVTMKVGFRLEDSLESPAAPAPASPA<br>SPAPSAPAASPAAGGSGAHIVMVDAYKP<br>TK* |
| SpyT-PAS2-*mt*Dod | cccgcgaaattaatacgactcactataggggaattgtgagcggataacaattcccctctagaaataattttgtttaactttaagaag<br>gagatatacatATGGCACATATCGTCATGGTTGATGCGTACAAACCGACCAAAGGTGGCAGC<br>GGTTCTCCAGCTGCGCCTGCTCCGGCAAGCCCTGCGTCTCCGGCACCGTCTGCGCCA<br>GCTGCATCTCCAGCAGCGAGCAATCACACCTACCGAGTGATCGAGATCGTCGGGACCT<br>CGCCCGACGGCGTCGACGCGGCAATCCAGGGCGGTCTGGCCCGAGCTGCGCAGACC<br>ATGCGCGCGCTGGACTGGTTCGAAGTACAGTCAATTCGAGGCCACCTGGTCGACGGA<br>GCGGTCGCGCACTTCCAGGTGACTATGAAAGTCGGCTTCCGCCTGGAGGATTCCTGAc<br>tcgagcaccaccaccaccaccactgagatccggctgctaacaaagcccgaaaggaagctgagttggctgctgccaccgctga<br>gcaataactagcataaccccttggggcctctaaacgggtcttgaggggttttttg | MAHIVMVDAYKPTKGGSGSPAAPAPAS<br>PASPAPSAPAASPAASNHTYRVIEIVGTS<br>PDGVDAAIQGGLARAAQTMRALDWFEV<br>QSIRGHLVDGAVAHFQVTMKVGFRLED<br>S* |

| | | |
|---|---|---|
| SpyT-*mt*Dod-SnpT | cccgcgaaattaatacgactcactataggggaattgtgagcggataacaattcccctctagaaataattttgtttaactttaagaag<br>gagatatacatATGGCACATATCGTCATGGTTGATGCGTACAAACCGACCAAAGGTGGCAGC<br>GGTTCTCCAGCTGCGCCTGCTCCGGCAAGCCCTGCGAGCAGCAATCACACCTACCGA<br>GTGATCGAGATCGTCGGGACCTCGCCCGACGGCGTCGACGCGGCAATCCAGGGCGG<br>TCTGGCCCGAGCTGCGCAGACCATGCGCGCGCTGGACTGGTTCGAAGTACAGTCAAT<br>TCGAGGCCACCTGGTCGACGGAGCGGTCGCGCACTTCCAGGTGACTATGAAAGTCGG<br>CTTCCGCCTGGAGGATTCCTCTCCAGCTGCGCCTGCTCCGGCAAGCCCTGCGAGCGG<br>TGGCAGCGGTGGCAAACTGGGCGATATTGAATTTATTAAAGTGAACAAAGGCTATTGAg<br>gatcccaccaccaccaccaccactgagatccggctgctaacaaagcccgaaaggaagctgagttggctgctgccaccgctga<br>gcaataactagcataaccccttggggcctctaaacgggtcttgaggggttttttg | MAHIVMVDAYKPTKGGSGSPAAPAPAS<br>PASSNHTYRVIEIVGTSPDGVDAAIQGGL<br>ARAAQTMRALDWFEVQSIRGHLVDGAV<br>AHFQVTMKVGFRLEDSSPAAPAPASPA<br>SGGSGGKLGDIEFIKVNKGY* |
| SnpT-*mt*Dod-SpyT | cccgcgaaattaatacgactcactataggggaattgtgagcggataacaattcccctctagaaataattttgtttaactttaagaag<br>gagatatacatATGGGCAAACTGGGCGATATTGAATTTATTAAAGTGAACAAAGGCTATGGT<br>GGCAGCGGTTCTCCAGCTGCGCCTGCTCCGGCAAGCCCTGCGAGCAGCAATCACACC<br>TACCGAGTGATCGAGATCGTCGGGACCTCGCCCGACGGCGTCGACGCGGCAATCCAG<br>GGCGGTCTGGCCCGAGCTGCGCAGACCATGCGCGCGCTGGACTGGTTCGAAGTACAG<br>TCAATTCGAGGCCACCTGGTCGACGGAGCGGTCGCGCACTTCCAGGTGACTATGAAA<br>GTCGGCTTCCGCCTGGAGGATTCCCTCGAGTCTCCAGCTGCGCCTGCTCCGGCAAGC<br>CCTGCGAGCGGTGGCAGCGGTGCACATATCGTCATGGTTGATGCGTACAAACCGACC<br>AAATGAggatcccaccaccaccaccaccactgagatccggctgctaacaaagcccgaaaggaagctgagttggctgctg<br>ccaccgctgagcaataactagcataaccccttggggcctctaaacgggtcttgaggggttttttg | MGKLGDIEFIKVNKGYGGSGSPAAPAPA<br>SPASSNHTYRVIEIVGTSPDGVDAAIQGG<br>LARAAQTMRALDWFEVQSIRGHLVDGA<br>VAHFQVTMKVGFRLEDSLESPAAPAPAS<br>PASGGSGAHIVMVDAYKPTK* |
| *mt*Dod-PAS-StrepII | cccgcgaaattaatacgactcactataggggaattgtgagcggataacaattcccctctagaaataattttgtttaactttaagaag<br>gagatatacatATGAGCAATCACACCTACCGAGTGATCGAGATCGTCGGGACCTCGCCCGA<br>CGGCGTCGACGCGGCAATCCAGGGCGGTCTGGCCCGAGCTGCGCAGACCATGCGCG<br>CGCTGGACTGGTTCGAAGTACAGTCAATTCGAGGCCACCTGGTCGACGGAGCGGTCG<br>CGCACTTCCAGGTGACTATGAAAGTCGGCTTCCGCCTGGAGGATTCCCTCGAGTCTCC<br>AGCTGCGCCTGCTCCGGCAAGCCCTGCGAGCTGGAGCCACCCGCAGTTCGAAAAATG<br>Agatccggctgctaacaaagcccgaaaggaagctgagttggctgctgccaccgctgagcaataactagcataaccccttggg<br>gcctctaaacgggtcttgaggggttttttg | MSNHTYRVIEIVGTSPDGVDAAIQGGLA<br>RAAQTMRALDWFEVQSIRGHLVDGAVA<br>HFQVTMKVGFRLEDSLESPAAPAPASPA<br>SWSHPQFEK* |
| *mt*Dod-PAS-H7 | cccgcgaaattaatacgactcactataggggaattgtgagcggataacaattcccctctagaaataattttgtttaactttaagaag<br>gagatatacatATGAGCAATCACACCTACCGAGTGATCGAGATCGTCGGGACCTCGCCCGA<br>CGGCGTCGACGCGGCAATCCAGGGCGGTCTGGCCCGAGCTGCGCAGACCATGCGCG<br>CGCTGGACTGGTTCGAAGTACAGTCAATTCGAGGCCACCTGGTCGACGGAGCGGTCG<br>CGCACTTCCAGGTGACTATGAAAGTCGGCTTCCGCCTGGAGGATTCCCTCGAGTCTCC<br>AGCTGCGCCTGCTCCGGCAAGCCCTGCGAGCCACCACCACCACCACCACCACTGAgat<br>ccggctgctaacaaagcccgaaaggaagctgagttggctgctgccaccgctgagcaataactagcataaccccttggggcctc<br>taaacgggtcttgaggggttttttg | MSNHTYRVIEIVGTSPDGVDAAIQGGLA<br>RAAQTMRALDWFEVQSIRGHLVDGAVA<br>HFQVTMKVGFRLEDSLESPAAPAPASPA<br>SHHHHHHH* |
| *mt*Dod-PAS-Pep1 | cccgcgaaattaatacgactcactataggggaattgtgagcggataacaattcccctctagaaataattttgtttaactttaagaag<br>gagatatacatATGAGCAATCACACCTACCGAGTGATCGAGATCGTCGGGACCTCGCCCGA<br>CGGCGTCGACGCGGCAATCCAGGGCGGTCTGGCCCGAGCTGCGCAGACCATGCGCG<br>CGCTGGACTGGTTCGAAGTACAGTCAATTCGAGGCCACCTGGTCGACGGAGCGGTCG<br>CGCACTTCCAGGTGACTATGAAAGTCGGCTTCCGCCTGGAGGATTCCCTCGAGTCTCC<br>AGCTGCGCCTGCTCCGGCAAGCCCTGCGAGCCCGAAAGGTGGCAGCGGTTCTGGCC<br>CGACCATTGAAGAAGTGGATTGAgatccggctgctaacaaagcccgaaaggaagctgagttggctgctgccac<br>cgctgagcaataactagcataaccccttggggcctctaaacgggtcttgaggggttttttg | MSNHTYRVIEIVGTSPDGVDAAIQGGLA<br>RAAQTMRALDWFEVQSIRGHLVDGAVA<br>HFQVTMKVGFRLEDSLESPAAPAPASPA<br>SPKGGSGSGPTIEEVD* |
| *mt*Dod-PAS-Pep2 | cccgcgaaattaatacgactcactataggggaattgtgagcggataacaattcccctctagaaataattttgtttaactttaagaag<br>gagatatacatATGAGCAATCACACCTACCGAGTGATCGAGATCGTCGGGACCTCGCCCGA<br>CGGCGTCGACGCGGCAATCCAGGGCGGTCTGGCCCGAGCTGCGCAGACCATGCGCG<br>CGCTGGACTGGTTCGAAGTACAGTCAATTCGAGGCCACCTGGTCGACGGAGCGGTCG<br>CGCACTTCCAGGTGACTATGAAAGTCGGCTTCCGCCTGGAGGATTCCCTCGAGTCTCC<br>AGCTGCGCCTGCTCCGGCAAGCCCTGCGAGCCCGCTGGAAGGCGATGACGATACCAG<br>CCGCATGGAAGAAGTGGATTGAgatccggctgctaacaaagcccgaaaggaagctgagttggctgctgccacc<br>gctgagcaataactagcataaccccttggggcctctaaacgggtcttgaggggttttttg | MSNHTYRVIEIVGTSPDGVDAAIQGGLA<br>RAAQTMRALDWFEVQSIRGHLVDGAVA<br>HFQVTMKVGFRLEDSLESPAAPAPASPA<br>SPLEGDDDTSRMEEVD* |
| *mt*Dod-PAS-Pep3 | cccgcgaaattaatacgactcactataggggaattgtgagcggataacaattcccctctagaaataattttgtttaactttaagaag<br>gagatatacatATGAGCAATCACACCTACCGAGTGATCGAGATCGTCGGGACCTCGCCCGA<br>CGGCGTCGACGCGGCAATCCAGGGCGGTCTGGCCCGAGCTGCGCAGACCATGCGCG<br>CGCTGGACTGGTTCGAAGTACAGTCAATTCGAGGCCACCTGGTCGACGGAGCGGTCG<br>CGCACTTCCAGGTGACTATGAAAGTCGGCTTCCGCCTGGAGGATTCCCTCGAGTCTCC<br>AGCTGCGCCTGCTCCGGCAAGCCCTGCGAGCGAATGCTATCCGAACGAGAAGAACAG<br>CGTGAACATGGATCTGGATTGAgatccggctgctaacaaagcccgaaaggaagctgagttggctgctgccacc<br>gctgagcaataactagcataaccccttggggcctctaaacgggtcttgaggggttttttg | MSNHTYRVIEIVGTSPDGVDAAIQGGLA<br>RAAQTMRALDWFEVQSIRGHLVDGAVA<br>HFQVTMKVGFRLEDSLESPAAPAPASPA<br>SECYPNEKNSVNMDLD* |
| *mt*Dod-PAS-Pep4 | cccgcgaaattaatacgactcactataggggaattgtgagcggataacaattcccctctagaaataattttgtttaactttaagaag<br>gagatatacatATGAGCAATCACACCTACCGAGTGATCGAGATCGTCGGGACCTCGCCCGA<br>CGGCGTCGACGCGGCAATCCAGGGCGGTCTGGCCCGAGCTGCGCAGACCATGCGCG<br>CGCTGGACTGGTTCGAAGTACAGTCAATTCGAGGCCACCTGGTCGACGGAGCGGTCG<br>CGCACTTCCAGGTGACTATGAAAGTCGGCTTCCGCCTGGAGGATTCCCTCGAGTCTCC<br>AGCTGCGCCTGCTCCGGCAAGCCCTGCGAGCGTGCCGAGCGACAGCGATAAGAAACT<br>GCCGGAAATGGATATTGATTGAgatccggctgctaacaaagcccgaaaggaagctgagttggctgctgccacc<br>gctgagcaataactagcataaccccttggggcctctaaacgggtcttgaggggttttttg | MSNHTYRVIEIVGTSPDGVDAAIQGGLA<br>RAAQTMRALDWFEVQSIRGHLVDGAVA<br>HFQVTMKVGFRLEDSLESPAAPAPASPA<br>SVPSDSDKKLPEMDID* |
| *mt*Dod-PAS-Pep5 | cccgcgaaattaatacgactcactataggggaattgtgagcggataacaattcccctctagaaataattttgtttaactttaagaag<br>gagatatacatATGAGCAATCACACCTACCGAGTGATCGAGATCGTCGGGACCTCGCCCGA<br>CGGCGTCGACGCGGCAATCCAGGGCGGTCTGGCCCGAGCTGCGCAGACCATGCGCG<br>CGCTGGACTGGTTCGAAGTACAGTCAATTCGAGGCCACCTGGTCGACGGAGCGGTCG<br>CGCACTTCCAGGTGACTATGAAAGTCGGCTTCCGCCTGGAGGATTCCCTCGAGTCTCC<br>AGCTGCGCCTGCTCCGGCAAGCCCTGCGAGCGATAGCAGCCAGCATACCAAGAGCTC<br>TGGCGAAATGGAAGTGGATTGAgatccggctgctaacaaagcccgaaaggaagctgagttggctgctgccacc<br>gctgagcaataactagcataaccccttggggcctctaaacgggtcttgaggggttttttg | MSNHTYRVIEIVGTSPDGVDAAIQGGLA<br>RAAQTMRALDWFEVQSIRGHLVDGAVA<br>HFQVTMKVGFRLEDSLESPAAPAPASPA<br>SDSSQHTKSSGEMEVD* |
| *mt*Dod-PAS-Pep6 | cccgcgaaattaatacgactcactataggggaattgtgagcggataacaattcccctctagaaataattttgtttaactttaagaag<br>gagatatacatATGAGCAATCACACCTACCGAGTGATCGAGATCGTCGGGACCTCGCCCGA<br>CGGCGTCGACGCGGCAATCCAGGGCGGTCTGGCCCGAGCTGCGCAGACCATGCGCG<br>CGCTGGACTGGTTCGAAGTACAGTCAATTCGAGGCCACCTGGTCGACGGAGCGGTCG<br>CGCACTTCCAGGTGACTATGAAAGTCGGCTTCCGCCTGGAGGATTCCCTCGAGTCTCC<br>AGCTGCGCCTGCTCCGGCAAGCCCTGCGAGCGAACAGAGCACCGGCCAGAAACGCC<br>CGCTGAAGAACGATGAACTGTGAgatccggctgctaacaaagcccgaaaggaagctgagttggctgctgccac<br>cgctgagcaataactagcataaccccttggggcctctaaacgggtcttgaggggttttttg | MSNHTYRVIEIVGTSPDGVDAAIQGGLA<br>RAAQTMRALDWFEVQSIRGHLVDGAVA<br>HFQVTMKVGFRLEDSLESPAAPAPASPA<br>SEQSTGQKRPLKNDEL* |
| *mt*Dod-PAS-Pep7 | cccgcgaaattaatacgactcactataggggaattgtgagcggataacaattcccctctagaaataattttgtttaactttaagaag<br>gagatatacatATGAGCAATCACACCTACCGAGTGATCGAGATCGTCGGGACCTCGCCCGA<br>CGGCGTCGACGCGGCAATCCAGGGCGGTCTGGCCCGAGCTGCGCAGACCATGCGCG<br>CGCTGGACTGGTTCGAAGTACAGTCAATTCGAGGCCACCTGGTCGACGGAGCGGTCG<br>CGCACTTCCAGGTGACTATGAAAGTCGGCTTCCGCCTGGAGGATTCCCTCGAGTCTCC<br>AGCTGCGCCTGCTCCGGCAAGCCCTGCGAGCGCGCTGATGGTGTATCGCTGCGCGCC<br>GCCGCGCAGCAGCCAGTTTTGAgatccggctgctaacaaagcccgaaaggaagctgagttggctgctgccacc<br>gctgagcaataactagcataaccccttggggcctctaaacgggtcttgaggggttttttg | MSNHTYRVIEIVGTSPDGVDAAIQGGLA<br>RAAQTMRALDWFEVQSIRGHLVDGAVA<br>HFQVTMKVGFRLEDSLESPAAPAPASPA<br>SALMVYRCAPPRSSQF* |

| | | |
|---|---|---|
| *mt*Dod-PAS-Pep8 | cccgcgaaattaatacgactcactataggggaattgtgagcggataacaattcccctctagaaataattttgtttaactttaagaag<br>gagatatacatATGAGCAATCACACCTACCGAGTGATCGAGATCGTCGGGACCTCGCCCGA<br>CGGCGTCGACGCGGCAATCCAGGGCGGTCTGGCCCGAGCTGCGCAGACCATGCGCG<br>CGCTGGACTGGTTCGAAGTACAGTCAATTCGAGGCCACCTGGTCGACGGAGCGGTCG<br>CGCACTTCCAGGTGACTATGAAAGTCGGCTTCCGCCTGGAGGATTCCCTCGAGTCTCC<br>AGCTGCGCCTGCTCCGGCAAGCCCTGCGAGCCTGGTGACCGGCGAAAGCCTGGAACA<br>GCTGCGCCGCGGCCTGGCGTGAgatccggctgctaacaaagcccgaaaggaagctgagttggctgctgccac<br>cgctgagcaataactagcataaccccttggggcctctaaacgggtcttgaggggttttttg | MSNHTYRVIEIVGTSPDGVDAAIQGGLA<br>RAAQTMRALDWFEVQSIRGHLVDGAVA<br>HFQVTMKVGFRLEDSLESPAAPAPASPA<br>SLVTGESLEQLRRGLA* |
| *mt*Dod-PAS-Pep9 | cccgcgaaattaatacgactcactataggggaattgtgagcggataacaattcccctctagaaataattttgtttaactttaagaag<br>gagatatacatATGAGCAATCACACCTACCGAGTGATCGAGATCGTCGGGACCTCGCCCGA<br>CGGCGTCGACGCGGCAATCCAGGGCGGTCTGGCCCGAGCTGCGCAGACCATGCGCG<br>CGCTGGACTGGTTCGAAGTACAGTCAATTCGAGGCCACCTGGTCGACGGAGCGGTCG<br>CGCACTTCCAGGTGACTATGAAAGTCGGCTTCCGCCTGGAGGATTCCCTCGAGTCTCC<br>AGCTGCGCCTGCTCCGGCAAGCCCTGCGAGCATGAAAGGCAAAGAAGAGAAAGAAGG<br>CGGCGCGCGCCTGGGCGCGTGAgatccggctgctaacaaagcccgaaaggaagctgagttggctgctgcca<br>ccgctgagcaataactagcataaccccttggggcctctaaacgggtcttgaggggttttttg | MSNHTYRVIEIVGTSPDGVDAAIQGGLA<br>RAAQTMRALDWFEVQSIRGHLVDGAVA<br>HFQVTMKVGFRLEDSLESPAAPAPASPA<br>SMKGKEEKEGGARLGA* |
| *mt*Dod-PAS-Pep10 | cccgcgaaattaatacgactcactataggggaattgtgagcggataacaattcccctctagaaataattttgtttaactttaagaag<br>gagatatacatATGAGCAATCACACCTACCGAGTGATCGAGATCGTCGGGACCTCGCCCGA<br>CGGCGTCGACGCGGCAATCCAGGGCGGTCTGGCCCGAGCTGCGCAGACCATGCGCG<br>CGCTGGACTGGTTCGAAGTACAGTCAATTCGAGGCCACCTGGTCGACGGAGCGGTCG<br>CGCACTTCCAGGTGACTATGAAAGTCGGCTTCCGCCTGGAGGATTCCCTCGAGTCTCC<br>AGCTGCGCCTGCTCCGGCAAGCCCTGCGAGCGAAGAACGCCGCATTCATCAGGAAAG<br>CGAATGAgatccggctgctaacaaagcccgaaaggaagctgagttggctgctgccaccgctgagcaataactagcataa<br>ccccttggggcctctaaacgggtcttgaggggttttttg | MSNHTYRVIEIVGTSPDGVDAAIQGGLA<br>RAAQTMRALDWFEVQSIRGHLVDGAVA<br>HFQVTMKVGFRLEDSLESPAAPAPASPA<br>SEERRIHQESE* |
| *mt*Dod-PAS-Pep11 | cccgcgaaattaatacgactcactataggggaattgtgagcggataacaattcccctctagaaataattttgtttaactttaagaag<br>gagatatacatATGAGCAATCACACCTACCGAGTGATCGAGATCGTCGGGACCTCGCCCGA<br>CGGCGTCGACGCGGCAATCCAGGGCGGTCTGGCCCGAGCTGCGCAGACCATGCGCG<br>CGCTGGACTGGTTCGAAGTACAGTCAATTCGAGGCCACCTGGTCGACGGAGCGGTCG<br>CGCACTTCCAGGTGACTATGAAAGTCGGCTTCCGCCTGGAGGATTCCCTCGAGTCTCC<br>AGCTGCGCCTGCTCCGGCAAGCCCTGCGAGCAACCATGAAGGCGATGAAGATGATAG<br>CCATTGAgatccggctgctaacaaagcccgaaaggaagctgagttggctgctgccaccgctgagcaataactagcataac<br>cccttggggcctctaaacgggtcttgaggggttttttg | MSNHTYRVIEIVGTSPDGVDAAIQGGLA<br>RAAQTMRALDWFEVQSIRGHLVDGAVA<br>HFQVTMKVGFRLEDSLESPAAPAPASPA<br>SNHEGDEDDSH* |
| ***poly-cistronic constructs*** | DNA sequence<br>T7 promotor to T7 terminator<br>(CDS uppercase, spacer sequence) | Amino acid sequence<br>(Linker) |
| bici<br>(*mt*Dod(WT);<br>*mt*Dod-PAS-StrepII) | cccgcgaaattaatacgactcactataggggaattgtgagcggataacaattcccctctagaaataattttgtttaactttaagaag<br>gagatatacatATGAGCAATCACACCTACCGAGTGATCGAGATCGTCGGGACCTCGCCCGA<br>CGGCGTCGACGCGGCAATCCAGGGCGGTCTGGCCCGAGCTGCGCAGACCATGCGCG<br>CGCTGGACTGGTTCGAAGTACAGTCAATTCGAGGCCACCTGGTCGACGGAGCGGTCG<br>CGCACTTCCAGGTGACTATGAAAGTCGGCTTCCGCCTGGAGGATTCCTGAcggtttgaaacc<br>ttccgaggagaattcaataattttgtttaactttaagaaggagatatacatATGAGCAATCACACCTACCGAGTGA<br>TCGAGATCGTCGGGACCTCGCCCGACGGCGTCGACGCGGCAATCCAGGGCGGTCTG<br>GCCCGAGCTGCGCAGACCATGCGCGCGCTGGACTGGTTCGAAGTACAGTCAATTCGA<br>GGCCACCTGGTCGACGGAGCGGTCGCGCACTTCCAGGTGACTATGAAAGTCGGCTTC<br>CGCCTGGAGGATTCCCTCGAGTCTCCAGCTGCGCCTGCTCCGGCAAGCCCTGCGAGC<br>TGGAGCCACCCGCAGTTCGAAAAATGAgaatcgcgattcaccatggtgggatccgatccggctgctaacaaa<br>gcccgaaaggaagctgagttggctgctgccaccgctgagcaataactagcataaccccttggggcctctaaacgggtcttgag<br>gggttttttg | MSNHTYRVIEIVGTSPDGVDAAIQGGLA<br>RAAQTMRALDWFEVQSIRGHLVDGAVA<br>HFQVTMKVGFRLEDS*<br>&<br>MSNHTYRVIEIVGTSPDGVDAAIQGGLA<br>RAAQTMRALDWFEVQSIRGHLVDGAVA<br>HFQVTMKVGFRLEDSLESPAAPAPASPA<br>SWSHPQFEK* |
| bici<br>(*mt*Dod-PAS-StrepII;<br>*mt*Dod(WT)) | cccgcgaaattaatacgactcactataggggaattgtgagcggataacaattcccctctagaaataattttgtttaactttaagaag<br>gagatatacatATGAGCAATCACACCTACCGAGTGATCGAGATCGTCGGGACCTCGCCCGA<br>CGGCGTCGACGCGGCAATCCAGGGCGGTCTGGCCCGAGCTGCGCAGACCATGCGCG<br>CGCTGGACTGGTTCGAAGTACAGTCAATTCGAGGCCACCTGGTCGACGGAGCGGTCG<br>CGCACTTCCAGGTGACTATGAAAGTCGGCTTCCGCCTGGAGGATTCCCTCGAGTCTCC<br>AGCTGCGCCTGCTCCGGCAAGCCCTGCGAGCTGGAGCCACCCGCAGTTCGAAAAATG<br>AcggtttgaaaccttccgaggagaattcaataattttgtttaactttaagaaggagatatacatATGAGCAATCACACCT<br>ACCGAGTGATCGAGATCGTCGGGACCTCGCCCGACGGCGTCGACGCGGCAATCCAGG<br>GCGGTCTGGCCCGAGCTGCGCAGACCATGCGCGCGCTGGACTGGTTCGAAGTACAGT<br>CAATTCGAGGCCACCTGGTCGACGGAGCGGTCGCGCACTTCCAGGTGACTATGAAAG<br>TCGGCTTCCGCCTGGAGGATTCCTGAgaatcgcgattcaccatggtgggatccgatccggctgctaacaaag<br>cccgaaaggaagctgagttggctgctgccaccgctgagcaataactagcataaccccttggggcctctaaacgggtcttgagg<br>ggttttttg | MSNHTYRVIEIVGTSPDGVDAAIQGGLA<br>RAAQTMRALDWFEVQSIRGHLVDGAVA<br>HFQVTMKVGFRLEDSLESPAAPAPASPA<br>SWSHPQFEK*<br>&<br>MSNHTYRVIEIVGTSPDGVDAAIQGGLA<br>RAAQTMRALDWFEVQSIRGHLVDGAVA<br>HFQVTMKVGFRLEDS* |
| tricis<br>(*mt*Dod(WT);<br>*mt*Dod(WT);<br>*mt*Dod-PAS-StrepII) | cccgcgaaattaatacgactcactataggggaattgtgagcggataacaattcccctctagaaataattttgtttaactttaagaag<br>gagatatacatATGAGCAATCACACCTACCGAGTGATCGAGATCGTCGGGACCTCGCCCGA<br>CGGCGTCGACGCGGCAATCCAGGGCGGTCTGGCCCGAGCTGCGCAGACCATGCGCG<br>CGCTGGACTGGTTCGAAGTACAGTCAATTCGAGGCCACCTGGTCGACGGAGCGGTCG<br>CGCACTTCCAGGTGACTATGAAAGTCGGCTTCCGCCTGGAGGATTCCTGAcggtttgaaacc<br>ttccgaggagaattcaataattttgtttaactttaagaaggagatatacatATGAGCAATCACACCTACCGAGTGA<br>TCGAGATCGTCGGGACCTCGCCCGACGGCGTCGACGCGGCAATCCAGGGCGGTCTG<br>GCCCGAGCTGCGCAGACCATGCGCGCGCTGGACTGGTTCGAAGTACAGTCAATTCGA<br>GGCCACCTGGTCGACGGAGCGGTCGCGCACTTCCAGGTGACTATGAAAGTCGGCTTC<br>CGCCTGGAGGATTCCTGAacctgtatcgatgcagctaggaagcttaataattttgtttaactttaagaaggagatatac<br>atATGAGCAATCACACCTACCGAGTGATCGAGATCGTCGGGACCTCGCCCGACGGCGT<br>CGACGCGGCAATCCAGGGCGGTCTGGCCCGAGCTGCGCAGACCATGCGCGCGCTGG<br>ACTGGTTCGAAGTACAGTCAATTCGAGGCCACCTGGTCGACGGAGCGGTCGCGCACTT<br>CCAGGTGACTATGAAAGTCGGCTTCCGCCTGGAGGATTCCCTCGAGTCTCCAGCTGCG<br>CCTGCTCCGGCAAGCCCTGCGAGCTGGAGCCACCCGCAGTTCGAAAAATGAgaatcgcga<br>ttcaccatggtgggatccgatccggctgctaacaaagcccgaaaggaagctgagttggctgctgccaccgctgagcaataact<br>agcataaccccttggggcctctaaacgggtcttgaggggttttttg | MSNHTYRVIEIVGTSPDGVDAAIQGGLA<br>RAAQTMRALDWFEVQSIRGHLVDGAVA<br>HFQVTMKVGFRLEDS*<br>&<br>MSNHTYRVIEIVGTSPDGVDAAIQGGLA<br>RAAQTMRALDWFEVQSIRGHLVDGAVA<br>HFQVTMKVGFRLEDS*<br>&<br>MSNHTYRVIEIVGTSPDGVDAAIQGGLA<br>RAAQTMRALDWFEVQSIRGHLVDGAVA<br>HFQVTMKVGFRLEDSLESPAAPAPASPA<br>SWSHPQFEK |
| ***mt*Dod-proteins** | DNA sequence<br>T7 promotor to T7 terminator<br>(CDS uppercase) | Amino acid sequence<br>(Linker) |

| | | |
|---|---|---|
| *mt*Dod-*mm*ACP | cccgcgaaattaatacgactcactataggggaattgtgagcggataacaattcccctctagaaataattttgtttaactttaagaag<br>gagatatacatATGAGCAATCACACCTACCGAGTGATCGAGATCGTCGGGACCTCGCCCGA<br>CGGCGTCGACGCGGCAATCCAGGGCGGTCTGGCCCGAGCTGCGCAGACCATGCGCG<br>CGCTGGACTGGTTCGAAGTACAGTCAATTCGAGGCCACCTGGTCGACGGAGCGGTCG<br>CGCACTTCCAGGTGACTATGAAAGTCGGCTTCCGCCTGGAGGATTCCCTCGAGTCTCC<br>AGCTGCGCCTGCTCCGGCAAGCCCTGCGAGCGACGGGGACACCCAGAGGGATCTGG<br>TGAAAGCTGTAGCACACATCCTAGGCATCCGAGACCTCGCAGGTATTAACCTGGACAG<br>CACGCTGGCAGACCTCGGCCTGGACTCGCTCATGGGTGTGGAAGTTCGTCAGATCCT<br>GGAACGAGAACACGATCTGGTGCTGCCCATGCGTGAGGTGCGGCAGCTCACGCTGCG<br>GAAACTTCAGGAAATGTCCTCCAAGACTGACTCGGCTACTGACACGACAGCCCCCTGA<br>gatccggctgctaacaaagcccgaaaggaagctgagttggctgctgccaccgctgagcaataactagcataaccccttggggc<br>ctctaaacgggtcttgaggggttttttg | MSNHTYRVIEIVGTSPDGVDAAIQGGLA<br>RAAQTMRALDWFEVQSIRGHLVDGAVA<br>HFQVTMKVGFRLEDSLESPAAPAPASPA<br>SDGDTQRDLVKAVAHILGIRDLAGINLDS<br>TLADLGLDSLMGVEVRQILEREHDLVLP<br>MREVRQLTLRKLQEMSSKTDSATDTTAP<br>* |
| *mt*Dod-*mm*ACP-H8 | cccgcgaaattaatacgactcactataggggaattgtgagcggataacaattcccctctagaaataattttgtttaactttaagaag<br>gagatatacatATGAGCAATCACACCTACCGAGTGATCGAGATCGTCGGGACCTCGCCCGA<br>CGGCGTCGACGCGGCAATCCAGGGCGGTCTGGCCCGAGCTGCGCAGACCATGCGCG<br>CGCTGGACTGGTTCGAAGTACAGTCAATTCGAGGCCACCTGGTCGACGGAGCGGTCG<br>CGCACTTCCAGGTGACTATGAAAGTCGGCTTCCGCCTGGAGGATTCCCTCGAGTCTCC<br>AGCTGCGCCTGCTCCGGCAAGCCCTGCGAGCGACGGGGACACCCAGAGGGATCTGG<br>TGAAAGCTGTAGCACACATCCTAGGCATCCGAGACCTCGCAGGTATTAACCTGGACAG<br>CACGCTGGCAGACCTCGGCCTGGACTCGCTCATGGGTGTGGAAGTTCGTCAGATCCT<br>GGAACGAGAACACGATCTGGTGCTGCCCATGCGTGAGGTGCGGCAGCTCACGCTGCG<br>GAAACTTCAGGAAATGTCCTCCAAGACTGACTCGGCTACTGACACGACAGCCCCCCTC<br>GAGCATCATCACCACCACCACCACCACTGAgatccggctgctaacaaagcccgaaaggaagctgagttg<br>gctgctgccaccgctgagcaataactagcataaccccttggggcctctaaacgggtcttgaggggttttttg | MSNHTYRVIEIVGTSPDGVDAAIQGGLA<br>RAAQTMRALDWFEVQSIRGHLVDGAVA<br>HFQVTMKVGFRLEDSLESPAAPAPASPA<br>SDGDTQRDLVKAVAHILGIRDLAGINLDS<br>TLADLGLDSLMGVEVRQILEREHDLVLP<br>MREVRQLTLRKLQEMSSKTDSATDTTAP<br>LEHHHHHHHH* |
| *mt*Dod-msfGFP-H8 | cccgcgaaattaatacgactcactataggggaattgtgagcggataacaattcccctctagaaataattttgtttaactttaagaag<br>gagatatacatATGAGCAATCACACCTACCGAGTGATCGAGATCGTCGGGACCTCGCCCGA<br>CGGCGTCGACGCGGCAATCCAGGGCGGTCTGGCCCGAGCTGCGCAGACCATGCGCG<br>CGCTGGACTGGTTCGAAGTACAGTCAATTCGAGGCCACCTGGTCGACGGAGCGGTCG<br>CGCACTTCCAGGTGACTATGAAAGTCGGCTTCCGCCTGGAGGATTCCCTCGAGTCTCC<br>AGCTGCGCCTGCTCCGGCAAGCCCTGCGAGCGTGAGCAAGGGCGAGGAGCTGTTCAC<br>CGGGGTGGTGCCCATCCTGGTCGAGCTGGACGGCGACGTAAACGGCCACAAGTTCAG<br>CGTGCGCGGCGAGGGCGAGGGCGATGCCACCAACGGCAAGCTGACCCTGAAGTTCAT<br>CTGCACCACCGGCAAGCTGCCCGTGCCCTGGCCCACCCTCGTGACCACCCTGACCTA<br>CGGCGTGCAGTGCTTCAGCCGCTACCCCGACCACATGAAGCAGCACGACTTCTTCAAG<br>TCCGCCATGCCCGAAGGCTACGTCCAGGAGCGCACCATCTCCTTCAAGGACGACGGC<br>ACCTACAAGACCCGCGCCGAGGTGAAGTTCGAGGGCGACACCCTGGTGAACCGCATC<br>GAGCTGAAGGGCATCGACTTCAAGGAGGACGGCAACATCCTGGGGCACAAGCTGGAG<br>TACAACTTCAACAGCCACAACGTCTATATCACGGCCGACAAGCAGAAGAACGGCATCA<br>AGGCGAACTTCAAGATCCGCCACAACGTCGAGGACGGCAGCGTGCAGCTCGCCGACC<br>ACTACCAGCAGAACACCCCCATCGGCGACGGCCCCGTGCTGCTGCCCGACAACCACT<br>ACCTGAGCACCCAGTCCAAGCTGAGCAAAGACCCCAACGAGAAGCGCGATCACATGG<br>TCCTGCTGGAGTTCGTGACCGCCGCCGGGATCACTCTCGGCATGGACGAGCTCGAGC<br>ATCATCACCACCACCACCACCACTGAtgagatccggctgctaacaaagcccgaaaggaagctgagttggctg<br>ctgccaccgctgagcaataactagcataaccccttggggcctctaaacgggtcttgaggggttttttg | MSNHTYRVIEIVGTSPDGVDAAIQGGLA<br>RAAQTMRALDWFEVQSIRGHLVDGAVA<br>HFQVTMKVGFRLEDSLESPAAPAPASPA<br>SVSKGEELFTGVVPILVELDGDVNGHKF<br>SVRGEGEGDATNGKLTLKFICTTGKLPV<br>PWPTLVTTLTYGVQCFSRYPDHMKQHD<br>FFKSAMPEGYVQERTISFKDDGTYKTRA<br>EVKFEGDTLVNRIELKGIDFKEDGNILGH<br>KLEYNFNSHNVYITADKQKNGIKANFKIR<br>HNVEDGSVQLADHYQQNTPIGDGPVLL<br>PDNHYLSTQSKLSKDPNEKRDHMVLLEF<br>VTAAGITLGMDELEHHHHHHHH* |
| *mt*Dod-SpyC-H8* | cccgcgaaattaatacgactcactataggggaattgtgagcggataacaattcccctctagaaataattttgtttaactttaagaag<br>gagatatacatATGAGCAATCACACCTACCGAGTGATCGAGATCGTCGGGACCTCGCCCGA<br>CGGCGTCGACGCGGCAATCCAGGGCGGTCTGGCCCGAGCTGCGCAGACCATGCGCG<br>CGCTGGACTGGTTCGAAGTACAGTCAATTCGAGGCCACCTGGTCGACGGAGCGGTCG<br>CGCACTTCCAGGTGACTATGAAAGTCGGCTTCCGCCTGGAGGATTCCCTCGAGTCTCC<br>AGCTGCGCCTGCTCCGGCAAGCCCTGCGAGCGCTATGGTGGACACCCTGTCGGGCCT<br>CTCTAGTGAACAGGGGCAAAGCGGCGATATGACTATCGAAGAAGATAGTGCTACCCAT<br>ATTAAATTCTCAAAACGTGATGAGGACGGCAAAGAGTTAGCTGGTGCAACTATGGAGTT<br>GCGTGATTCATCTGGTAAAACTATTAGTACATGGATTTCAGATGGACAAGTGAAAGATTT<br>CTACCTGTATCCAGGAAAATATACATTTGTCGAAACCGCAGCACCAGACGGTTATGAGG<br>TAGCAACTGCTATTACCTTTACAGTTAATGAGCAAGGTCAGGTTACTGTAAATGGCAAA<br>GCAACTAAAGGTGACGCTCATATTCTCGAGCATCATCACCACCACCACCACCACTGAgat<br>ccggctgctaacaaagcccgaaaggaagctgagttggctgctgccaccgctgagcaataactagcataaccccttggggcctc<br>taaacgggtcttgaggggttttttg | MSNHTYRVIEIVGTSPDGVDAAIQGGLA<br>RAAQTMRALDWFEVQSIRGHLVDGAVA<br>HFQVTMKVGFRLEDSLESPAAPAPASPA<br>SAMVDTLSGLSSEQGQSGDMTIEEDSA<br>THIKFSKRDEDGKELAGATMELRDSSGK<br>TISTWISDGQVKDFYLYPGKYTFVETAAP<br>DGYEVATAITFTVNEQGQVTVNGKATKG<br>DAHILEHHHHHHHH* |
| H8-SpyC-*mt*Dod | cccgcgaaattaatacgactcactataggggaattgtgagcggataacaattcccctctagaaataattttgtttaactttaagaag<br>gagatatacatATGCATCATCACCACCACCACCATCACGGTTCTGCTATGGTGGACACCCTG<br>TCGGGCCTCTCTAGTGAACAGGGGCAAAGCGGCGATATGACTATCGAAGAAGATAGTG<br>CTACCCATATTAAATTCTCAAAACGTGATGAGGACGGCAAAGAGTTAGCTGGTGCAACT<br>ATGGAGTTGCGTGATTCATCTGGTAAAACTATTAGTACATGGATTTCAGATGGACAAGT<br>GAAAGATTTCTACCTGTATCCAGGAAAATATACATTTGTCGAAACCGCAGCACCAGACG<br>GTTATGAGGTAGCAACTGCTATTACCTTTACAGTTAATGAGCAAGGTCAGGTTACTGTA<br>AATGGCAAAGCAACTAAAGGTGACGCTCATATTTCTCCAGCTGCGCCTGCTCCGGCAA<br>GCCCTGCGAGCAGCAATCACACCTACCGAGTGATCGAGATCGTCGGGACCTCGCCCG<br>ACGGCGTCGACGCGGCAATCCAGGGCGGTCTGGCCCGAGCTGCGCAGACCATGCGC<br>GCGCTGGACTGGTTCGAAGTACAGTCAATTCGAGGCCACCTGGTCGACGGAGCGGTC<br>GCGCACTTCCAGGTGACTATGAAAGTCGGCTTCCGCCTGGAGGATTCCTGActcgagcacc<br>accaccaccaccactgagatccggctgctaacaaagcccgaaaggaagctgagttggctgctgccaccgctgagcaataact<br>agcataaccccttggggcctctaaacgggtcttgaggggttttttg | MHHHHHHHHGSAMVDTLSGLSSEQGQ<br>SGDMTIEEDSATHIKFSKRDEDGKELAG<br>ATMELRDSSGKTISTWISDGQVKDFYLY<br>PGKYTFVETAAPDGYEVATAITFTVNEQ<br>GQVTVNGKATKGDAHISPAAPAPASPAS<br>SNHTYRVIEIVGTSPDGVDAAIQGGLARA<br>AQTMRALDWFEVQSIRGHLVDGAVAHF<br>QVTMKVGFRLEDS* |
| *mt*Dod-SZ1 | cccgcgaaattaatacgactcactataggggaattgtgagcggataacaattcccctctagaaataattttgtttaactttaagaag<br>gagatatacatATGAGCAATCACACCTACCGAGTGATCGAGATCGTCGGGACCTCGCCCGA<br>CGGCGTCGACGCGGCAATCCAGGGCGGTCTGGCCCGAGCTGCGCAGACCATGCGCG<br>CGCTGGACTGGTTCGAAGTACAGTCAATTCGAGGCCACCTGGTCGACGGAGCGGTCG<br>CGCACTTCCAGGTGACTATGAAAGTCGGCTTCCGCCTGGAGGATTCCCTCGAGTCTCC<br>AGCTGCGCCTGCTCCGGCAAGCCCTGCGAGCAATCTAGTCGCTCAGCTAGAGAACGA<br>AGTAGCATCATTAGAGAATGAAAACGAAACCTTGAAAAAGAAGAATCTACACAAAAAGG<br>ATCTTATAGCCTACCTAGAAAAGGAAATTGCTAACTTAAGGAAAAAGATTGAGGAATGAg<br>gatcccaccaccaccaccaccactgagatccggctgctaacaaagcccgaaaggaagctgagttggctgctgccaccgctga<br>gcaataactagcataaccccttggggcctctaaacgggtcttgaggggttttttg | MSNHTYRVIEIVGTSPDGVDAAIQGGLA<br>RAAQTMRALDWFEVQSIRGHLVDGAVA<br>HFQVTMKVGFRLEDSLESPAAPAPASPA<br>SNLVAQLENEVASLENENETLKKKNLHK<br>KDLIAYLEKEIANLRKKIEE* |
| *mt*Dod-SZ3 | cccgcgaaattaatacgactcactataggggaattgtgagcggataacaattcccctctagaaataattttgtttaactttaagaag<br>gagatatacatATGAGCAATCACACCTACCGAGTGATCGAGATCGTCGGGACCTCGCCCGA<br>CGGCGTCGACGCGGCAATCCAGGGCGGTCTGGCCCGAGCTGCGCAGACCATGCGCG<br>CGCTGGACTGGTTCGAAGTACAGTCAATTCGAGGCCACCTGGTCGACGGAGCGGTCG<br>CGCACTTCCAGGTGACTATGAAAGTCGGCTTCCGCCTGGAGGATTCCCTCGAGTCTCC<br>AGCTGCGCCTGCTCCGGCAAGCCCTGCGAGCAACGAAGTTACAACACTTGAGAATGAC<br>GCTGCCTTTATCGAAAATGAAAATGCTTATCTAGAAAAAGAGATAGCACGTTTGAGAAA<br>GGAGAAAGCAGCATTGAGAAATAGACTGGCACACAAAAAGTGAggatcccaccaccaccaccac<br>cactgagatccggctgctaacaaagcccgaaaggaagctgagttggctgctgccaccgctgagcaataactagcataacccct<br>tggggcctctaaacgggtcttgaggggttttttg | MSNHTYRVIEIVGTSPDGVDAAIQGGLA<br>RAAQTMRALDWFEVQSIRGHLVDGAVA<br>HFQVTMKVGFRLEDSLESPAAPAPASPA<br>SNEVTTLENDAAFIENENAYLEKEIARLR<br>KEKAALRNRLAHKK* |

| | | |
|---|---|---|
| SZ1-*mt*Dod | cccgcgaaattaatacgactcactataggggaattgtgagcggataacaattcccctctagaaataattttgtttaactttaagaaggagatatacatATGAATCTAGTCGCTCAGCTAGAGAACGAAGTAGCATCATTAGAGAATGAAAACGAAACCTTGAAAAAGAAGAATCTACACAAAAAGGATCTTATAGCCTACCTAGAAAAGGAAATTGCTAACTTAAGGAAAAAGATTGAGGAATCTCCAGCTGCGCCTGCTCCGGCAAGCCCTGCGAGCAGCAATCACACCTACCGAGTGATCGAGATCGTCGGGACCTCGCCCGACGGCGTCGACGCGGCAATCCAGGGCGGTCTGGCCCGAGCTGCGCAGACCATGCGCGCGCTGGACTGGTTCGAAGTACAGTCAATTCGAGGCCACCTGGTCGACGGAGCGGTCGCGCACTTCCAGGTGACTATGAAAGTCGGCTTCCGCCTGGAGGATTCCTGAggatcccaccaccaccaccaccactgagatccggctgctaacaaagcccgaaaggaagctgagttggctgctgccaccgctgagcaataactagcataaccccttggggcctctaaacgggtcttgaggggttttttg | MNLVAQLENEVASLENENETLKKKNLHKKDLIAYLEKEIANLRKKIEESPAAPAPASPASSNHTYRVIEIVGTSPDGVDAAIQGGLARAAQTMRALDWFEVQSIRGHLVDGAVAHFQVTMKVGFRLEDS* |
| SZ3-*mt*Dod | cccgcgaaattaatacgactcactataggggaattgtgagcggataacaattcccctctagaaataattttgtttaactttaagaaggagatatacatATGAACGAAGTTACAACACTTGAGAATGACGCTGCCTTTATCGAAAATGAAAATGCTTATCTAGAAAAAGAGATAGCACGTTTGAGAAAGGAGAAAGCAGCATTGAGAAATAGACTGGCACACAAAAAGTCTCCAGCTGCGCCTGCTCCGGCAAGCCCTGCGAGCAGCAATCACACCTACCGAGTGATCGAGATCGTCGGGACCTCGCCCGACGGCGTCGACGCGGCAATCCAGGGCGGTCTGGCCCGAGCTGCGCAGACCATGCGCGCGCTGGACTGGTTCGAAGTACAGTCAATTCGAGGCCACCTGGTCGACGGAGCGGTCGCGCACTTCCAGGTGACTATGAAAGTCGGCTTCCGCCTGGAGGATTCCTGAggatcccaccaccaccaccaccactgagatccggctgctaacaaagcccgaaaggaagctgagttggctgctgccaccgctgagcaataactagcataaccccttggggcctctaaacgggtcttgaggggttttttg | MNEVTTLENDAAFIENENAYLEKEIARLRKEKAALRNRLAHKKSPAAPAPASPASSNHTYRVIEIVGTSPDGVDAAIQGGLARAAQTMRALDWFEVQSIRGHLVDGAVAHFQVTMKVGFRLEDS* |
| *mt*Dod-*se*ACP** | cccgcgaaattaatacgactcactataggggaattgtgagcggataacaattcccctctagaaataattttgtttaactttaagaaggagatatacatATGAGCAATCACACCTACCGAGTGATCGAGATCGTCGGGACCTCGCCCGACGGCGTCGACGCGGCAATCCAGGGCGGTCTGGCCCGAGCTGCGCAGACCATGCGCGCGCTGGACTGGTTCGAAGTACAGTCAATTCGAGGCCACCTGGTCGACGGAGCGGTCGCGCACTTCCAGGTGACTATGAAAGTCGGCTTCCGCCTGGAGGATTCCCTcgagTCTCCAGCTGCGCCTGCTCCGGCAAGCCCTGCGAGCCGCGTAGACGAGATCGAGGACAAGTTGGGAAACTATATCCGCAGGCACCTGCTGACTGAGGACCCTCCAGAGGAATTCACTTACTCCACCGCCCTCTTCGGCGATGGGGTGCTGGATTCGCTCCGGCTGGCGATGCTGATCAACTTCATCCGCAACGAGCTGGCCGTGGAGATCCCGTACGAGCACGTGAACCGGGACGACTTCCACGATGTGCACACTATCGCCAAGATGGTGGTCGGCCTGTCGAGCGAAGCGAAACTCGAGCATCATCACCACCACCACCACCACTGAgatccggctgctaacaaagcccgaaaggaagctgagttggctgctgccaccgctgagcaataactagcataaccccttggggcctctaaacgggtcttgaggggttttttg | MSNHTYRVIEIVGTSPDGVDAAIQGGLARAAQTMRALDWFEVQSIRGHLVDGAVAHFQVTMKVGFRLEDSLESPAAPAPASPASRVDEIEDKLGNYIRRHLLTEDPPEEFTYSTALFGDGVLDSLRLAMLINFIRNELAVEIPYEHVNRDDFHDVHTIAKMVVGLSSEAKLEHHHHHHHH* |
| **non-dodecin *constructs*** | DNA sequence<br>T7 promotor to T7 terminator<br>(CDS uppercase) | Amino acid sequence<br>(Linker) |
| SpyT-*se*ACP | cccgcgaaattaatacgactcactataggggaattgtgagcggataacaattcccctctagaaataattttgtttaactttaagaaggagatatacatATGGCACATATCGTCATGGTTGATGCGTACAAACCGACCAAAGGTGGCAGCGGTTCTCCAGCTGCGCCTGCTCCGGCAAGCCCTGCGAGCCGCGTAGACGAGATCGAGGACAAGTTGGGAAACTATATCCGCAGGCACCTGCTGACTGAGGACCCTCCAGAGGAATTCACTTACTCCACCGCCCTCTTCGGCGATGGGGTGCTGGATTCGCTCCGGCTGGCGATGCTGATCAACTTCATCCGCAACGAGCTGGCCGTGGAGATCCCGTACGAGCACGTGAACCGGGACGACTTCCACGATGTGCACACTATCGCCAAGATGGTGGTCGGCCTGTCGAGCGAAGCGAAACTCGAGCATCATCACCACCACCACCACCACTGAgatccggctgctaacaaagcccgaaaggaagctgagttggctgctgccaccgctgagcaataactagcataaccccttggggcctctaaacgggtcttgaggggttttttg | MAHIVMVDAYKPTKGGSGSPAAPAPASPASRVDEIEDKLGNYIRRHLLTEDPPEEFTYSTALFGDGVLDSLRLAMLINFIRNELAVEIPYEHVNRDDFHDVHTIAKMVVGLSSEAKLEHHHHHHHH* |
| *se*ACP-SpyC | cccgcgaaattaatacgactcactataggggaattgtgagcggataacaattcccctctagaaataattttgtttaactttaagaaggagatatacatATGCGCGTAGACGAGATCGAGGACAAGTTGGGAAACTATATCCGCAGGCACCTGCTGACTGAGGACCCTCCAGAGGAATTCACTTACTCCACCGCCCTCTTCGGCGATGGGGTGCTGGATTCGCTCCGGCTGGCGATGCTGATCAACTTCATCCGCAACGAGCTGGCCGTGGAGATCCCGTACGAGCACGTGAACCGGGACGACTTCCACGATGTGCACACTATCGCCAAGATGGTGGTCGGCCTGTCGAGCGAAGCGAAAGGTGGCAGCGGTGCTATGGTGGACACCCTGTCGGGCCTCTCTAGTGAACAGGGGCAAAGCGGCGATATGACTATCGAAGAAGATAGTGCTACCCATATTAAATTCTCAAAACGTGATGAGGACGGCAAAGAGTTAGCTGGTGCAACTATGGAGTTGCGTGATTCATCTGGTAAAACTATTAGTACATGGATTTCAGATGGACAAGTGAAAGATTTCTACCTGTATCCAGGAAAATATACATTTGTCGAAACCGCAGCACCAGACGGTTATGAGGTAGCAACTGCTATTACCTTTACAGTTAATGAGCAAGGTCAGGTTACTGTAAATGGCAAAGCAACTAAAGGTGACGCTCATATTCTCGAGCATCATCACCACCACCACCACCACTGAgatccggctgctaacaaagcccgaaaggaagctgagttggctgctgccaccgctgagcaataactagcataaccccttggggcctctaaacgggtcttgaggggtttttttgctgaaag | MRVDEIEDKLGNYIRRHLLTEDPPEEFTYSTALFGDGVLDSLRLAMLINFIRNELAVEIPYEHVNRDDFHDVHTIAKMVVGLSSEAKGGSGAMVDTLSGLSSEQGQSGDMTIEEDSATHIKFSKRDEDGKELAGATMELRDSSGKTISTWISDGQVKDFYLYPGKYTFVETAAPDGYEVATAITFTVNEQGQVTVNGKATKGDAHILEHHHHHHHH* |
| mClover3-SnpC | cccgcgaaattaatacgactcactataggggaattgtgagcggataacaattcccctctagaaataattttgtttaactttaagaaggagatatacatATGGTGAGCAAGGGCGAGGAGCTGTTCACCGGGGTGGTGCCCATCCTGGTCGAGCTGGACGGCGACGTAAACGGCCACAAGTTCAGCGTCCGCGGCGAGGGCGAGGGCGATGCCACCAACGGCAAGCTGACCCTGAAGTTCATCTGCACCACCGGCAAGCTGCCCGTGCCCTGGCCCACCCTCGTGACCACCTTCGGCTACGGCGTGGCCTGCTTCAGCCGCTACCCCGACCACATGAAGCAGCACGACTTCTTCAAGTCCGCCATGCCCGAAGGCTACGTCCAGGAGCGCACCATCTCTTTCAAGGACGACGGTACCTACAAGACCCGCGCCGAGGTGAAGTTCGAGGGCGACACCCTGGTGAACCGCATCGAGCTGAAGGGCATCGACTTCAAGGAGGACGGCAACATCCTGGGGCACAAGCTGGAGTACAACTTCAACAGCCACTACGTCTATATCACGGCCGACAAGCAGAAGAACTGCATCAAGGCTAACTTCAAGATCCGCCACAACGTTGAGGACGGCAGCGTGCAGCTCGCCGACCACTACCAGCAGAACACCCCCATCGGCGACGGCCCCGTGCTGCTGCCCGACAACCACTACCTGAGCCATCAGTCCAAGCTGAGCAAAGACCCCAACGAGAAGCGCGATCACATGGTCCTGCTGGAGTTCGTGACCGCCGCCGGCATTACCCATGGCATGGATGAACTGTATAAAGGTGGCAGCGGTAGCGGTAGCGGCAAGCCGCTGCGTGGTGCCGTGTTTAGCCTGCAGAAACAGCATCCCGACTATCCCGATATCTATGGCGCGATTGATCAGAATGGGACCTATCAAAATGTGCGTACCGGCGAAGATGGTAAACTGACCTTTAAGAATCTGAGCGATGGCAAATATCGCCTGTTTGAAAATAGCGAACCCGCTGGCTATAAACCGGTGCAGAATAAGCCGATTGTGGCGTTTCAGATTGTGAATGGCGAAGTGCGTGATGTGACCAGCATTGTGCCGCAGGATATTCCGGCTACATATGAATTTACCAACGGTAAACATTATATCACCAATGAACCGATACCGCCGAAACTCGAGCATCATCACCACCACCACCACCACTGAgatccggctgctaacaaagcccgaaaggaagctgagttggctgctgccaccgctgagcaataactagcataaccccttggggcctctaaacgggtcttgaggggtttttttgctgaaag | MVSKGEELFTGVVPILVELDGDVNGHKFSVRGEGEGDATNGKLTLKFICTTGKLPVPWPTLVTTFGYGVACFSRYPDHMKQHDFFKSAMPEGYVQERTISFKDDGTYKTRAEVKFEGDTLVNRIELKGIDFKEDGNILGHKLEYNFNSHYVYITADKQKNCIKANFKIRHNVEDGSVQLADHYQQNTPIGDGPVLLPDNHYLSHQSKLSKDPNEKRDHMVLLEFVTAAGITHGMDELYKGGSGSGSGKPLRGAVFSLQKQHPDYPDIYGAIDQNGTYQNVRTGEDGKLTFKNLSDGKYRLFENSEPAGYKPVQNKPIVAFQIVNGEVRDVTSIVPQDIPATYEFTNGKHYITNEPIPPKLEHHHHHHHH* |

| | | |
|---|---|---|
| SpyT-mClover3 | cccgcgaaattaatacgactcactataggggaattgtgagcggataacaattcccctctagaaataattttgtttaactttaagaag<br>gagatatacatATGGCACATATCGTCATGGTTGATGCGTACAAACCGACCAAAGGTGGCAGC<br>GGTTCTGTGAGCAAGGGCGAGGAGCTGTTCACCGGGGTGGTGCCCATCCTGGTCGAG<br>CTGGACGGCGACGTAAACGGCCACAAGTTCAGCGTCCGCGGCGAGGGCGAGGGCGA<br>TGCCACCAACGGCAAGCTGACCCTGAAGTTCATCTGCACCACCGGCAAGCTGCCCGT<br>GCCCTGGCCCACCCTCGTGACCACCTTCGGCTACGGCGTGGCCTGCTTCAGCCGCTA<br>CCCCGACCACATGAAGCAGCACGACTTCTTCAAGTCCGCCATGCCCGAAGGCTACGTC<br>CAGGAGCGCACCATCTCTTTCAAGGACGACGGTACCTACAAGACCCGCGCCGAGGTG<br>AAGTTCGAGGGCGACACCCTGGTGAACCGCATCGAGCTGAAGGGCATCGACTTCAAG<br>GAGGACGGCAACATCCTGGGGCACAAGCTGGAGTACAACTTCAACAGCCACTACGTCT<br>ATATCACGGCCGACAAGCAGAAGAACTGCATCAAGGCTAACTTCAAGATCCGCCACAA<br>CGTTGAGGACGGCAGCGTGCAGCTCGCCGACCACTACCAGCAGAACACCCCCATCGG<br>CGACGGCCCCGTGCTGCTGCCCGACAACCACTACCTGAGCCATCAGTCCAAGCTGAG<br>CAAAGACCCCAACGAGAAGCGCGATCACATGGTCCTGCTGGAGTTCGTGACCGCCGC<br>CGGCATTACCCATGGCATGGATGAACTGTATAAAcTCGAGCATCATCACCACCACCACC<br>ACCACTGAgatccggctgctaacaaagcccgaaaggaagctgagttggctgctgccaccgctgagcaataactagcata<br>acccccttggggcctctaaacgggtcttgaggggttttttg | MAHIVMVDAYKPTKGGSGSVSKGEELFT<br>GVVPILVELDGDVNGHKFSVRGEGEGD<br>ATNGKLTLKFICTTGKLPVPWPTLVTTFG<br>YGVACFSRYPDHMKQHDFFKSAMPEGY<br>VQERTISFKDDGTYKTRAEVKFEGDTLV<br>NRIELKGIDFKEDGNILGHKLEYNFNSHY<br>VYITADKQKNCIKANFKIRHNVEDGSVQL<br>ADHYQQNTPIGDGPVLLPDNHYLSHQSK<br>LSKDPNEKRDHMVLLEFVTAAGITHGMD<br>ELYKLEHHHHHHHH* |
| SZ2-mRuby3 | cccgcgaaattaatacgactcactataggggaattgtgagcggataacaattcccctctagaaataattttgtttaactttaagaag<br>gagatatacatATGGCCAGAAATGCATACTTAAGGAAAAAGATTGCTAGATTGAAAAAGGAC<br>AACTTACAATTAGAAAGAGATGAGCAAAATCTTGAAAAGATCATTGCCAATTTGAGAGAT<br>GAAATCGCCAGACTTGAAAATGAGGTGGCCTCTCATGAACAAGGTGGCAGCGGTGTGT<br>CTAAGGGCGAAGAGCTGATCAAGGAAAATATGCGTATGAAGGTGGTCATGGAAGGTTC<br>GGTCAACGGCCACCAATTCAAATGCACAGGTGAAGGAGAAGGCAGACCGTACGAGGG<br>AGTGCAAACCATGAGGATCAAAGTCATCGAGGGAGGACCCCTGCCATTTGCCTTTGAC<br>ATTCTTGCCACGTCGTTCATGTATGGCAGCCGTACCTTTATCAAGTACCCGGCCGACAT<br>CCCTGATTTCTTTAAACAGTCCTTTCCTGAGGGTTTTACTTGGGAAAGAGTTACGAGATA<br>CGAAGATGGTGGAGTCGTCACCGTCACGCAGGACACCAGCCTTGAGGATGGCGAGCT<br>CGTCTACAACGTCAAGGTCAGAGGGGTAAACTTTCCCTCCAATGGTCCCGTGATGCAG<br>AAGAAGACCAAGGGTTGGGAGCCTAATACAGAGATGATGTATCCAGCAGATGGTGGTC<br>TGAGAGGATACACTGACATCGCACTGAAAGTTGATGGTGGTGGCCATCTGCACTGCAA<br>CTTCGTGACAACTTACAGGTCAAAAAAGACCGTCGGGAACATCAAGATGCCCGGTGTC<br>CATGCCGTTGATCACCGCCTGGAAAGGATCGAGGAGAGTGACAATGAAACCTACGTAG<br>TGCAAAGAGAAGTGGCAGTTGCCAAATACAGCAACCTTGGTGGTGGCATGGACGAGCT<br>GTACAAGCTCGAGCATCATCACCACCACCACCACCACTGAgatccggctgctaacaaagcccgaa<br>aggaagctgagttggctgctgccaccgctgagcaataactagcataacccccttggggcctctaaacgggtcttgaggggttttttg | MARNAYLRKKIARLKKDNLQLERDEQNL<br>EKIIANLRDEIARLENEVASHEQGGSGVS<br>KGEELIKENMRMKVVMEGSVNGHQFKC<br>TGEGEGRPYEGVQTMRIKVIEGGPLPFA<br>FDILATSFMYGSRTFIKYPADIPDFFKQSF<br>PEGFTWERVTRYEDGGVVTVTQDTSLE<br>DGELVYNVKVRGVNFPSNGPVMQKKTK<br>GWEPNTEMMYPADGGLRGYTDIALKVD<br>GGGHLHCNFVTTYRSKKTVGNIKMPGV<br>HAVDHRLERIEESDNETYVVQREVAVAK<br>YSNLGGGMDELYKLEHHHHHHHH* |
| SZ2-mClover3 | cccgcgaaattaatacgactcactataggggaattgtgagcggataacaattcccctctagaaataattttgtttaactttaagaag<br>gagatatacatATGGCCAGAAATGCATACTTAAGGAAAAAGATTGCTAGATTGAAAAAGGAC<br>AACTTACAATTAGAAAGAGATGAGCAAAATCTTGAAAAGATCATTGCCAATTTGAGAGAT<br>GAAATCGCCAGACTTGAAAATGAGGTGGCCTCTCATGAACAAGGTGGCAGCGGTGTGA<br>GCAAGGGCGAGGAGCTGTTCACCGGGGTGGTGCCCATCCTGGTCGAGCTGGACGGC<br>GACGTAAACGGCCACAAGTTCAGCGTCCGCGGCGAGGGCGAGGGCGATGCCACCAA<br>CGGCAAGCTGACCCTGAAGTTCATCTGCACCACCGGCAAGCTGCCCGTGCCCTGGCC<br>CACCCTCGTGACCACCTTCGGCTACGGCGTGGCCTGCTTCAGCCGCTACCCCGACCA<br>CATGAAGCAGCACGACTTCTTCAAGTCCGCCATGCCCGAAGGCTACGTCCAGGAGCG<br>CACCATCTCTTTCAAGGACGACGGTACCTACAAGACCCGCGCCGAGGTGAAGTTCGAG<br>GGCGACACCCTGGTGAACCGCATCGAGCTGAAGGGCATCGACTTCAAGGAGGACGGC<br>AACATCCTGGGGCACAAGCTGGAGTACAACTTCAACAGCCACTACGTCTATATCACGG<br>CCGACAAGCAGAAGAACTGCATCAAGGCTAACTTCAAGATCCGCCACAACGTTGAGGA<br>CGGCAGCGTGCAGCTCGCCGACCACTACCAGCAGAACACCCCCATCGGCGACGGCCC<br>CGTGCTGCTGCCCGACAACCACTACCTGAGCCATCAGTCCAAGCTGAGCAAAGACCCC<br>AACGAGAAGCGCGATCACATGGTCCTGCTGGAGTTCGTGACCGCCGCCGGCATTACC<br>CATGGCATGGATGAACTGTATAAACTCGAGCATCATCACCACCACCACCACCACTGAgat<br>ccggctgctaacaaagcccgaaaggaagctgagttggctgctgccaccgctgagcaataactagcataacccccttggggcctc<br>taaacgggtcttgaggggttttttg | MARNAYLRKKIARLKKDNLQLERDEQNL<br>EKIIANLRDEIARLENEVASHEQGGSGVS<br>KGEELFTGVVPILVELDGDVNGHKFSVR<br>GEGEGDATNGKLTLKFICTTGKLPVPWP<br>TLVTTFGYGVACFSRYPDHMKQHDFFK<br>SAMPEGYVQERTISFKDDGTYKTRAEVK<br>FEGDTLVNRIELKGIDFKEDGNILGHKLE<br>YNFNSHYVYITADKQKNCIKANFKIRHNV<br>EDGSVQLADHYQQNTPIGDGPVLLPDN<br>HYLSHQSKLSKDPNEKRDHMVLLEFVTA<br>AGITHGMDELYKLEHHHHHHHH* |
| SZ4-mRuby3 | cccgcgaaattaatacgactcactataggggaattgtgagcggataacaattcccctctagaaataattttgtttaactttaagaag<br>gagatatacatATGCAGAAAGTGGCTGAATTGAAAAACAGAGTTGCTGTTAAACTTAACAGAA<br>ATGAACAATTGAAAAACAAGGTAGAAGAGTTGAAAAACCGTAATGCTTACCTGAAAAAC<br>GAACTGGCTACATTAGAAAATGAAGTCGCCAGATTGGAGAACGATGTTGCTGAAGGTG<br>GCAGCGGTGTGTCTAAGGGCGAAGAGCTGATCAAGGAAAATATGCGTATGAAGGTGGT<br>CATGGAAGGTTCGGTCAACGGCCACCAATTCAAATGCACAGGTGAAGGAGAAGGCAGA<br>CCGTACGAGGGAGTGCAAACCATGAGGATCAAAGTCATCGAGGGAGGACCCCTGCCA<br>TTTGCCTTTGACATTCTTGCCACGTCGTTCATGTATGGCAGCCGTACCTTTATCAAGTAC<br>CCGGCCGACATCCCTGATTTCTTTAAACAGTCCTTTCCTGAGGGTTTTACTTGGGAAAG<br>AGTTACGAGATACGAAGATGGTGGAGTCGTCACCGTCACGCAGGACACCAGCCTTGAG<br>GATGGCGAGCTCGTCTACAACGTCAAGGTCAGAGGGGTAAACTTTCCCTCCAATGGTC<br>CCGTGATGCAGAAGAAGACCAAGGGTTGGGAGCCTAATACAGAGATGATGTATCCAGC<br>AGATGGTGGTCTGAGAGGATACACTGACATCGCACTGAAAGTTGATGGTGGTGGCCAT<br>CTGCACTGCAACTTCGTGACAACTTACAGGTCAAAAAAGACCGTCGGGAACATCAAGAT<br>GCCCGGTGTCCATGCCGTTGATCACCGCCTGGAAAGGATCGAGGAGAGTGACAATGA<br>AACCTACGTAGTGCAAAGAGAAGTGGCAGTTGCCAAATACAGCAACCTTGGTGGTGGC<br>ATGGACGAGCTGTACAAGCTCGAGCATCATCACCACCACCACCACCACTGAgatccggctg<br>ctaacaaagcccgaaaggaagctgagttggctgctgccaccgctgagcaataactagcataacccccttggggcctctaaacgg<br>gtcttgaggggttttttg | MQKVAELKNRVAVKLNRNEQLKNKVEE<br>LKNRNAYLKNELATLENEVARLENDVAE<br>GGSGVSKGEELIKENMRMKVVMEGSVN<br>GHQFKCTGEGEGRPYEGVQTMRIKVIE<br>GGPLPFAFDILATSFMYGSRTFIKYPADI<br>PDFFKQSFPEGFTWERVTRYEDGGVVT<br>VTQDTSLEDGELVYNVKVRGVNFPSNG<br>PVMQKKTKGWEPNTEMMYPADGGLRG<br>YTDIALKVDGGGHLHCNFVTTYRSKKTV<br>GNIKMPGVHAVDHRLERIEESDNETYVV<br>QREVAVAKYSNLGGGMDELYKLEHHHH<br>HHHH* |
| SZ4-mClover3 | cccgcgaaattaatacgactcactataggggaattgtgagcggataacaattcccctctagaaataattttgtttaactttaagaag<br>gagatatacatATGCAGAAAGTGGCTGAATTGAAAAACAGAGTTGCTGTTAAACTTAACAGAA<br>ATGAACAATTGAAAAACAAGGTAGAAGAGTTGAAAAACCGTAATGCTTACCTGAAAAAC<br>GAACTGGCTACATTAGAAAATGAAGTCGCCAGATTGGAGAACGATGTTGCTGAAGGTG<br>GCAGCGGTGTGAGCAAGGGCGAGGAGCTGTTCACCGGGGTGGTGCCCATCCTGGTC<br>GAGCTGGACGGCGACGTAAACGGCCACAAGTTCAGCGTCCGCGGCGAGGGCGAGGG<br>CGATGCCACCAACGGCAAGCTGACCCTGAAGTTCATCTGCACCACCGGCAAGCTGCC<br>CGTGCCCTGGCCCACCCTCGTGACCACCTTCGGCTACGGCGTGGCCTGCTTCAGCCG<br>CTACCCCGACCACATGAAGCAGCACGACTTCTTCAAGTCCGCCATGCCCGAAGGCTAC<br>GTCCAGGAGCGCACCATCTCTTTCAAGGACGACGGTACCTACAAGACCCGCGCCGAG<br>GTGAAGTTCGAGGGCGACACCCTGGTGAACCGCATCGAGCTGAAGGGCATCGACTTC<br>AAGGAGGACGGCAACATCCTGGGGCACAAGCTGGAGTACAACTTCAACAGCCACTAC<br>GTCTATATCACGGCCGACAAGCAGAAGAACTGCATCAAGGCTAACTTCAAGATCCGCC<br>ACAACGTTGAGGACGGCAGCGTGCAGCTCGCCGACCACTACCAGCAGAACACCCCCA<br>TCGGCGACGGCCCCGTGCTGCTGCCCGACAACCACTACCTGAGCCATCAGTCCAAGC<br>TGAGCAAAGACCCCAACGAGAAGCGCGATCACATGGTCCTGCTGGAGTTCGTGACCG<br>CCGCCGGCATTACCCATGGCATGGATGAACTGTATAAACTCGAGCATCATCACCACCA<br>CCACCACCACTGAgatccggctgctaacaaagcccgaaaggaagctgagttggctgctgccaccgctgagcaataac<br>tagcataacccccttggggcctctaaacgggtcttgaggggttttttg | MQKVAELKNRVAVKLNRNEQLKNKVEE<br>LKNRNAYLKNELATLENEVARLENDVAE<br>GGSGVSKGEELFTGVVPILVELDGDVNG<br>HKFSVRGEGEGDATNGKLTLKFICTTGK<br>LPVPWPTLVTTFGYGVACFSRYPDHMK<br>QHDFFKSAMPEGYVQERTISFKDDGTYK<br>TRAEVKFEGDTLVNRIELKGIDFKEDGNI<br>LGHKLEYNFNSHYVYITADKQKNCIKANF<br>KIRHNVEDGSVQLADHYQQNTPIGDGPV<br>LLPDNHYLSHQSKLSKDPNEKRDHMVLL<br>EFVTAAGITHGMDELYKLEHHHHHHHH* |

| | | |
|---|---|---|
| *mm*ACP | cccgcgaaattaatacgactcactataggggaattgtgagcggataacaattcccctctagaaataattttgtttaactttaagaag<br>gagatatacatATGAGCGCTTGGAGCCACCCGCAGTTCGAAAAAGGCGCCGGGGACGGGG<br>ACACCCAGAGGGATCTGGTGAAAGCTGTAGCACACATCCTAGGCATCCGAGACCTCGC<br>AGGTATTAACCTGGACAGCACGCTGGCAGACCTCGGCCTGGACTCGCTCATGGGTGT<br>GGAAGTTCGTCAGATCCTGGAACGAGAACACGATCTGGTGCTGCCCATGCGTGAGGT<br>GCGGCAGCTCACGCTGCGGAAACTTCAGGAAATGTCCTCCAAGACTGACTCGGCTACT<br>GACACGACAGCCCCCCTCGAGCATCATCACCACCACCACCACCACTGAgatccggctgctaa<br>caaagcccgaaaggaagctgagttggctgctgccaccgctgagcaataactagcataaccccttggggcctctaaacgggtctt<br>gaggggttttttg | MSAWSHPQFEKGAGDGDTQRDLVKAV<br>AHILGIRDLAGINLDSTLADLGLDSLMGV<br>EVRQILEREHDLVLPMREVRQLTLRKLQ<br>EMSSKTDSATDTTAPLEHHHHHHHH* |
| Sfp | tcataaaaaatttatttgctttgtgagcggataacaattataatagattcaattgtgagcggataacaatttcacacagaattctgcag<br>acggaggatctagaATGAAGATTTACGGAATTTATATGGACCGCCCGCTTTCACAGGAAGAAA<br>ATGAACGGTTCATGACTTTCATATCACCTGAAAAACGGGAGAAATGCCGGAGATTTTAT<br>CATAAAGAAGATGCTCACCGCACCCTGCTGGGAGATGTGCTCGTTCGCTCAGTCATAA<br>GCAGGCAGTATCAGTTGGACAAATCCGATATCCGCTTTAGCACGCAGGAATACGGGAA<br>GCCGTGCATCCCTGATCTTCCCGACGCTCATTTCAACATTTCTCACTCCGGCCGCTGG<br>GTCATTGGTGCGTTTGATTCACAGCCGATCGGCATAGATATCGAAAAAACGAAACCGAT<br>CAGCCTTGAGATCGCCAAGCGCTTCTTTTCAAAAACAGAGTACAGCGACCTTTTAGCAA<br>AAGACAAGGACGAGCAGACAGACTATTTTTATCATCTATGGTCAATGAAAGAAAGCTTT<br>ATCAAACAGGAAGGCAAAGGCTTATCGCTCCCGCTTGATTCCTTTTCAGTGCGCCTGCA<br>TCAGGACGGACAAGTATCCATTGAGCTTCCGGACAGCCATTCCCCATGCTATATCAAAA<br>CGTATGAGGTCGATCCCGGCTACAAAATGGCTGTATGCGCCGCACACCCTGATTTCCC<br>CGAGGATATCACAATGGTCTCGTACGAAGAGCTTTTAAGATCTCATCACCATCACCATC<br>ACTAAgcttaattagctgagcttggactcctgttgatagatccagtaatgacctcagaactccatctggatttgttcagaacgctc | MKIYGIYMDRPLSQEENERFMTFISPEKR<br>EKCRRFYHKEDAHRTLLGDVLVRSVISR<br>QYQLDKSDIRFSTQEYGKPCIPDLPDAH<br>FNISHSGRWVIGAFDSQPIGIDIEKTKPIS<br>LEIAKRFFSKTEYSDLLAKDKDEQTDYFY<br>HLWSMKESFIKQEGKGLSLPLDSFSVRL<br>HQDGQVSIELPDSHSPCYIKTYEVDPGY<br>KMAVCAAHPDFPEDITMVSYEELLRSHH<br>HHHH* |

**Table S2:** Amino acid sequence of proteins from which the peptide sequences were selected for the *mt*Dod-PAS-Pep constructs. Selected peptide sequences highlighted in red.

| Construct<br>&<br>peptide | Identifier<br>&<br>name | Amino acid sequence<br>(peptide) |
|---|---|---|
| *mt*Dod-PAS-Pep1<br>PKGGSGSGPTIEEVD | >AAD21816.1 HSP70-1<br>[Homo sapiens] | MAKAAAIGIDLGTTYSCVGVFQHGKVEIIANDQGNRTTPSYVAFTDTERLIGDAAKNQVALNPQNTVFDAKRLIGRKF<br>GDPVVQSDMKHWPFQVINDGDKPKVQVSYKGDTKAFYPEEISSMVLTKMKEIAEAYLGYPVTNAVITVPAYFNDSQ<br>RQATKDAGVIAGLNVLRIINEPTAAAIAYGLDRTGKGERNVLIFDLGGGTFDVSILTIDDGIFEVKATAGDTHLGGEDF<br>DNRLVNHFVEEFKRKHKKDISQNKRAVRRLRTACERAKRTLSSSTQASLEIDSLFEGIDFYTSITRARFEELCSDLFR<br>STLEPVEKALRDAKLDKAQIHDLVLVGGSTRIPKVQKLLQDFFNGRDLNKSINPDEAVAYGAAVQAAILMGDKSENV<br>QDLLLLDVAPLSLGLETAGGVMTALIKRNSTIPTKQTQIFTTYSDNQPGVLIQVYEGERAMTKDNNLLGRFELSGIPP<br>APRGVPQIEVTFDIDANGILNVTATDKSTGKANKITITNDKGRLSKEEIERMVQEAEKYKAEDEVQRERVSAKNALES<br>YAFNMKSAVEDEGLKGKISEADKKKVLDKCQEVISWLDANTLAEKDEFEHKRKELEQVCNPIISGLYQGAGGPGPG<br>GFGAQGPKGGSGSGPTIEEVD |
| *mt*Dod-PAS-Pep2<br>PLEGDDDTSRMEEVD | >NP_001017963.2 heat shock protein HSP90-alpha isoform 1<br>[Homo sapiens] | MPPCSGGDGSTPPGPSLRDRDCPAQSAEYPRDRLDPRPGSPSEASSPPFLRSRAPVNWYQEKAQVFLWHLMVS<br>GSTTLLCLWKQPFHVSAFPVTASLAFRQSQGAGQHLYKDLQPFILLRLLMPEETQTQDQPMEEEEVETFAFQAEIA<br>QLMSLIINTFYSNKEIFLRELISNSSDALDKIRYESLTDPSKLDSGKELHINLIPNKQDRTLTIVDTGIGMTKADLINNLGT<br>IAKSGTKAFMEALQAGADISMIGQFGVGFYSAYLVAEKVTVITKHNDDEQYAWESSAGGSFTVRTDTGEPMGRGTK<br>VILHLKEDQTEYLEERRIKEIVKKHSQFIGYPITLFVEKERDKEVSDDEAEEKEDKEEEKEKEEKESEDKPEIEDVGSD<br>EEEEKKDGDKKKKKKIKEKYIDQEELNKTKPIWTRNPDDITNEEYGEFYKSLTNDWEDHLAVKHFSVEGQLEFRALL<br>FVPRRAPFDLFENRKKKNNIKLYVRRVFIMDNCEELIPEYLNFIRGVVDSEDLPLNISREMLQQSKILKVIRKNLVKKC<br>LELFTELAEDKENYKKFYEQFSKNIKLGIHEDSQNRKKLSELLRYYTSASGDEMVSLKDYCTRMKENQKHIYYITGET<br>KDQVANSAFVERLRKHGLEVIYMIEPIDEYCVQQLKEFEGKTLVSVTKEGLELPEDEEEKKKQEEKKTKFENLCKIMK<br>DILEKKVEKVVVSNRLVTSPCCIVTSTYGWTANMERIMKAQALRDNSTMGYMAAKKHLEINPDHSIIETLRQKAEAD<br>KNDKSVKDLVILLYETALLSSGFSLEDPQTHANRIYRMIKLGLGIDEDDPTADDTSAAVTEEMPPLEGDDDTSRMEE<br>VD |
| *mt*Dod-PAS-Pep3<br>ECYPNEKNSVNMDLD | NP_006635.2 heat shock protein 105 kDa isoform 1<br>[Homo sapiens] | MSVVGLDVGSQSCYIAVARAGGIETIANEFSDRCTPSVISFGSKNRTIGVAAKNQQITHANNTVSNFKRFHGRAFND<br>PFIQKEKENLSYDLVPLKNGGVGIKVMYMGEEHLFSVEQITAMLLTKLKETAENSLKKPVTDCVISVPSFFTDAERRS<br>VLDAAQIVGLNCLRLMNDMTAVALNYGIYKQDLPSLDEKPRIVVFVDMGHSAFQVSACAFNKGKLKVLGTAFDPFLG<br>GKNFDEKLVEHFCAEFKTKYKLDAKSKIRALLRLYQECEKLKKLMSSNSTDLPLNIECFMNDKDVSGKMNRSQFEEL<br>CAELLQKIEVPLYSLLEQTHLKVEDVSAVEIVGGATRIPAVKERIAKFFGKDISTTLNADEAVARGCALQCAILSPAFKV<br>REFSVTDAVPFPISLIWNHDSEDTEGVHEVFSRNHAAPFSKVLTFLRRGPFELEAFYSDPQGVPYPEAKIGRFVVQN<br>VSAQKDGEKSRVKVKVRVNTHGIFTISTASMVEKVPTEENEMSSEADMECLNQRPPENPDTDKNVQQDNSEAGTQ<br>PQVQTDAQQTSQSPPSPELTSEENKIPDADKANEKKVDQPPEAKKPKIKVVNVELPIEANLVWQLGKDLLNMYIETE<br>GKMIMQDKLEKERNDAKNAVEEYVYEFRDKLCGPYEKFICEQDHQNFLRLLTETEDWLYEEGEDQAKQAYVDKLE<br>ELMKIGTPVKVRFQEAEERPKMFEELGQRLQHYAKIAADFRNKDEKYNHIDESEMKKVEKSVNEVMEWMNNVMNA<br>QAKKSLDQDPVVRAQEIKTKIKELNNTCEPVVTQPKPKIESPKLERTPNGPNIDKKEEDLEDKNNFGAEPPHQNGEC<br>YPNEKNSVNMDLD |
| *mt*Dod-PAS-Pep4<br>VPSDSDKKLPEMDID | EAW62295.1 heat shock 70kDa protein 4, isoform CRA_b<br>[Homo sapiens] | MSVVGIDLGFQSCYVAVARAGGIETIANEYSDRCTPACISFGPKNRSIGAAAKSQVISNAKNTVQGFKRFHGRAFSD<br>PFVEAEKSNLAYDIVQLPTGLTGIKVTYMEEERNFTTEQVTAMLLSKLKETAESVLKKPVVDCVVSVPCFYTDAERR<br>SVMDATQIAGLNCLRLMNETTAVALAYGIYKQDLPALEEKPRNVVFVDMGHSAYQVSVCAFNRGKLKVLATAFDTT<br>LGGRKFDEVLVNHFCEEFGKKYKLDIKSKIRALLRLSQECEKLKKLMSANASDLPLSIECFMNDVDVSGTMNRGKFL<br>EMCNDLLARVEPPLRSVLEQTKLKKEDIYAVEIVGGATRIPAVKEKISKFFGKELSTTLNADEAVTRGCALQCAILSPA<br>FKVREFSITDVVPYPISLRWNSPAEEGSSDCEVFSKNHAAPFSKVLTFYRKEPFTLEAYYSSPQDLPYPDPAIEKMQ<br>VDQEEPHVEEQQQQTPAENKAESEEMETSQAGSKDKKMDQPPQAKKAKVKTSTVDLPIENQLLWQIDREMLNLYI<br>ENEGKMIMQDKLEKERNDAKNAVEEYVYEMRDKLSGEYEKFVSEDDRNSFTLKLEDTENWLYEDGEDQPKQVYV<br>DKLAELKNLGQPIKIRFQESEERPKLFEELGKQIQQYMKIISSFKNKEDQYDHLDAADMTKVEKSTNEAMEWMNNKL<br>NLQNKQSLTMDPVVKSKEIEAKIKELTSTCSPIISKPKPKVEPPKEEQKNAEQNGPVDGQGDNPGPQAAEQGTDTA<br>VPSDSDKKLPEMDID |
| *mt*Dod-PAS-Pep5<br>DSSQHTKSSGEMEVD | NP_055093.2 heat shock 70 kDa protein 4L isoform 1<br>[Homo sapiens] | MSVVGIDLGFLNCYIAVARSGGIETIANEYSDRCTPACISLGSRTRAIGNAAKSQIVTNVRNTIHGFKKLHGRSFDDPI<br>VQTERIRLPYELQKMPNGSAGVKVRYLEEERPFAIEQVTGMLLAKLKETSENALKKPVADCVISIPSFFTDAERRSV<br>MAAAQVAGLNCLRLMNETTAVALAYGIYKQDLPPLDEKPRNVVFIDMGHSAYQVLVCAFNKGKLKVLATTFDPYLG<br>GRNFDEALVDYFCDEFKTKYKINVKENSRALLRLYQECEKLKKLMSANASDLPLNIECFMNDLDVSSKMNRAQFEQ<br>LCASLLARVEPPLKAVMEQANLQREDISSIEIVGGATRIPAVKEQITKFFLKDISTTLNADEAVARGCALQCAILSPAFK<br>VREFSITDLVPYSITLRWKTSFEDGSGECEVFCKNHPAPFSKVITFHKKEPFELEAFYTNLHEVPYPDARIGSFTIQNV<br>FPQSDGDSSKVKVKVRVNIHGIFSVASASVIEKQNLEGDHSDAPMETETSFKNENKDNMDKMQVDQEEGHQKCHA<br>EHTPEEEIDHTGAKTKSAVSDKQDRLNQTLKKGKVKSIDLPIQSSLCRQLGQDLLNSYIENEGKMIMQDKLEKERND<br>AKNAVEEYVYDFRDRLGTVYEKFITPEDLSKLSAVLEDTENWLYEDGEDQPKQVYVDKLQELKKYGQPIQMKYMEH<br>EERPKALNDLGKKIQLVMKVIEAYRNKDERYDHLDPTEMEKVEKCISDAMSWLNSKMNAQNKLSLTQDPVVKVSEI<br>VAKSKELDNFCNPIIYKPKPKAEVPEDKPKANSEHNGPMDGQSGTETKSDSTKDSSQHTKSSGEMEVD |
| *mt*Dod-PAS-Pep6<br>EQSTGQKRPLKNDEL | XP_005271449.1 hypoxia up-regulated protein 1 isoform X1<br>[Homo sapiens] | MADKVRRQRPRRRVCWALVAVLLADLLALSDTLAVMSVDLGSESMKVAIVKPGVPMEIVLNKESRRKTPVIVTLKE<br>NERFFGDSAASMAIKNPKATLRYFQHLLGKQADNPHVALYQARFPEHELTFDPQRQTVHFQISSQLQFSPEEVLGM<br>VLNYSRSLAEDFAEQPIKDAVITVPVFFNQAERRAVLQAARMAGLKVLQLINDNTATALSYGVFRRKDINTTAQNIMF<br>YDMGSGSTVCTIVTYQMVKTKEAGMQPQLQIRGVGFDRTLGGLEMELRLRERLAGLFNEQRKGQRAKDVRENPR<br>AMAKLLREANRLKTVLSANADHMAQIEGLMDDVDFKAKVTRVEFEELCADLFERVPGPVQQALQSAEMSLDEIEQV<br>ILVGGATRVPRVQEVLLKAVGKEELGKNINADEAAAMGAVYQAAALSKAFKVKPFVVRDAVVYPILVEFTREVEEEP<br>GIHSLKHNKRVLFSRMGPYPQRKVITFNRYSHDFNFHINYGDLGFLGPEDLRVFGSQNLTTVKLKGVGDSFKKYPD<br>YESKGIKAHFNLDESGVLSLDRVESVFETLVEDSAEEESTLTKLGNTISSLFGGGTTPDAKENGTDTVQEEEESPAE<br>GSKDEPGEQVELKEEAEAPVEDGSQPPPPEPKGDATPEGEKATEKENGDKSEAQKPSEKAEAGPEGVAPAPEGE<br>KKQKPARKRRMVEEIGVELVVLDLPDLPEDKLAQSVQKLQDLTLRDLEKQEREKAANSLEAFIFETQDKLYQPEYQE<br>VSTEEQREEISGKLSAASTWLEDEGVGATTVMLKEKLAELRKLCQGLFFRVEERKKWPERLSALDNLLNHSSMFLK<br>GARLIPEMDQIFTEVEMTTLEKVINETWAWKNATLAEQAKLPATEKPVLLSKDIEAKMMALDREVQYLLNKAKFTKP<br>RPRPKDKNGTRAEPPLNASASDQGEKVIPPAGQTEDAEPISEPEKVETAGSEPGDTEPLELGGPGAEPEQKEQST<br>GQKRPLKNDEL |
| *mt*Dod-PAS-Pep8<br>LVTGESLEQLRRGLA | sp\|Q09118.1\|HBEGF_CHLAE | MKLLPSVVLKLLLAAVLSALVTGESLEQLRRGLAAGTSNPDPSTGSTDQLLRLGGGRDRKVRDLQEADLDLLRVTL<br>SSKPQALATPSKEEHGKRKKKGKGLGKKRDPCLRKYKDFCIHGECKYVKELRAPSCICHPGYHGERCHGLSLPVE<br>NRLYTYDHTTILAVVAVVLSSVCLLVIVGLLMFRYHRRGGYDVENEEKVKLGMTNSH |
| *mt*Dod-PAS-Pep9<br>MKGKEEKEGGARLGA | NP_005852.2 E3 ubiquitin-protein ligase CHIP isoform a<br>[Homo sapiens] | MKGKEEKEGGARLGAGGGSPEKSPSAQELKEQGNRLFVGRKYPEAAACYGRAITRNPLVAVYYTNRALCYLKMQ<br>QHEQALADCRRALELDGQSVKAHFFLGQCQLEMESYDEAIANLQRAYSLAKEQRLNFGDDIPSALRIAKKKRWNSI<br>EERRIHQESELHSYLSRLIAAERERELEECQRNHEGDEDDSHVRAQQACIEAKHDKYMADMDELFSQVDEKRKKR<br>DIPDYLCGKISFELMREPCITPSGITYDRKDIEEHLQRVGHFDPVTRSPLTQEQLIPNLAMKEVIDAFISENGWVEDY |
| *mt*Dod-PAS-Pep10<br>EERRIHQESE | NP_005852.2 E3 ubiquitin-protein ligase CHIP isoform a<br>[Homo sapiens]<br>Pos. 151-160 | MKGKEEKEGGARLGAGGGSPEKSPSAQELKEQGNRLFVGRKYPEAAACYGRAITRNPLVAVYYTNRALCYLKMQ<br>QHEQALADCRRALELDGQSVKAHFFLGQCQLEMESYDEAIANLQRAYSLAKEQRLNFGDDIPSALRIAKKKRWNSI<br>EERRIHQESELHSYLSRLIAAERERELEECQRNHEGDEDDSHVRAQQACIEAKHDKYMADMDELFSQVDEKRKKR<br>DIPDYLCGKISFELMREPCITPSGITYDRKDIEEHLQRVGHFDPVTRSPLTQEQLIPNLAMKEVIDAFISENGWVEDY |
| *mt*Dod-PAS-Pep11<br>NHEGDEDDSH | NP_005852.2 E3 ubiquitin-protein ligase CHIP isoform a<br>[Homo sapiens]<br>Pos. 183-192 | MKGKEEKEGGARLGAGGGSPEKSPSAQELKEQGNRLFVGRKYPEAAACYGRAITRNPLVAVYYTNRALCYLKMQ<br>QHEQALADCRRALELDGQSVKAHFFLGQCQLEMESYDEAIANLQRAYSLAKEQRLNFGDDIPSALRIAKKKRWNSI<br>EERRIHQESELHSYLSRLIAAERERELEECQRNHEGDEDDSHVRAQQACIEAKHDKYMADMDELFSQVDEKRKKR<br>DIPDYLCGKISFELMREPCITPSGITYDRKDIEEHLQRVGHFDPVTRSPLTQEQLIPNLAMKEVIDAFISENGWVEDY |

**Literature:**


1. Golovanov, A. P., Hautbergue, G. M., Wilson, S. A. & Lian, L.-Y. A Simple Method for Improving Protein Solubility and Long-Term Stability. *J. Am. Chem. Soc.* **126**, 8933–8939 (2004).
2. Bajar, B. T. *et al.* Improving brightness and photostability of green and red fluorescent proteins for live cell imaging and FRET reporting. *Sci. Rep.* **6**, (2016).
3. Veggiani, G. *et al.* Programmable polyproteams built using twin peptide superglues. *Proc. Natl. Acad. Sci.* **113**, 1202–1207 (2016).
4. Kick, A., Bönsch, M. & Mertig, M. EGNAS: an exhaustive DNA sequence design algorithm. *BMC Bioinformatics* **13**, 138 (2012).
